# Prioritizing virtual screening with interpretable interaction fingerprints

**DOI:** 10.1101/2022.05.25.493419

**Authors:** Alexandre V. Fassio, Laura Shub, Luca Ponzoni, Jessica McKinley, Matthew J. O’Meara, Rafaela S. Ferreira, Michael J. Keiser, Raquel C. de Melo Minardi

**Affiliations:** São Carlos Institute of Physics, University of São Paulo, São Carlos, São Paulo, Brazil; Department of Biochemistry and Immunology, Federal University of Minas Gerais, Belo Horizonte, Minas Gerais, Brazil; Department of Pharmaceutical Chemistry, Department of Bioengineering & Therapeutic Sciences, Institute for Neurodegenerative Diseases, Kavli Institute for Fundamental Neuroscience, Bakar Computational Health Sciences Institute, University of California, San Francisco, San Francisco, California, United States; Department of Computational Medicine and Bioinformatics, University of Michigan, Ann Arbor, Michigan, United States; Department of Computer Science, Federal University of Minas Gerais, Belo Horizonte, Minas Gerais, Brazil

## Abstract

Machine learning-based drug discovery success depends on molecular representation. Yet traditional molecular fingerprints omit both the protein and pointers back to structural information that would enable better model interpretability. Therefore, we propose LUNA, a Python 3 toolkit that calculates and encodes protein-ligand interactions into new hashed fingerprints inspired by Extended Connectivity FingerPrint (ECFP): EIFP (Extended Interaction FingerPrint), FIFP (Functional Interaction FingerPrint), and Hybrid Interaction FingerPrint (HIFP). LUNA also provides visual strategies to make the fingerprints interpretable. We performed three major experiments exploring the fingerprints’ use. First, we trained machine learning models to reproduce DOCK3.7 scores using 1 million docked Dopamine D4 complexes. We found that *EIFP-4,096* performed (*R*^2^ = 0.61) superior to related molecular and interaction fingerprints. Secondly, we used LUNA to support interpretable machine learning models. Finally, we demonstrate that interaction fingerprints can accurately identify similarities across molecular complexes that other fingerprints overlook. Hence, we envision LUNA and its interface fingerprints as promising methods for machine learning-based virtual screening campaigns. LUNA is freely available at https://github.com/keiserlab/LUNA.

## Introduction

In the last few decades, machine learning (ML), and in particular deep neural networks (DNN), have opened up new opportunities in all stages of drug discovery and development.^1–19^ In particular, ML has the potential to revolutionize structural-based virtual screening (SBVS) campaigns through the prioritization and automatic selection of promising compounds. SBVS nowadays^20–23^ commonly involves a manual and intensive process called hit-picking, in which researchers evaluate and determine hit selection through molecular graphics programs to inspect the binding modes of hundreds of top-scoring compounds. This process is time-intensive and highly dependent on the researcher’s expertise. Thus, automating hit selection would bring great benefit to the drug discovery community.

However, the success of any ML-based drug discovery endeavor heavily relies on data representation.^24–26^ Not surprisingly, several representations have been proposed since the early days of chemoinformatics. One of the most popular manners to represent molecules or protein-ligand complexes is the so-called “fingerprint” (FP), a bit vector in which each bit accounts for the presence (1s) or absence (0s) of a given chemical feature or substructure.^27^ Alternatively, bits can be substituted by the explicit frequency (count) of a given chemical feature or substructure.

FPs are of two major types – structural or hashed – and can be sub-classified into molecular (MFP) and interaction (IFP) FPs according to the information they encode. In structural FPs, each bit in a binary sequence encodes a predefined molecular^28–33^ or protein-ligand interaction (PLI)^34–53^ substructure. In contrast, hashed FPs do not rely on a predefined list of substructures, but instead explicitly identify and enumerate all substructures in a given data set of molecules^54–59^ or protein-ligand complexes,^60^ which are then mapped to a bit position through a folding operation given by a hash function. The major advantage of hashed FPs is the generality as they do not require defining a set of substructures *a priori*. However, the use of hash functions may produce information loss due to bit collisions, which occur when two different substructures are encoded at the same bit position.

Recently, learned representations^6, 61, 62^ have also become popular with advancements in DNN techniques such as convolutional neural networks,^8, 63, 64^ graph convolutional networks,^65–67^ variational autoencoders,^68^ and deep metric learning.^69^ In summary, these approaches transform raw input features into a continuous learned representation, which means that the data representation is learned directly from the training data. The advantage of learned representations over traditional FPs is that they are unique and do not suffer from information loss due to bit collisions. However, they require considerably more data to achieve generalization and learn even simple patterns, ^25^ and have yet to prove their superiority consistently over traditional FPs. ^70, 71^

In this paper, we present LUNA^1^, an object-oriented Python 3 toolkit for drug design that makes it easy to analyze very large data sets of 3D molecular structures and complexes, and that allows identifying, filtering, and visualizing atomic interactions. LUNA also implements three novel hashed IFPs: Extended Interaction FingerPrint (EIFP), Functional Interaction FingerPrint (FIFP), and Hybrid Interaction FingerPrint (HIFP). These FPs encode molecular interactions at different levels of detail and are fully interpretable, providing several functionalities to trace individual bits back to their original atomic substructures in the context of the binding site.

To validate and illustrate the applicability of the IFPs, we first present a case study where we trained several ML models on a data set containing 1 million molecules docked against Dopamine D4^72^ to reproduce their DOCK3.7^73^ scores as in a re-scoring task. We then compare our results to four related FPs: Extended-Connectivity FingerPrint (ECFP), ^58^ Functional-Class FingerPrint (FCFP),^58^ Extended Three-Dimensional FingerPrint (E3FP), ^59^ and Protein–Ligand Extended Connectivity (PLEC). ^60^ Secondly, we demonstrate how key interactions emerge from looking at FP features with an interpretability study. Finally, we present two experiments where we evaluate the capacity of the novel FPs to distinguish ligands by pose similarity and to assess how different feature representations impacted these analyses. To do so, we built five protein-ligand complex data sets with varied diversity and complexity, and compared the results to ECFP, FCFP, E3FP, and PLEC.

## Results

### Novel hashed interaction fingerprints

In this work, we propose three novel hashed IFPs inspired by ECFP,^58^ FCFP,^58^ and E3FP.^59^ ECFP and FCFP are 2D hashed MFPs that encode molecular graph connectivity using either explicit atomic substructures or “coarse-grained” pharmacophoric properties, respectively. E3FP, another hashed MFP, attempts to circumvent ECFP’s and FCFP’s lack of explicit 3D molecular structure patterns. However, MFPs do not explicitly encode protein-interface information. PLEC,^60^ a hashed IFP also motivated by ECFP, encodes interface information, but only in the form of proximal protein-ligand interactions, without interaction typing. It also omits pointers back to structural information that would enable better model interpretability. Thus, we develop the new IFPs to extend prior approaches to nonbonded interaction patterns at the ligand-protein interface and call them EIFP, FIFP, and HIFP.

While EIFP and FIFP apply the logic of ECFP and FCFP, respectively, HIFP adopts a “hybrid” approach, encoding pharmacophoric properties for atom groups and precise environment information for atoms. All these IFPs are RDKit^74^-compatible and their features represent interactions and contacts between protein residues, ligand atoms, and water molecules, by detecting their presence or absence (bit FPs), or their frequency (count FPs). The IFPs rely on three parameters: the FP length, the radius growth rate, and the number of levels (Figure 1). While the FP length determines the maximum number of features that can be individually represented in it, the radius growth rate and the number of levels control the number of features generated. In Section *Interaction fingerprints*, we discuss how these FPs are generated and how these parameters influence the featurization process.

**Figure 1:**
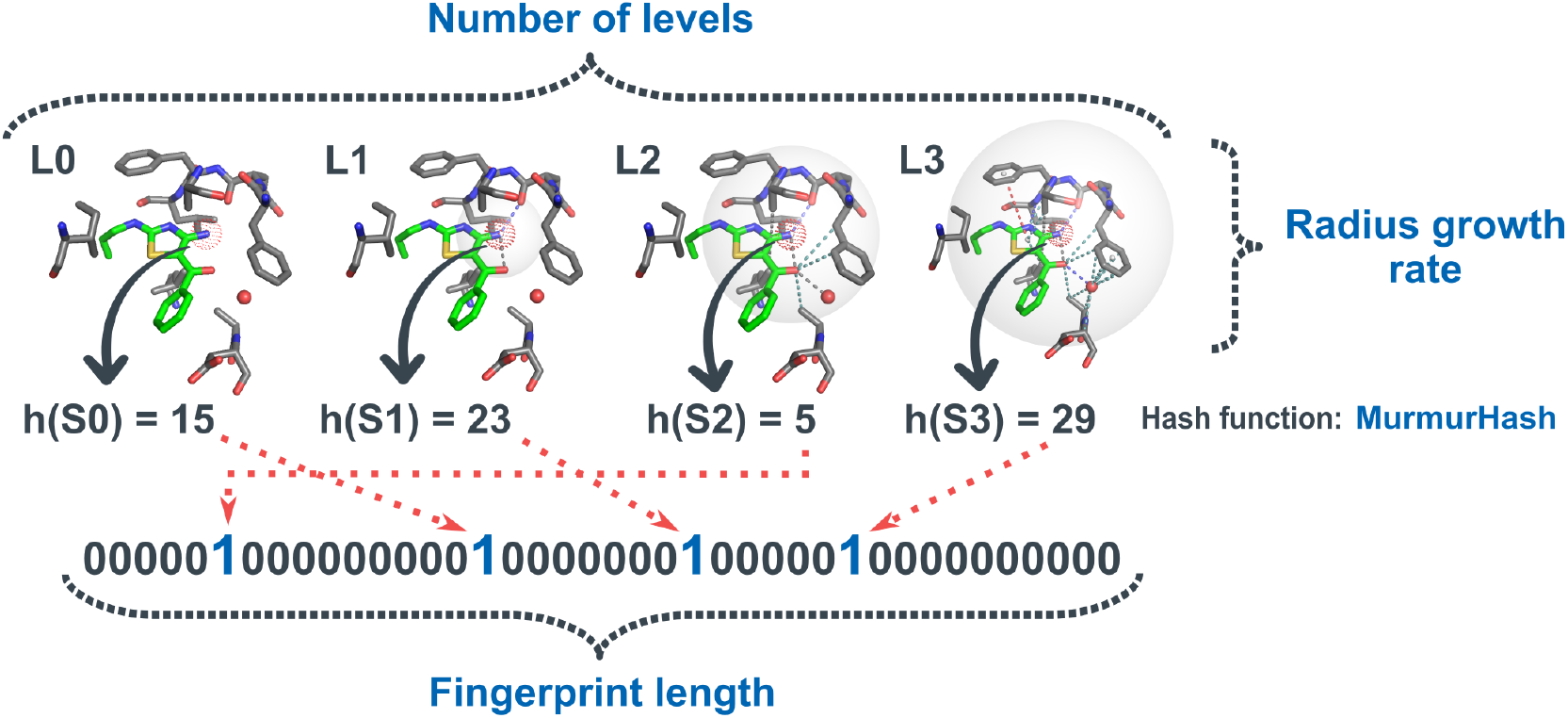
Parameters to control the IFPs creation: the FP length, the radius growth rate, and the number of levels

The new IFPs are provided as part of LUNA, an object-oriented Python 3 toolkit for drug discovery. In addition, LUNA implements several features geared towards the analysis of molecular complexes, such as: *a*) accepting any molecular complex type, including protein-ligand and protein-protein; *b*) accepting multiple file formats, including PDB, MOL, and MOL2; *c*) providing pre- and post-processing functions to control how interactions are calculated or selected based on geometric constraints; and *d*) providing several functions to summarize, characterize, and visualize molecular interactions in Pymol (Figure 2).

**Figure 2:**
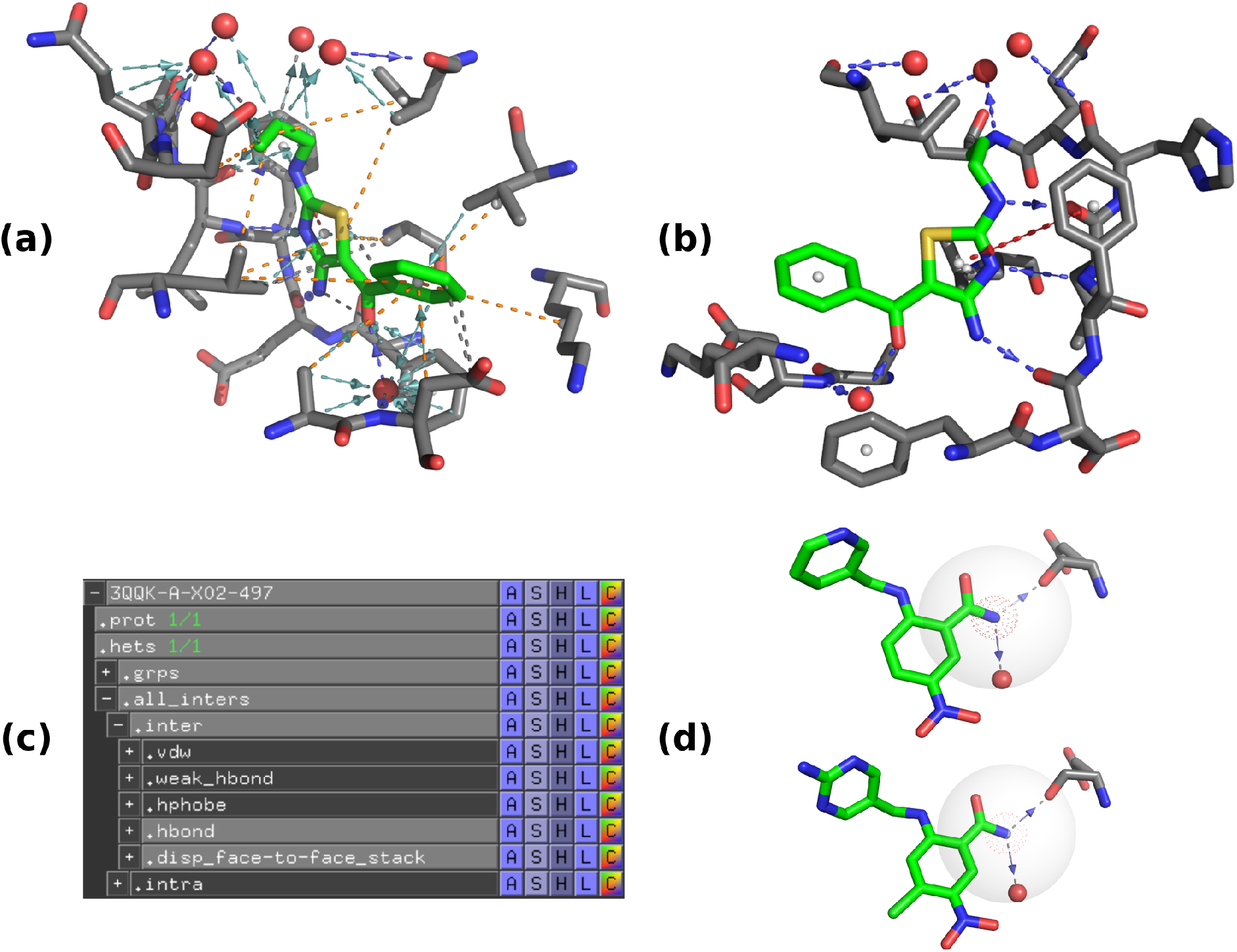
Examples of Pymol sessions automatically generated by LUNA. (a) An overview of all interactions found in a CDK2 complex (PDB 3QQK). (b) The same view shown in (a) but rotated and filtered for *hydrogen bonds* (blue) and a *displaced face-to-face stacking* (red), which can be easily achieved directly through Pymol as shown in (c). (d) Two similar bits (neighborhoods) found in the FPs of two other CDK2-inhibitor complexes (PDBs 3QQF and 3QWJ, top and bottom). Arrows represent directional interactions.

Most importantly, LUNA provides powerful functionalities to make the IFPs interpretable. One drawback of hashed FPs is that the structural information encoded in a bit is usually discarded after hashing. On top of that, due to FP folding, multiple substructures may end up encoded in the same bit, the so-called collision problem. To address this problem, LUNA keeps a record of the original features hashed into a new bit position. Thus, users can recover and reconstruct the neighborhood (collection of atoms or atom groups and their interactions) that generated a particular bit, and they can select one or more recovered neighborhoods to perform statistical analysis or export them to a rich Pymol session, where they will be able to visually assess the information present in a bit position (Figure 2d).

In the following sections, we will explore the potential of LUNA and the novel IFPs through a ML case study, an interpretability study, and demonstrate how they can be applied for detecting similar binding modes, an essential task in virtual screening.

### Interaction fingerprints improve DOCK3.7 score regression

To assess the use of the IFPs for prioritizing hit compounds, we trained DNN models using a large data set recently published by Lyu et al..^72^ This library consists of *∼*96 million molecules docked against the Dopamine D4 receptor (D4R), a G protein-coupled receptor (GPCR) superfamily member involved in many different roles in the central nervous system. Given its relevance as a neurotransmitter, dysregulation of the Dopamine D4 signalling cascade is linked to several pathological disorders including Parkinson’s disease and schizophrenia.^75^

Departing from Lyu et al.’s work, we built a random diverse and stratified data set based on DOCK3.7 score bins in order to achieve model generalizability through chemical diversity and representativeness across the DOCK3.7 score distribution. In total, we obtained 1 million molecules, from which a random 90% and 10% were set apart for model development and held-out testing, respectively.

As baselines, we chose the RDKit implementations of ECFP and FCFP. We also chose E3FP and PLEC for further comparison. FP parameters remained as default, except for the cases described in Section *Model development and training*. Additionally, by default, PLEC generated FPs of length 16,384. However, we hypothesized that a length of 4,096 might lead to results matching its longer version. To test this hypothesis, we optimized DNN parameters for the smaller PLEC version as well. To specify length, we will refer to a FP using its name followed by a hyphen and its length. For instance, *PLEC-4,096* and *EIFP-4,096* refer to PLEC and EIFP of length 4,096, respectively.

First, we performed a hyperparameter search on LUNA’s and the IFP parameters (Table S1). Given the large parameter space to cover, we defined a separate and smaller training subsample of size 200,000 using the same strategy we described for the 1 million molecules data set, used exclusively for this experiment. From these, 20% was set apart for model validation. In all experiments, we used the best architecture found in previous investigations (results not shown) and, in total, 294 parameter combinations were evaluated (Figures S1 and S2; Excel file in the Supporting Information). Overall, we observed that FPs with fewer levels, high radius growth rate, precise atomic topology definition (EIFP), non-covalent interactions, and count information produced the best models (Section *Interaction fingerprints* details how these parameters influence the featurization process). In Table S2, we show the parameters for the best IFP (*EIFP-4,096*; *R*^2^ = 0.52) selected for subsequent analysis.

The DNN hyperparameter search was employed on the space shown in Table S3. We trained a single deterministic model (seed 54,343) for each sampled parameter using 80% of the training set and evaluating it with the remaining 20%. Moreover, to test the hypothesis that an IFP of length 4,096 might be sufficient, we also evaluated *EIFP-16,384*. In total, 165 and 30 different parameter configurations were evaluated for FPs of length 4,096 and 16,384, respectively (Figures S3-S9). Results for the best models are shown in Figures 3, S10-S15, and Table 1. These results showed that relatively simple model designs performed best at predicting the DOCK3.7 scores. Also, we found that *EIFP-4,096* achieved a performance as good as that obtained for the longer FPs and superior to other FPs with the same length. The complete list of parameters covered for each FP and resulting *R*^2^ values are available as an Excel file in the Supporting Information.

**Figure 3:**
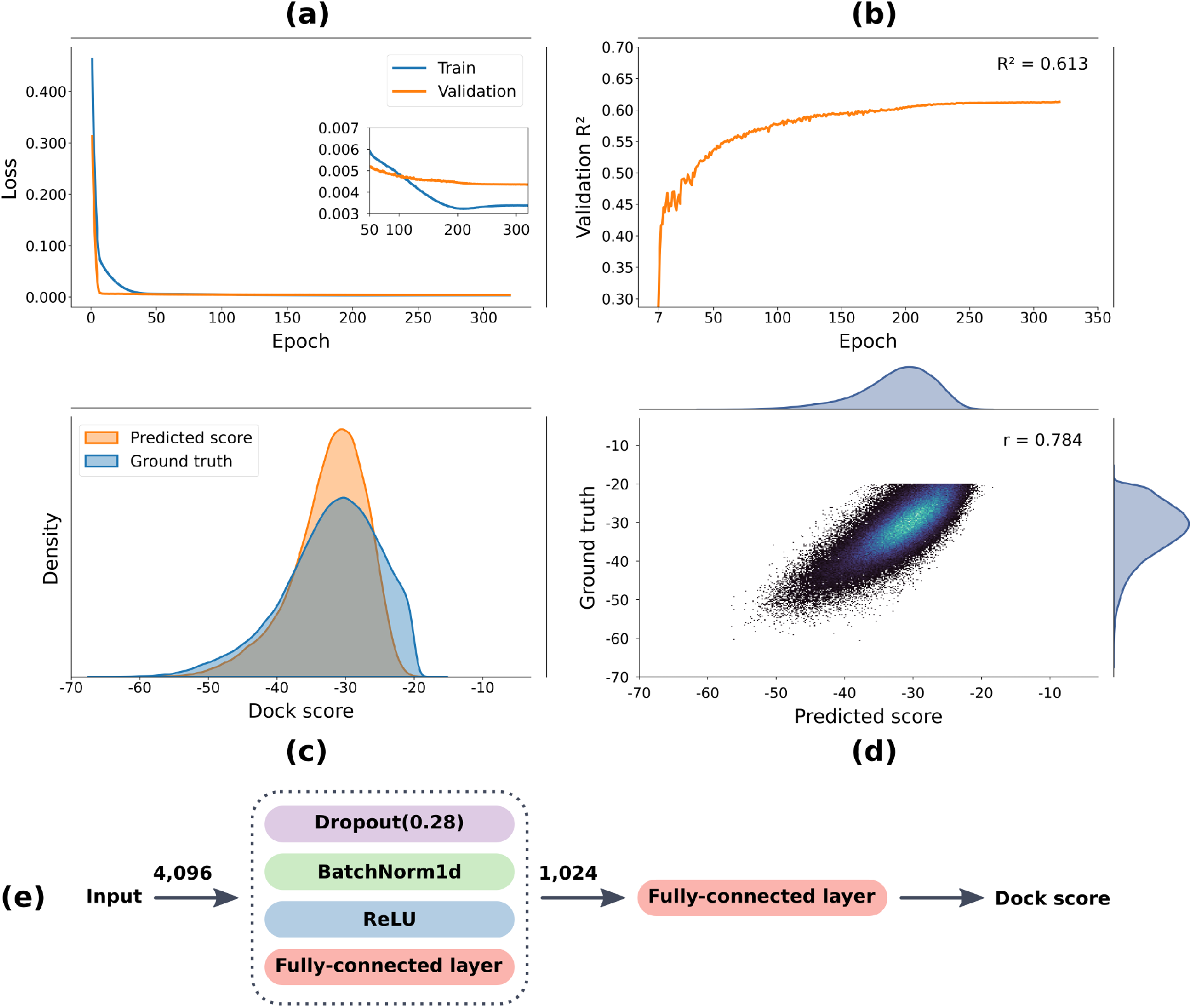
Performance of the best DNN model trained with *EIFP-4,096*. (a) The loss curve for the training and validation sets across the epochs. (b) The validation *R*^2^ across the epochs. (c) The predicted and ground truth score distributions. (d) A scatter plot for the ground truth versus the predicted scores, as well as the Pearson correlation coefficient (*r*) between them. (e) The best architecture found for *EIFP-4,096*.

**Table 1:**
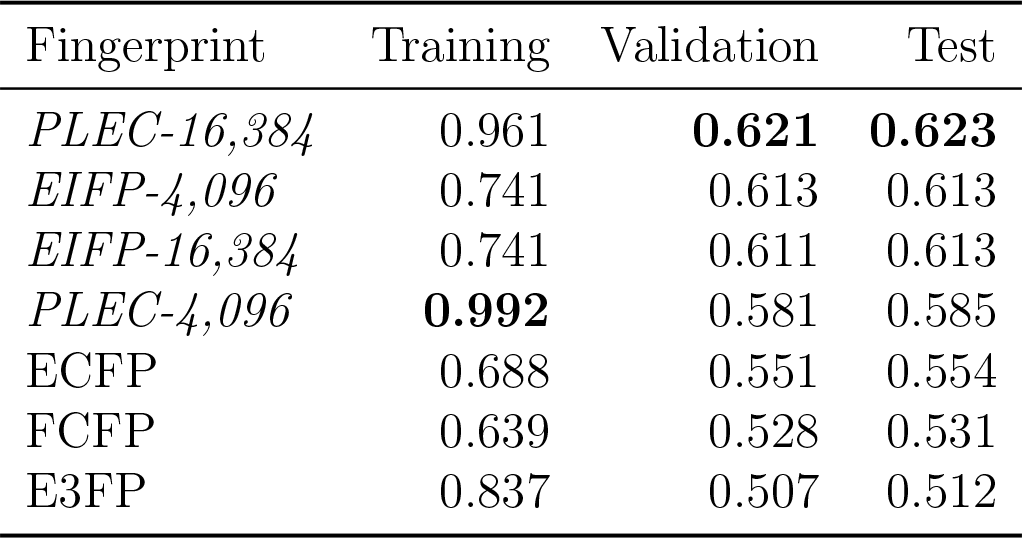
Training, validation, and test set *R*^2^ for the best DNN model of each FP.

| Fingerprint | Training | Validation | Test |
| --- | --- | --- | --- |
| *PLEC-16,384* | 0.961 | **0.621** | **0.623** |
| *EIFP-4,096* | 0.741 | 0.613 | 0.613 |
| *EIFP-16,384* | 0.741 | 0.611 | 0.613 |
| *PLEC-4,096* | **0.992** | 0.581 | 0.585 |
| ECFP | 0.688 | 0.551 | 0.554 |
| FCFP | 0.639 | 0.528 | 0.531 |
| E3FP | 0.837 | 0.507 | 0.512 |

Finally, we applied the resulting DNN models to the test set, which consisted of 100,000 unseen protein-ligand complexes (Table 1). Overall, *PLEC-16,384*, *EIFP-4,096*, and *EIFP-16,384* were the best FPs, followed by *PLEC-4,096*. 5-fold cross-validation showed similar results and demonstrated the differences between models were significant (Tables S4-S6 and Figure S16), which again highlights the best trade-off between FP length and performance of *EIFP-4,096*.

We observed that the validation and test set performances were similar. Considering that the diverse cluster heads (molecules) composing the sets used for model development (training and validation sets) and testing were mutually exclusive, these results suggested we succeeded at constructing balanced data sets by guaranteeing chemical diversity and representativeness across the DOCK3.7 score distribution. To test this hypothesis, we conducted an experiment of drastically hampering the model’s expected generalization ability by training a version of the model on randomly increasingly smaller fractions of the development data set. To do so, we performed seven cuts that progressively removed 50% of the remaining data until reaching a minimum development data set of size 7,032 poses, of which 20% was always used for validation. At each cut, a new single model was trained for each FP and the performance of the resulting model was measured with the full test set. This experiment was performed for both *EIFP-4,096* and *EIFP-16,384*.

As the training and validation sets may share common cluster heads, we expected the model performance for the validation set to be higher than that of the test set when the model was trained with fewer data examples. However, the *R*^2^ obtained for *EIFP-4,096* across the different cuts showed that the performance of validation and test sets reduced proportionally and were still similar even after the cuts on the training set exceeding two orders of magnitude (Figure 4). These findings confirmed our hypothesis that we obtained considerably generalized models due to the chemical structural diversity and representation across the DOCK3.7 score distribution.

**Figure 4:**
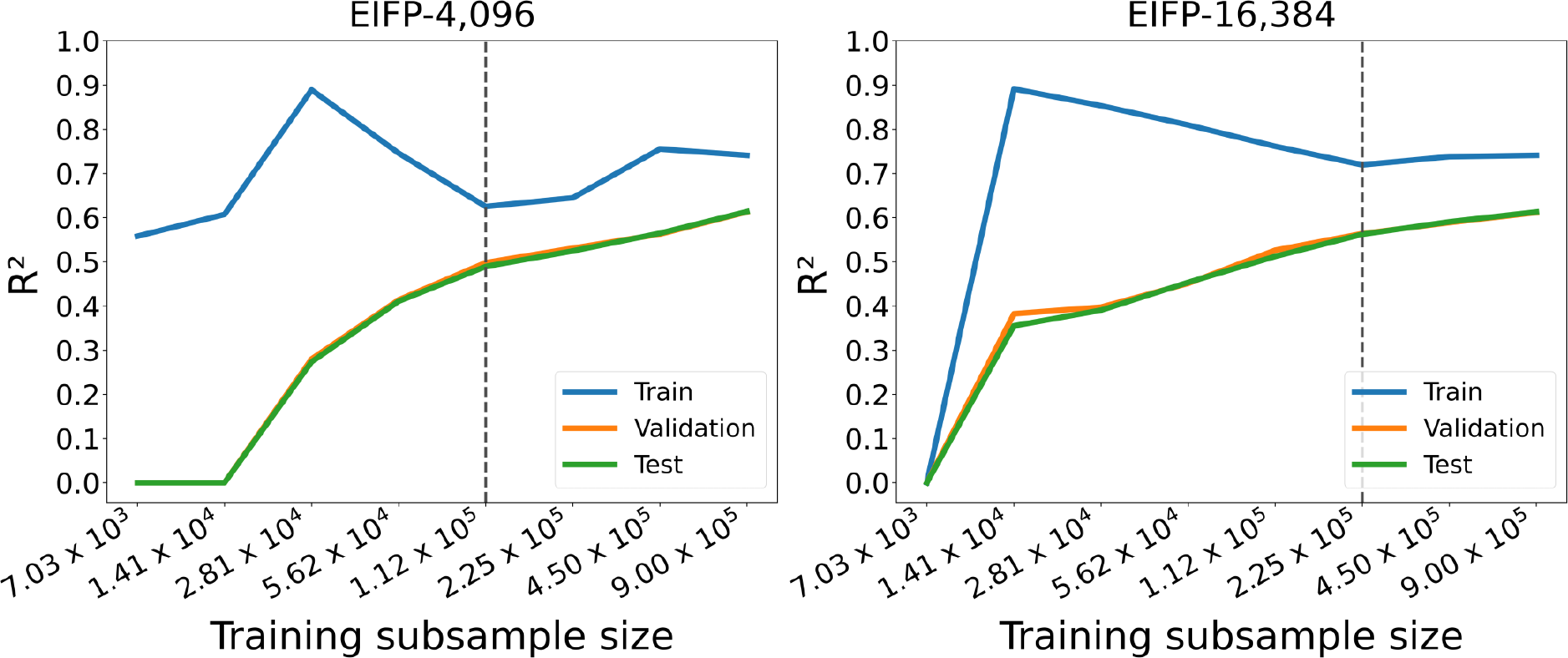
Performance of the best DNN architecture trained with *EIFP-4,096* and *EIFP-16,384* across different training set sizes.

Additionally, we observed that from subsample size 112,500 downwards, the models started overfitting. Also, the subsample 112,500 could be considered a good starting point data set size given its reasonable performance. Notably, the slope on the right-side of Figure 4 also indicated that the model could further improve if more data points were provided as input.

Larger IFPs (Figure 4) showed similar behavior. However, the overfitting problem started earlier (after the subsample 225,000), indicating that larger FPs are more of a risk than a benefit for small data sets.

### Key interactions emerge from fingerprint features

To illustrate how LUNA can be used to support ML model interpretability, we performed a feature attribution analysis using the XGBoost library, whereby each feature received a score based on its contribution to the model’s predictions. We favored XGBoost models (performance reported in Table S5) rather than the DNNs for this analysis due to the comparative ease of attribution interpretability. The more complicated attribution techniques for DNNs such as a integrated gradients^76^ rely on defining a baseline input hyperparameter,^77, 78^ the ideal choice of which is less obvious for MFPs.

We trained the XGBoost model using the best EIFP identified in previous analysis and 80% of the training set, while the remaining 20% was used for model validation. To facilitate the interpretation, the 4,096 features were grouped into classes, according to whether they encode: *ligand’s level 0 features only*; *protein’s level 0 features only*; *upper level with ligand atomic information only*; *upper level with protein atomic information only*; *intra-ligand interactions only*; *intra-protein interactions only*; *has non-covalent interactions with the protein*; and *features with collision in the same complex*. An example of each class is shown in Figure 5.

**Figure 5:**
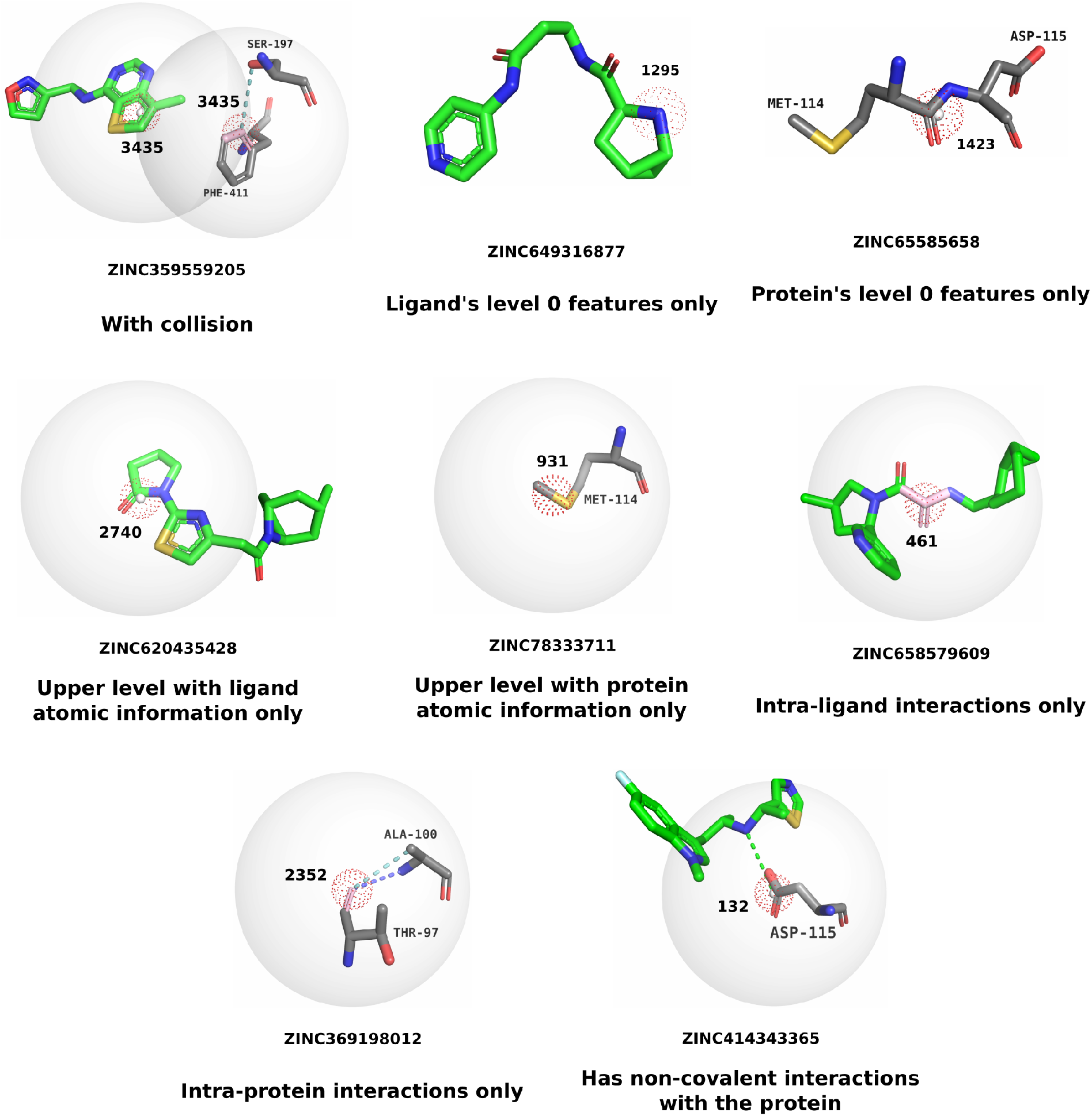
Examples of classes attributed to *EIFP-4,096* features. Ionic, hydrogen bonds, and weak hydrogen bonds are depicted as green, blue, and teal dashed lines, respectively. Features at level 0 contain only atomic information and are represented by dotted red spheres. Features at higher levels may contain interactions and atomic information and are represented by solid spheres, whereas dotted red spheres are their centroids. Covalent bonds comprising the feature are depicted as pink sticks.

Due to bit collision resulting from FP folding, some features may be assigned to multiple classes. Thus, whenever it was not possible to assign a predominant class (threshold of 69.49%, chosen by z-scores), a feature was classified as ‘unreliable’ (see Section *Model interpretability* for more details).

Analysis of the feature importance scores revealed that a small number of features comprised the major contribution to model performance. To identify them, we calculated z-scores for each feature using their importance scores, which were then converted into p-values (see Section *Model interpretability* for more details). Most important features were then defined as those having p-values less than 0.01. In total, 38 features matched this requirement.

Among the most important features’ classes, 34% (13) comprised shells containing PLIs, 29% (11) represented ligand atomic information, and the remaining classes sum to 37% (11) (Figure S17). Another three features were considered unreliable as their most prevalent class had percentages smaller than the minimum value (69.49%) (Figure S18). Interestingly, when accounting for all features regardless of importance, those containing PLIs were the rarest. PLI features were more frequent among the most potent ligands (Figures S19 and S20).

We then assessed which interactions were encoded in the thirteen most important features comprising PLIs. As multiple interactions were assigned to the same feature (Figure S21), we calculated z-scores with the percentage of the most prevalent interaction in each feature. Then, a threshold to identify the most prevalent interaction in a given feature was defined as the minimum percentage (82.78%) among features presenting z-scores higher than 1. Two features presented percentages below the threshold and were classified as *unreliable feature*. Prevalent interacting residues were also identified with the same threshold. We summarized the number of ligands having the most prevalent interactions and identify which residue established them in Figure 6a. An example of each feature is shown in Figures 6b and S22.

**Figure 6:**
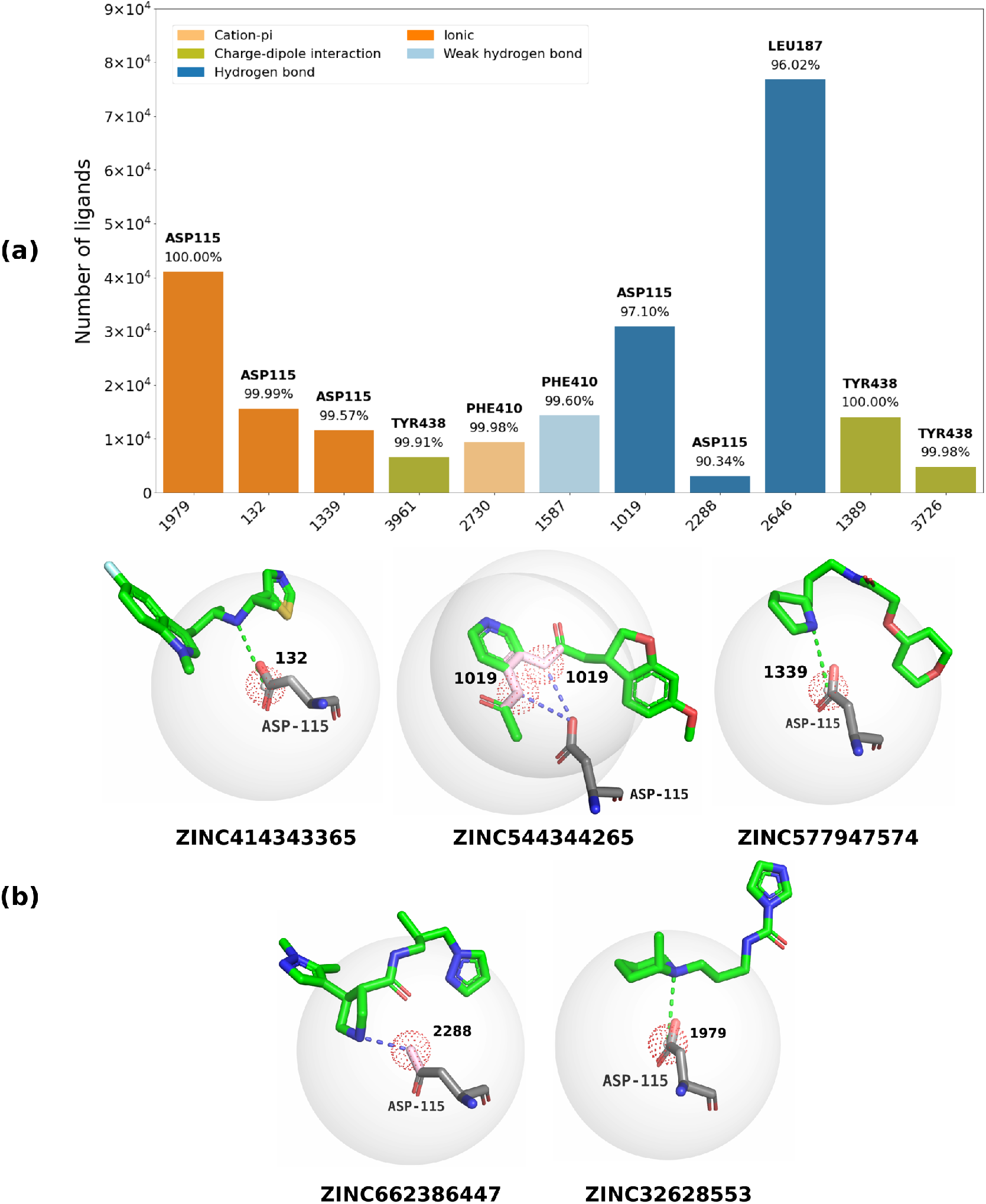
(a) Number of ligands that contain the most important features (identified by FP vector index, x-axis) encoding protein-ligand interactions. The residues that established the most common interaction in a given feature and its frequency are shown above the bars. (b) Examples of important features comprising interactions with the key residue ASP115. Ionic and hydrogen bonds are depicted as green and blue dashed lines, respectively. The feature is represented by the bigger sphere, while the dotted red sphere highlights its centroid. Covalent bonds comprising the feature are depicted as pink sticks. Features are sorted in descending order of the importance score.

After identifying the prevalent interactions, we observed that the three best features comprised ionic interactions with ASP115 and two others formed hydrogen bonds with that same residue (Figure 6b). Strikingly, ASP115 is a key residue present in all aminergic receptors, a subfamily of GPCRs that include the dopamine receptors.^72, 79–81^ Further analysis also revealed that interactions encoded at the features *1979*, *132*, and *1339* were established by the ligand’s nitrogen atoms, which also emerged as highly important and whose information were encoded in features *2*, *530*, and *1295*, respectively (Figure S19). Moreover, we also observed that these ionic interactions tended to occur primarily in the most potent ligands (Figure 7).

**Figure 7:**
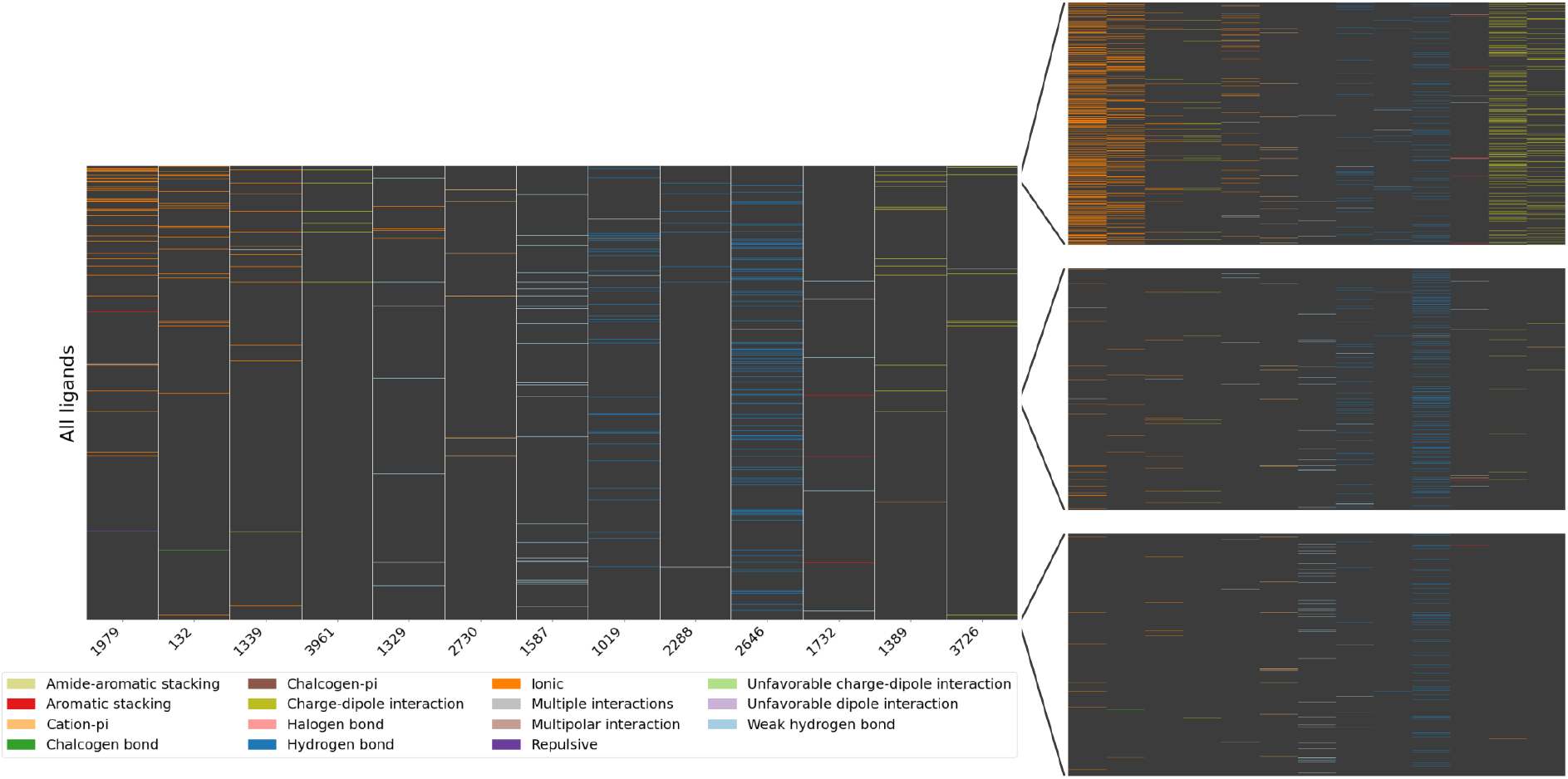
Heat map for the thirteen most important features comprising protein-ligand interactions across all ligands in the training set. Zoomed-in heat maps highlight the presence of an interaction among the best, median, and worst ligands. Features are sorted in descending order of the importance score.

### Interaction fingerprints enhance their 2D counterparts

To validate the proposed IFPs and assess how different molecular and interaction information impact the identification of similar complexes, we compared the IFPs to molecular (ECFP, FCFP, and E3FP) and interaction (PLEC) FPs. Given the close relationship between molecular structure and interactions, we expected to find a high correlation between these FPs regarding their similarities. We also generated IFPs with restrictions on the encoded information as a means to simulate ECFP, FCFP, E3FP, and PLEC using the LUNA toolkit, for further comparison. *ECFP-like EIFP*, for instance, contained only intra-ligand covalent interactions, as in classical ECFP. The complete list of features for each IFP is presented in Table 2. All FPs were generated considering a length of 4,096 bits and without encoding counts.

**Table 2:**
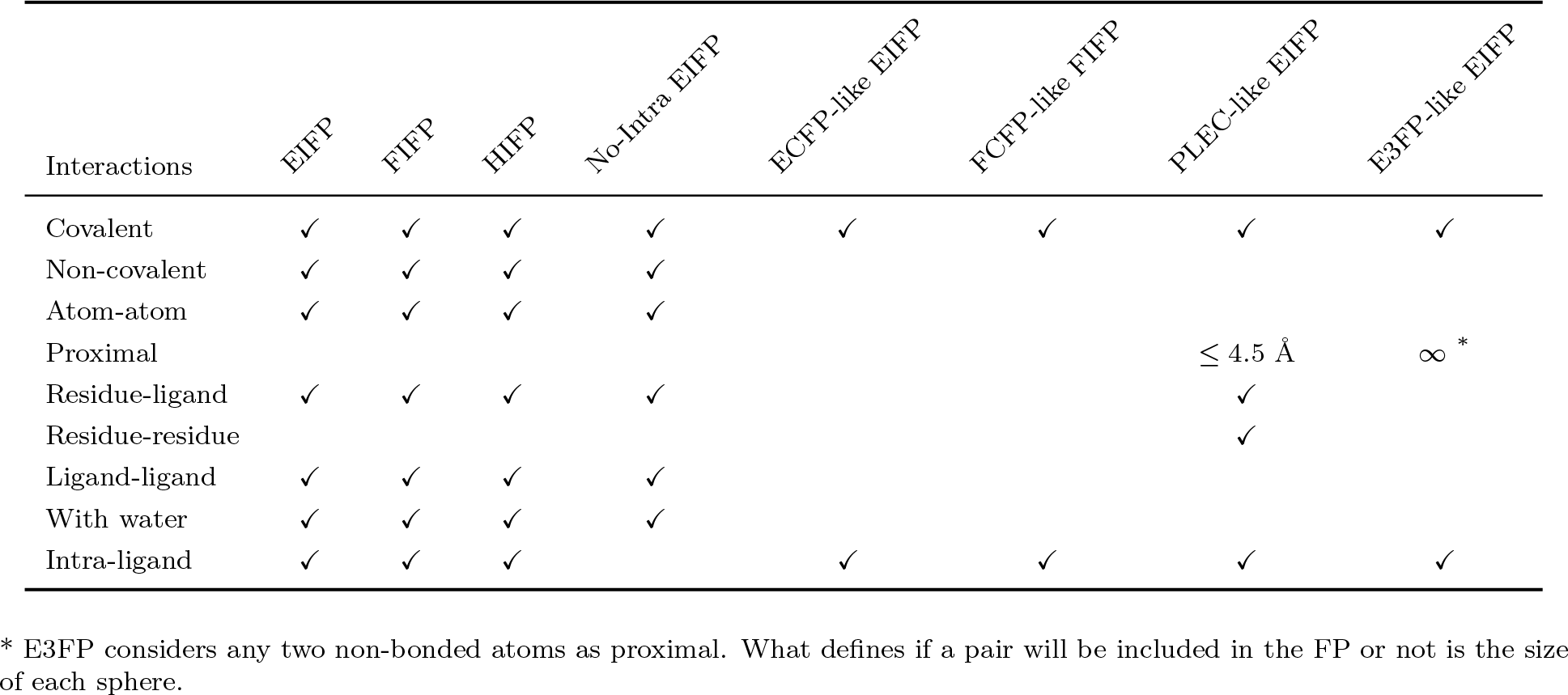
Parameters employed to construct each version of the IFPs.

In this analysis, we used five different data sets, progressively increasing protein and ligand variability, in order to evaluate how well FP similarity alone could discriminate similar poses as feature complexity progressed. The first data set contained two series of conformers for the ligand X02 in complex with the human cyclin-dependent kinase 2 (CDK2; PDB id 3QQK). The maximum conformers’ distance to the crystal pose at each series were 0.4 Å and 0.8 Å, respectively. The second data set contained 74 CDK2-ligand complexes solved through X-ray crystallography and obtained from Schonbrunn et al.. ^82^ The third data set contained all CDK2 complexes found in the RCSB PDB (350 PDB ids in total). The fourth data set comprised non-CDK2 kinase complexes obtained from RCSB PDB (1,364 PDB ids in total). Finally, the fifth data set were also obtained from RCSB PDB and covered randomly selected diverse classes of proteins except for kinases (37,507 PDB ids in total). Fourth and fifth data sets were balanced to avoid bias towards a receptor and only crystal structures with X-ray resolution below 2 Å were retained. All complexes were recovered from RCSB PDB in May 2020. In Table 3, we presented each data set and the mean and standard deviation (std) for proteins’ and ligands’ similarities.

**Table 3:** Data set summary.

| # | Data set | # Complexes | # Unique | | TM-score | | $T_C$ | |
| --- | --- | --- | --- | --- | --- | --- | --- | --- |
|  |  |  | Proteins | Ligands | Mean | Std | Mean | Std |
| 1a | Automatically generated conformers (0.4 Å) | 53 | 1 | 1 | 1.00 | 0.00 | 1.00 | 0.00 |
| 1b | Automatically generated conformers (0.8 Å) | 431 | 1 | 1 | 1.00 | 0.00 | 1.00 | 0.00 |
| 2 | CDK2 complexes from Schonbrunn’s work<sup>82</sup> | 74 | 1 | 73 | 0.98 | 0.01 | 0.27 | 0.19 |
| 3 | Extended CDK2 set | 337 | 1 | 281 | 0.91 | 0.13 | 0.12 | 0.10 |
| 4 | Other non-CDK2 kinases | 685 | 143 | 399 | 0.65 | 0.20 | 0.10 | 0.13 |
| 5 | Diverse proteins | 550 | 111 | 183 | 0.30 | 0.13 | 0.11 | 0.15 |

TM-score<sup>83</sup> and $T_C$ measure proteins and ligands similarities, respectively.

After generating FPs for complexes from data sets 1-5, we calculated the Tanimoto (*T_C_*) similarity between pairs of FPs of a given type and used these similarities to calculate the Pearson correlation coefficient between different FPs overall (Figure 8a). Full similarity distributions are available in Figures S23-S28. For data set 1, we also calculated the root mean square deviation (RMSD) between the ligands and measured the correlation of the obtained RMSDs to the FP similarities (Figures 8b, S29, and S30).

**Figure 8:**
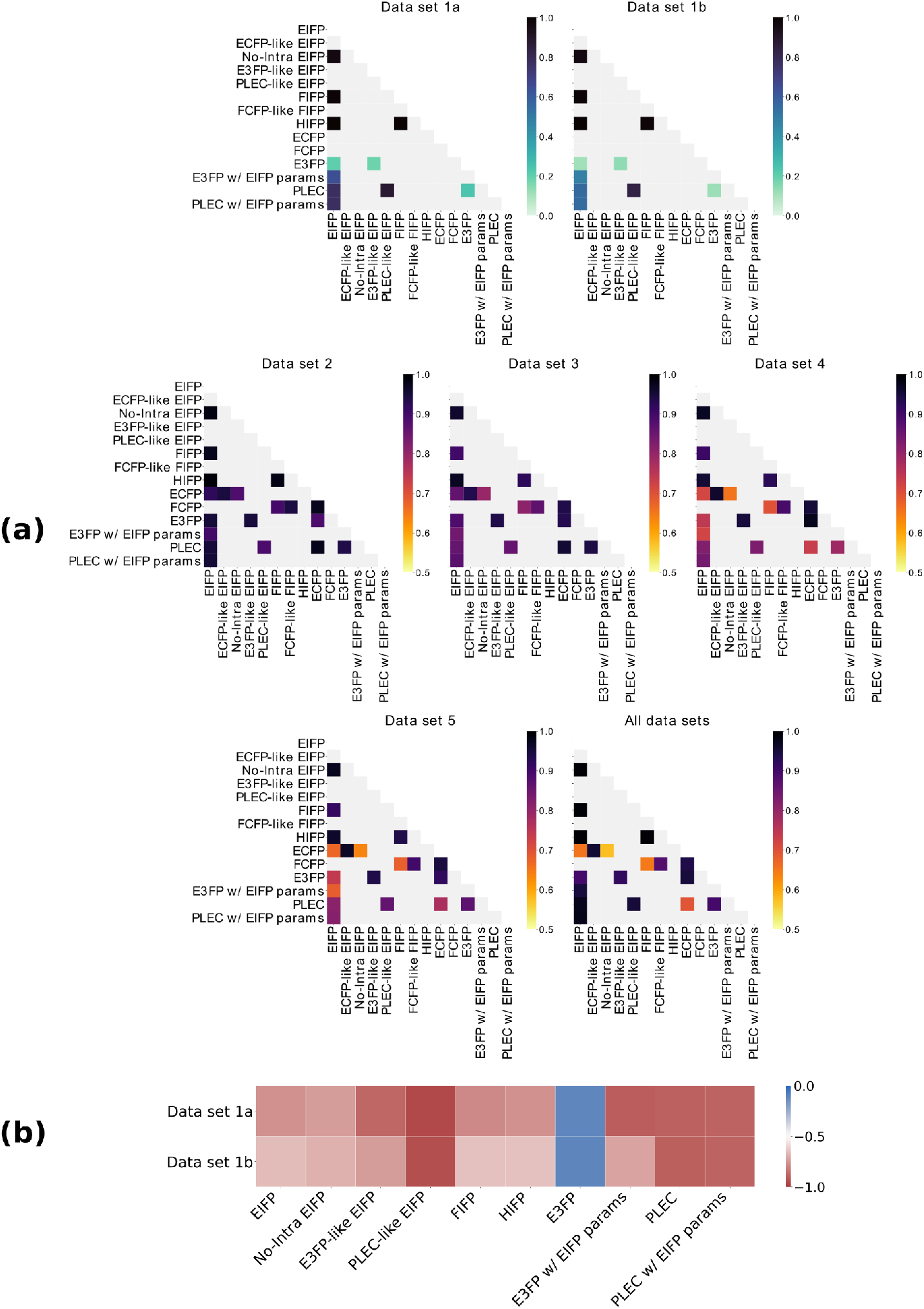
(a) Correlation between different FPs across five data sets with varied protein and ligands diversity. Correlations were obtained by calculating the Tanimoto (*T_C_*) similarity between pairs of FPs of a given type. The overall correlation between different FP pairs was also assessed by joining all data sets into a single one (*All data sets*). Color scales for all data sets vary from 0.5 to 1, except for data set 1 whose color scales vary from 0 to 1. Light gray squares represent untested pairs. (b) Correlation between the ligand pairs’ RMSDs and the FP similarities. The color scale varies from no correlation (blue color) to high negative correlation (red color).

Overall, we found high correlations between the different FP representations in all data sets (Figure 8a) despite the increasing spread of data points and Pearson correlation reduction as we moved towards the more diverse scenarios (Figures S23-S28). This finding supported our expectations regarding the close relationship between molecular topology and interactions when limited to the same protein crystal structure. This result was further supported by the comparison between ECFP and *No-Intra EIFP* for data set 2 (Figure S25), in which molecular topology information was strongly correlated to an interaction-only representation. Other EIFP variations (mimicking other FPs) were also highly correlated with their counterparts despite their differences in pharmacophoric or topological definitions and FP parameters.

Additionally, analysis of data set 1 revealed a negative correlation between the ligands’ RMSDs and the IFP similarities (Figures 8b, S29, and S30). We also observed that similarity data points of IFPs tended to form groups according to the RMSD range (Figures S23 and S24). These expected observations highlighted the capacity of the IFPs and PLEC to identify similar binding modes given by ligands with very similar conformation.

The only exception for the overall RMSD trend was found for E3FP, which did not correlate either with the RMSD data or with EIFP and PLEC (Figure 8). In data set 1, E3FP tended to separate data points into two regions based on FP similarities (first boxes in Figures S29 and S30). However, further separation by RMSD was not possible as low- and high-RMSD data points lay down at the same similarity range. Interestingly, the same has already been observed in Wang et al., ^64^ in which molecules with low RMSD produced small molecular similarities. Only after setting the FP parameters as in EIFP (*E3FP w/ EIFP params*), a reasonable correlation with the RMSD and EIFP emerged. Thus, considering that E3FP was conceived and parameterized to compare different ligands and not conformers of a single ligand, we believe the default configuration may not be appropriate in this scenario as it was not intended to distinguish two poses of the same ligand regardless of its RMSD.

Finally, we also found a near-random correlation between EIFP and PLEC in the context of conformers with high RMSDs (*>* 0.8 Å) at data set 1 (Figure S24). Setting PLEC with the same number of levels and radius growth rate from EIFP did not increase the correlation as observed for E3FP. A high correlation (*r ∼* 0.84) was only recovered with *PLEC-like EIFP*, an EIFP version that mimicked PLEC implementation. Therefore, we believe the low observed correlation between PLEC and EIFP may be caused primarily by the difference between their implementations.

### Interaction fingerprints identify similar poses

The need to rank molecules based on molecular or binding mode similarity comes up frequently in drug discovery. To evaluate the use of IFPs in distinguishing ligands by binding mode similarity, we again turned to data sets 1-5 (Table 3). We compared the IFPs against ECFP, FCFP, E3FP, and PLEC for this task.

In this analysis, we separated the pairs of complexes based on their proteins and ligands into six groups: complexes with the same proteins and ligands; complexes with the same proteins and ligands with high molecular similarity (ECFP *T_C_ >* 0.5); complexes with the same proteins and ligands with low molecular similarity (ECFP *T_C_ ≤* 0.5); complexes with different proteins and the same ligands; complexes with different proteins and ligands with high molecular similarity (ECFP *T_C_ >* 0.5); and complexes with different proteins and ligands with low molecular similarity (ECFP *T_C_ ≤* 0.5). By doing so, we expected to reveal that complexes comprising the same proteins and ligands with high molecular similarity (ECFP *T_C_ >* 0.5) would tend to present the highest IFP similarities as well.

Indeed, we found that ligands with low molecular similarities presented the lowest IFP similarities, whose values continuously reduced as we moved towards the more diverse scenarios (Figure 9). Moreover, similarity distributions for ligands with high (ECFP *T_C_ >* 0.5) and low (ECFP *T_C_ ≤* 0.5) molecular similarities over the same binding site did not overlap with EIFP and PLEC as expected. In contrast, the observed overlapping similarity distributions for complexes comprising the same ligand or ligands with high molecular similarity (ECFP *T_C_ >* 0.5) bound to different proteins indicated that the binding site played the major role in defining the ligands’ binding mode.

**Figure 9:**
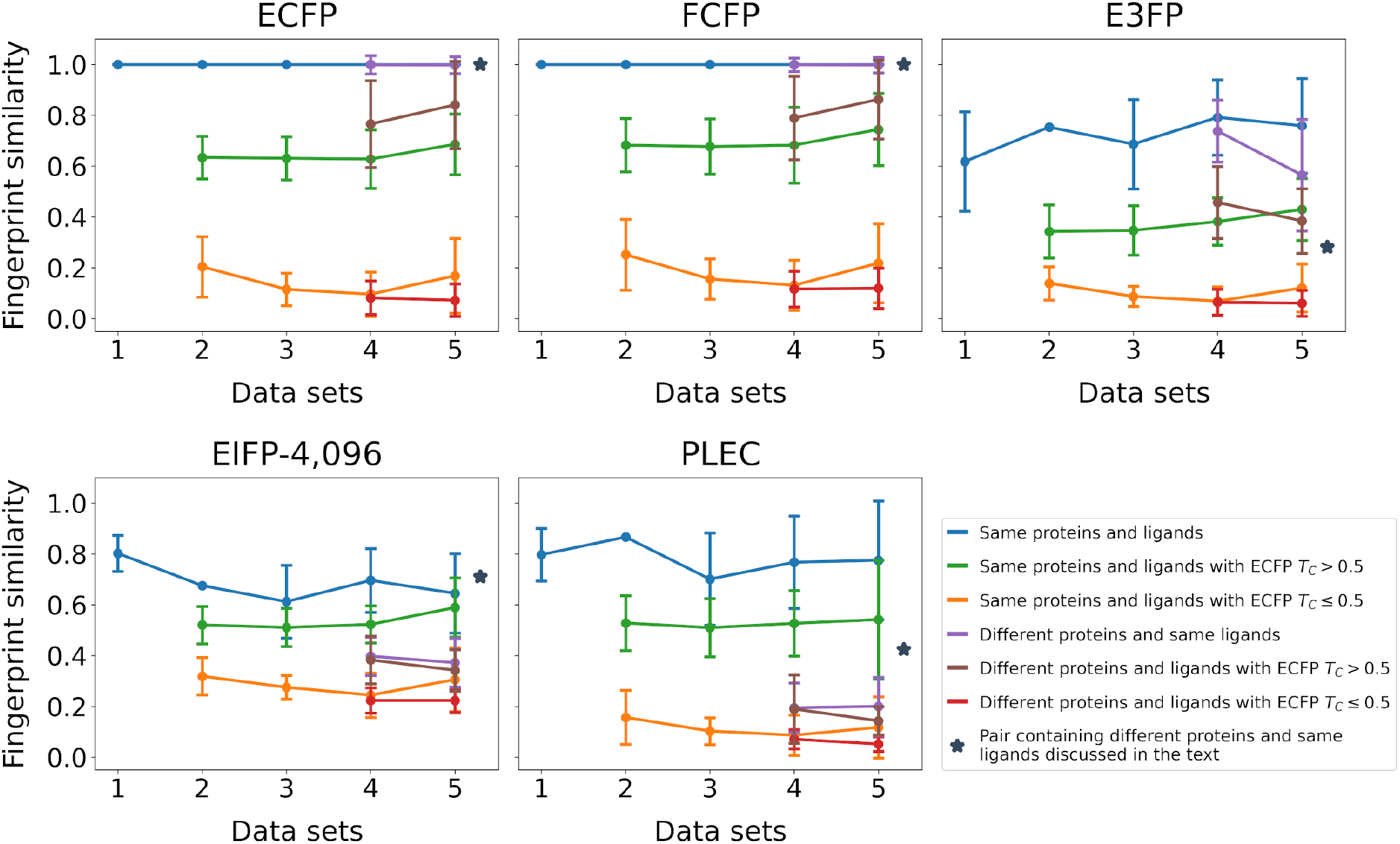
Similarity between FP pairs for the five data sets presented in Table 3. Pairs of complexes were divided into six groups based on their proteins and the molecular similarity (ECFP) between their ligands. Marker represents complexes with same ligands discussed in the text.

Overall, EIFP and PLEC showed similar separability trends across the different data sets and pair types. However, we also identified divergences between these FPs. For instance, the pair of complexes highlighted by the marker in Figure 9 was an example where EIFP produced a high similarity (*∼* 0.71), while PLEC and E3FP attributed to them low similarities (0.42 and 0.28, respectively). Interestingly, these complexes contained different proteins (proteasome beta type-5 subunit, PDB id 5LF4; and beta type-1 proteasome subunit, PDB id 5LF7) bound to the same ligand (pentaethylene glycol). Nonetheless, despite being different proteins, they presented high structural similarities (TM-Score^83^ = 0.857), which explained the high similarity between these two complexes according to EIFP. In Figures S31 and S32, we show common and exclusive features between these two complexes found with EIFP.

Finally, varying EIFP lengths (512, 1,024, 2,048, 8,192, 16,384, and unfolded), or using a different IFP representation (FIFP) did not impact the separability between similar and dissimilar binding modes (Figures S33 and S34). Thus, despite the expectation that shorter FPs would produce more feature collision rates and information losses, we observed consistency on all variations of IFP.

Altogether, these findings demonstrate the capacity of the IFPs to identify complexes with high similarity.

## Discussion

Traditional representations such as MFPs and IFPs have been applied for a long time in drug discovery. Such simple representations are useful for fast filtering or querying a database to identify molecules by structural or binding mode similarity to a set of known molecules and complexes. Although these FPs were also widely applied in ML-based drug discovery, there is still a need for new methods that satisfactorily capture the molecular recognition process and encode it into a representation appropriate for ML tasks. Achieving it would open up opportunities for the accurate automatic selection of promising molecules or the prioritization of molecules by re-scoring them with a predicted binding affinity measure.

These remarkable applications and the long history of FPs motivated us to propose LUNA and three novel hashed IFPs: EIFP, FIFP, HIFP. The IFPs encode molecular complexes and their interactions, and were specially designed for ML tasks and the comparison of complexes by similarity. Moreover, the common lack of interpretability in hashed FPs and ML models applied on drug discovery led us to set up several visual functionalities to trace a feature back to its corresponding context in the binding site. To explore the potential of the IFPs, we set up three major experiments: re-scoring, interpretability, and identification of molecules with similar binding modes.

With the first experiment, our goal was to assess if IFPs could reproduce the observed DOCK3.7 scores as in a re-scoring task. To do so, we leveraged a large dopamine D4 receptor docking data set by Lyu et al. ^72^ and built two random diverse and stratified data sets based on DOCK3.7 scoring bins. The first data set contained 200,000 docking poses and was employed for the identification of the best parameters for LUNA and IFP. Among the evaluated parameters was the FP length, whose value plays a central role in hashed FPs as it controls how many unique substructures may be represented. Short FPs suffer from greater bit collision frequency and, consequently, information loss. In contrast, long FPs cause compute performance and storage issues and may produce overfitted ML models, a problem known as the curse of dimensionality.^84^ Several authors explored different MFP and IFP lengths,^7, 59, 60, 71, 85–95^ however there is no single consensus on length, as results vary with the problem and data sets.

In our work, we explored three different lengths (1,024; 2,048; and 4,096) and evaluated which IFP representation and other LUNA parameters performs better. After exploring 294 parameter choices, the best IFP identified was *EIFP-4,096*. Notably, the HIFP that unites both EIFP and FIFP worlds – pharmacophoric properties for atom groups and precise atom environment information for atoms – presented intermediary results. Overall, these results suggested that precise atomic topology definition majorly impacts the performance of DOCK3.7 score prediction and, by extension, of binding affinity regression tasks. Similar results were already observed for MFPs.^58^

Interestingly, the best IFP configuration included intra-residue interactions. Intuitively, one might have expected that because the protein structure is always the same across all complexes in a single-target docking campaign, the intra-residue interactions would always be the same for the protein structure, and thus yield no meaningful contribution to the model. However, the ligand-pose-specific inclusion of intra-residue interactions and the residues themselves do contribute, by augmenting local ligand context and implicitly providing information about the binding site region occupied by the ligand.

After identifying the best IFP, we trained multiple DNN models on the second data set (1 million molecules docked against the same dopamine D4R structure). We explored multiple DNN architectures and hyperparameter choices for the best EIFP and each alternative FP method, and compared the DNN performances to two baseline ML methods: random forest and XGBoost. We also assessed a longer EIFP version (16,384) to compare its performance with the default PLEC. Overall, DNN models presented the best results with relatively simple model designs. The top three FPs were *PLEC-16,384*, followed by *EIFP-4,096*, and *EIFP-16,384* (Figures 3, S10, and S15). Notably, the difference between the performance of these models was small and, therefore, we believe it may not justify the use of such longer FPs as they may produce overfitted models and increase processing time. Moreover, in our experiments, IFPs presented a superior performance in comparison to all MFPs, a similar observation also found by Rodŕıguez-Pérez et al., ^96^ which highlights the importance of the protein-ligand interaction information.

Besides EIFP’s competitive regression performance, we found that LUNA permits visual assessment of individual FP features, making it possible to scrutinize the binding mode of ligands in PyMOL.^97^ To open the “black box” of ML modeling, we carried out an interpretability study with LUNA using feature importance scores calculated from XGBoost trained on the best EIFP for 1 million dopamine D4 complexes from docking. Of the key interactions that emerged, five features summarized ionic and hydrogen bonds established with ASP115, a key residue present in all aminergic receptors. These features were most frequently among the best ligands and almost absent among the worst ligands.

Finally, in the third experiment, we compared the IFPs to “baselines” of standard molecular (ECFP, FCFP, and E3FP) and interaction (PLEC) FPs employing five data sets with increasing degrees of complexity by protein and ligand variability. We assessed how IFP-based similarity corresponded to baseline-FP similarities, as a function of differing molecular and interaction information. Secondly and most importantly, we evaluated the IFPs’ capacity to identify similar vs dissimilar poses no matter the data set diversity, which is a task commonly performed in virtual screening campaigns.

We found medium to high correlations between the FPs in all data sets. In part, high correlations are consistent with the common core strategy of defining structural features (circular substructures centered around an atom or atom group) and with the similarities across atomic or pharmacophoric properties definitions. This applies especially for MFPs, for which previous studies report high cross-correlations. ^59, 92^ Accordingly, ligands with high molecular similarity tended to present similar binding modes within the same binding site. Indeed, we observed that molecular topology (ECFP) was strongly correlated to an interaction-only representation (*No-Intra EIFP*) when considering only CDK2 complexes (data set 2).

IFPs effectively captured ligand pose similarity. Overall, we found high EIFP similarities especially for pairs of complexes containing the same proteins and ligands in different conformations. Second, IFPs distinguished among ligand pairs with high (ECFP *T_C_ >* 0.5) versus low (ECFP *T_C_ ≤* 0.5) molecular similarities within the same binding site, speaking to differences as well in their binding modes. Conversely, we identified several pairs of complexes for differing proteins and ligands with, nonetheless, high EIFP similarities, supporting the IFPs’ capacity to capture similar protein-ligand complexes, rather than similar ligand structures alone.

Overall, we envision LUNA and the novel hashed IFPs as unique tools to advance drug discovery. First, LUNA provides powerful functionalities for analyzing protein-ligand complexes obtained by docking or molecular dynamics. Second, the IFPs can accelerate virtual screening campaigns by identifying molecules with binding modes similar to a set of desired molecules. Last, the IFPs were explicitly designed for ML tasks such as ligand-protein complex re-scoring and selection of promising compounds. Indeed, in a recent example by Alon et al.,^20^ the authors obtained improved hit-rate selection and compound prioritization using a pre-release prototype of LUNA and EIFP.

To conclude, EIFP outperformed standard FPs (ECFP, FCFP, E3FP, and PLEC) representations in our ML use-case studies. Still, the task of reproducing human-made binding scores, which are considerably simpler than experimental binding energy measures, leaves space for improvement. Despite the consistency of literature results^2, 5, 6, 62, 98–100^ with our findings, future work might explore whether the inclusion of unsatisfied hydrogen bonds and other terms such as desolvation penalties can further improve the performance of DNN models trained with IFPs. Moreover, in our work, geometric criteria were used only during the calculation of PLIs, but not during the FP creation. Thus, we hypothesize that including distances and angles for electrostatic interactions within the representation may likewise improve the prioritization of hit compounds and protein-ligand complex discovery through IFPs.

## Methods

### Physicochemical properties of atoms and atom groups

Departing from our previous work^101^ and a literature search,^42, 51, 74, 102–123^ we classified atoms and groups of atoms according to their physicochemical properties into one or more of the following types: *acceptor*, *amide*, *aromatic*, *atom*, *chalcogen donor*, *donor*, *electrophile*, *halogen acceptor*, *halogen donor*, *hydrophobe*, *hydrophobic*, *metal*, *negatively ionizable*, *nucleophile*, *positively ionizable*, *weak acceptor*, and *weak donor*. In LUNA, an atom group can represent both chemical functional groups or simply an arrangement of atoms as in *hydrophobes*. The latter is an optional property that represents a group of hydrophobic atoms and better mimics how the hydrophobic effect occurs (i.e., a favorable contact between two hydrophobic surfaces). It is also important to highlight that although atoms may belong to a group, they all have their own physicochemical properties.

The terms *negatively ionizable* and *positively ionizable* were chosen to indicate that a specific atom or group of atoms may be ionized. However, if necessary, one can set a different pH in order to alter the resulting classification of an atom or group.

For more details, refer to Section *Physicochemical feature assignment* in the Supporting Information.

### Molecular interactions calculation

For each pair of atoms or group of atoms, molecular interactions are characterized using physicochemical properties and geometrical criteria (distance and angle). For atom groups, their centroids are used in the geometrical analysis.

LUNA identifies the following interactions: *amide-aromatic stacking*, *anion-electrophile*, *antiparallel multipolar*, *cation-nucleophile*, *cation-pi*, *chalcogen bond*, *chalcogen-pi*, *covalent bond*, *displaced face-to-edge pi-stacking*, *displaced face-to-face pi-stacking*, *displaced face-to-slope pi-stacking*, *edge-to-edge pi-stacking*, *edge-to-face pi-stacking*, *edge-to-slope pi-stacking*, *face-to-edge pi-stacking*, *face-to-face pi-stacking*, *face-to-slope pi-stacking*, *halogen-pi*, *halogen bond*, *hydrogen bond*, *hydrophobic*, *ionic*, *multipolar*, *orthogonal multipolar*, *parallel multipolar*, *pi-stacking*, *repulsive*, *salt bridge*, *tilted multipolar*, *unfavorable anion-nucleophile*, *unfavorable cation-electrophile*, *unfavorable electrophile-electrophile*, *unfavorable nucleophile-nucleophile*, *van der Waals*, *water-bridged hydrogen bond*, and *weak hydrogen bond*.

Besides interactions, LUNA also provides three contacts: *atom overlap*, *proximal*, and *van der Waals clash*. *Atom overlap* identifies artifacts generated by low-resolution structures and homology models, which consist of the unnatural overlap of two atoms. *Proximal* is an optional contact, which simply indicates that two atoms are close to each other by a specific threshold. And *van der Waals clash* characterizes the repulsion between two atoms when they become too close.

The combination of physicochemical features, distance, angle criteria, and geometrical models utilized to calculate interactions are presented in the Supporting Information. Although LUNA implements its own methods and criteria to calculate interactions, users can also define their own functions and cutoffs – thanks to the object-oriented style employed in LUNA.

### Interaction fingerprints

We propose three novel hashed IFPs called EIFP, FIFP, and HIFP, which are inspired by ECFP, FCFP,^58^ and E3FP.^59^ Similar to ECFP, FCFP, and E3FP, our IFPs depend on three parameters: the FP length, the radius growth rate, and the number of levels (Figure 1). Below, we detail how the IFPs are generated.

#### Generating initial identifiers

At iteration 0, initial identifiers are assigned to each atom or atom group. In EIFP, the atoms are assigned seven invariant atomic properties derived from ECFP and E3FP: the number of heavy atoms covalently bound to the atom; the valence minus the number of neighboring hydrogens; the atomic number; the atomic isotope number; the atomic charge; the number of bound hydrogens; and a flag indicating whether the atom belongs to a ring or not. For atom groups, EIFP includes the initial identifier of each atom comprising the group. As a result, EIFP precisely characterizes topological information like ECFP.

In FIFP, only physicochemical features are considered during the identifier generation. For atoms, FIFP assigns their own physicochemical features and features from any atom group to which they belong. For atom groups, FIFP encodes only the physicochemical features of the group. For example, the chemical feature *Aromatic* assigned for an aromatic ring (atom group) is also included as the feature of each one of its atoms. Therefore, FIFP represents more abstract role-based sub-structural features.

Finally, HIFP is a hybrid flavor that encodes atomic invariants for atoms and physico-chemical features for atom groups.

After characterizing each atom or atom group according to the IFP flavor, their information is encoded to their initial 32-bit integer identifiers using a hash function. LUNA uses a Python implementation^2^ of MurmurHash3^124^ hash function. However, any hash function can be applied as long as it generates uniform and random identifiers.

#### Subsequent identifiers update

After generating each initial identifier, the algorithm subsequently updates them through an iterative process that continues until it converges or it reaches the maximum number of levels, which is given by the parameter *Number of levels* (Figure 1). At each iteration, a sphere of size *R ∗ L*, where *R* is the *radius growth* and *L* the current level (iteration number), is centered at each atom and atom group. Their neighborhood is then characterized by capturing all interactions within the shell. A hash function is applied to the neighborhood, and a new identifier to the central atom or atom group is generated.

A single iteration for a given atom or atom group is performed as follows. First, LUNA centers a sphere of size *R ∗ L* on an atom or atom group, and initializes an array of tuples with a pair consisting of the current level number and the previous identifier of this atom or atom group. Next, it captures all atoms (atom groups) inside the shell and verifies if these entities establish any interaction with the atoms or atom groups from the previous iteration. Then, for each identified interaction, LUNA generates a tuple containing the interaction type and the previous identifier of the interacting partner. These pairs are sorted and included in the array of tuples. Note that sorting the list is essential for avoiding dependence on the order of its elements; otherwise, the FP generation would not be deterministic.

Finally, LUNA generates a new 32-bit integer identifier to the central entity by hashing the obtained list. Any atom or atom group interacting with it becomes part of its neighborhood to be considered in the subsequent levels.

This iterative process continues for each atom or atom group until it reaches the maximum number of levels or until the algorithm converges, which happens when the shells cannot be expanded anymore. In other words, all interactions established by the atoms and atom groups of all shells have already been added in the last iteration.

### Machine learning data set

We obtained *∼*96 million molecules from a large data set recently published by Lyu et al..^72^ This huge library consists of molecules docked against Dopamine D4 and comprises 6,453,455 unique cluster heads (molecules). From these, 90% of the cluster heads were randomly selected for model development and training, while the remaining 10% were set apart as a hold-out set and were not released until the model development phase of the study had been completed. Finally, to obtain diverse and stratified sets, we divided the cluster heads into 10 equal-sized bins by their DOCK3.7 score and randomly sampled 90,000 and 10,000 molecules from each bin to form the expanded training (900,000) and testing (100,000) data sets, respectively.

Using the same procedure and bin sizes, we defined another independent training subsample composed of 200,000 molecules specific for hyperparameter optimization of LUNA and IFP parameters.

### Model development and training

The DNN models were trained using the open-source package Pytorch^125^ coupled with Skorch, ^126^ and the hyperparameter search was powered by the library Optuna.^127^ In all experiments, we used the MSELoss criterion. We chose the scikit-learn implementation of random forest and XGBoost^128^ with default parameters as the baseline ML methods. During model development and training, 20% of the training data set was used for model validation. Models’ performance was evaluated in terms of the *R*^2^ metric.

Regarding baseline models, we chose the RDKit implementations of ECFP and FCFP. Additional molecular (E3FP) and interaction (PLEC) FPs were also selected for further comparison. The length of ECFP, FCFP, and E3FP were set to 4,096. In contrast, we used two lengths for PLEC: 4,096 or 16,384 (the default). In addition, we turned on counting information and turned off stereochemical information for E3FP according to Axen et al..^59^ For LUNA and IFP hyperparameter optimization, models were trained using the training subsample (200,000 complexes) and the best DNN architecture found in previous investigations (results not shown). We explored combinations of ten different parameters (Table S1).

For the best IFP and for the chosen alternative FP methods, models were trained using the complete training set (900,000 complexes) and the best DNN architecture for each FP was identified through hyperparameter search on the space shown in Table S3. In total, 165 and 30 different parameter configurations were evaluated for FPs of length 4,096 and 16,384, respectively. Additionally, the following preprocessing and regularization strategies were employed: counting information (where applicable) and DOCK3.7 scores normalized using log1p (log(1+*x*)) and min-max normalization (MinMaxScaler^129^), respectively; weight decay (L2 regularization); batch normalization; dropout; and early-stopping with patience equal to 20 and 30 for FPs of length 4,096 and 16,384, respectively, and at most 3,500 epochs. For FPs of length 16,384, a minimum of 100 epochs were run before early-stopping was activated. Besides the standard dropout strategy where the same probability was employed in all dropout units, we also applied a procedure where the dropout probability was decreased at an exponential decay rate – shallow hidden layers had a greater decay.

Finally, we assessed the best models’ stability using 5-fold cross-validation for the DNN, random forest, and XGBoost methods. To guarantee reproducibility, we used an initial seed (54,343) to generate five random integer values that were used as seeds to randomly define the hold-out set and initialize the DNN models at each fold iteration. Mean and standard deviation were then calculated for the *R*^2^ values obtained from the 5-fold cross-validation strategy.

### Model interpretability

To illustrate how the out-of-the-box LUNA functionalities are suitable for creating interpretable ML models, we performed a feature attribution analysis based on the feature importance scores calculated by the XGBoost library.

The XGBoost model was trained using *EIFP-4,096* (the best FP identified in the ML case study) and 80% of the training set (Section *Model development and training*).

To make sense of a feature and its importance, LUNA was used to assign a class to each one of the 4,096 features according to the information they contained, where the class was one of the following: *a*) *ligand’s level 0 features only* or *protein’s level 0 features only* indicating whether it represented a level 0 shell containing only ligand’s or protein’s atomic information, respectively; *b*) *upper level with ligand atomic information only* or *upper level with protein atomic information only* indicating whether it represented upper shells containing only ligand’s or protein’s atomic information, respectively; *c*) *intra-ligand interactions only* or *intra-protein interactions only* indicating whether it represented shells containing only intra-ligand or intra-protein covalent and non-covalent interactions, respectively; *d*) *has non-covalent interactions with the protein* indicating whether any non-covalent interactions between the protein and the ligand was identified in the shell; or *e*) *features with collision in the same complex* whether two or more distinct shells from the same complex were hashed into the same bit position.

As bit collision resulting from FP folding may generate features with different information, LUNA may assign multiple classes to them. To deal with this expected artifact of hash functions, we first calculated the percentage of the most prevalent class in a given feature and used these percentages to calculate z-scores for each feature. We then filtered out features presenting z-scores higher than 1 and chose the minimum percentage (69.49) among the remaining features as the threshold to attribute a class to each feature (Figure S35). Thus, the most prevalent class was assigned to a feature only if its percentage was greater than or equal to the threshold. Whenever a feature position did not meet this condition, we attributed the class *unreliable feature* to it.

To identify the set of features that played the major contribution to the model performance, we calculated the z-scores for each feature using their importance scores. As data followed a generalized extreme value distribution (Figure S36), we transformed z-scores to p-values as in Keiser and Hert: ^130^ 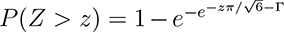, where *z* is a z-score value and Γ is the Euler–Mascheroni constant (*≈* 0.577215665). Most important features were then defined as those having p-values less than 0.01. The importance scores, z-scores, p-values, and classes attributed to each feature are available as an Excel file in Supporting Information.

### Fingerprint assessment data sets

To assess the proposed IFPs, we built five different data sets. For the first data set, we automatically generated two series of conformers for the compound 2-(allylamino)-4-aminothiazol-5-yl(phenyl) methanone (PDB id X02) in complex with the human cyclin-dependent kinase 2 (CDK2; PDB id 3QQK). Conformers were generated with the function EmbedMultipleConfs from RDKit with the parameter *numConfs* and *pruneRmsThresh* set to 10,000 and 0.1 Å, respectively. The first parameter defined the number of conformers the algorithm should generate, while the second removed conformers whose distance (RMSD) to other conformers was less than 0.1 Å. This pruning procedure only retained the first conformation generated. From then on, only those that were at least pruneRmsThresh away from all retained conformations were kept. After generating the conformers, we aligned them to the ligand X02 and removed the conformers whose distance to the crystal pose was higher than 0.4 Å (Figure S37a) or 0.8 Å (Figure S37b). As a result, we obtained 53 and 431 conformers, respectively.

The second data set was obtained from Schonbrunn et al.. ^82^ In their work, the authors proposed 95 analogs by systematically modifying the flanking allyl and phenyl moieties of the ligand X02. Structures of 36 CDK2-ligand complexes were solved through X-ray crystallography. We also found in the PDB an additional 38 related structures that were deposited by the authors, totaling 74 complexes in this data set.

The third data set was obtained from RCSB PDB in May 2020 after searching for structures that present 100% sequence identity to CDK2 (350 PDB ids in total). We then used LUNA to identify PDB structures containing exactly one ligand, excluding crystallography artifacts, ions, and cofactors (Tables S7, S8, and S9).

Finally, the fourth and fifth data sets were obtained from RCSB PDB in May 2020 through its custom report API. The former comprises non-CDK2 kinases filtered from the recovered report based on their Uniprot identifiers (1,364 PDB ids in total). The latter consists of a randomly selected diverse class of proteins except kinases (37,507 PDB ids in total). To avoid bias towards a specific receptor, we generated a random subsample of both data sets using the *Spreadsubsample* algorithm from Weka^131^ with at most a 5:1 difference in receptor frequencies. We then used LUNA to identify PDB structures containing exactly one ligand, excluding crystallography artifacts, ions, and cofactors (Tables S7, S8, and S9). For these data sets, only crystal structures with X-ray resolution below 2 Å were retained.

### Comparing LUNA fingerprints to other fingerprint methods

To compare the IFPs to other molecular (ECFP, FCFP, and E3FP) and interaction (PLEC) FPs, we computed the similarity between all pairs of complexes in data sets 1-5 (Section *Fingerprint assessment data sets*) using the Tanimoto coefficient (*T_C_*). We then measured the degree of correlation between these different FPs using the Pearson correlation coefficient.

Bit FPs were generated using default parameters from each method. Interactions for the IFPs were calculated using all LUNA’s default parameters. Molecular structures for the baseline methods were prepared as described in Section *Preparing molecular structures for ECFP, FCFP, E3FP, and PLEC*.

We also generated different flavors of EIFP and FIFP as a means to mimic the other baseline methods. *ECFP-like EIFP*, for instance, only contains intra-ligand covalent interactions exactly as ECFP. The complete list of features included in each IFP flavor is presented in Table 2. We easily obtained these different IFP variations through flags that control LUNA behavior or by providing filtering functions.

For each pair of conformers from data set 1, we also calculated their RMSD using RDKit. Then, Pearson correlation coefficient between RMSD values and interaction FP similarities was measured.

### Preparing molecular structures for ECFP, FCFP, E3FP, and PLEC

To prepare the inputs for alternative FP methods, we used the same LUNA functionalities employed for the IFPs. First, we extracted compounds from each protein-ligand complex; standardized their topological structure based on their SMILES representation obtained from the Ligand Expo database;^132^ properly charged the molecules considering a physiological environment; and converted them from PDB to MOL. Finally, we filtered out molecules whose number of heavy atoms differed by more than one atom compared to their expected topological structure or whose structure did not match its expected topology. The structural match between ligands was computed using the maximum common substructure algorithm (FindMCS) from RDKit.

## Conclusion

Herein, we propose three novel hashed IFPs: EIFP, FIFP, and HIFP. Our FPs are inspired by ECFP, FCFP, and E3FP, and encode protein-ligand interactions and ligand environment information as either bit or count FPs. Additionally, we propose LUNA^3^, a novel object-oriented Python 3 toolkit that permits researchers to control every processing step during the analysis of a drug discovery campaign. LUNA also provides functionalities for visualizing interactions and FP features to open up the “black box” of ML methods. These characteristics make LUNA IFPs suitable for shedding light on the molecular recognition process and ML models, as well as for prioritizing compounds in virtual screening campaigns.

In this work, we also validated and demonstrated that the IFPs are capable of distinguishing ligands by pose similarity using data sets of differing size and structural diversity. We also assessed how different feature representations impact these analyses and compared our results to existing MFPs (ECFP, FCFP, and E3FP) and to an additional IFP (PLEC).

Finally, to illustrate how LUNA FPs can be applied to binding affinity prediction, we presented a case study wherein we trained ML models on a data set containing 1 million molecules docked against the Dopamine D4 receptor to reproduce calculated Dock scores. We also applied ECFP, FCFP, E3FP, and PLEC to the same study case. Overall, *EIFP-4,096*’s performance was comparable to substantially longer FPs (e.g., *EIFP-16,384* and *PLEC-16,384*) and superior to other FPs of the same length. Finally, we demonstrated how key interactions emerge from feature importance calculations on the FPs in an interpretability study.

## Supporting information

Importance scores and classes attributed to each EIFP feature

List of parameters covered for each FP evaluated during the DNN hyperparameter search and their resulting R<sup>2</sup> values

List of all 294 parameter combinations evaluated during the optimization of LUNA's and IFP parameters and their resulting R<sup>2</sup> values

## Author Contributions

Conceptualization, A.V.F., R.S.F., M.J.K., and R.C.M.M.; Methodology, A.V.F., R.S.F., M.J.K., and R.C.M.M.; Software, A.V.F.; Validation, A.V.F., L.S., L.P., and J.M.; Formal Analysis, A.V.F.; Investigation, A.V.F.; Resources, M.J.K; Data Curation, M.J.O.; Writing – Original Draft, A.V.F.; Writing – Review & Editing, all authors; Visualization, A.V.F.; Supervision, R.S.F., M.J.K., and R.C.M.M.; Project Administration, A.V.F., R.S.F., M.J.K., and R.C.M.M.; Funding Acquisition, M.J.K and R.C.M.M.

## Notes

The authors declare no competing financial interest.

## Abbreviations

CDK2: cyclin-dependent kinase 2
D4R: D4 receptor
DNN: deep neural networks
E3FP: Extended Three-Dimensional FingerPrint
ECFP: Extended-Connectivity FingerPrint
EIFP: Extended Interaction FingerPrint
FCFP: Functional-Class FingerPrint
FIFP: Functional Interaction FingerPrint
FP: fingerprint
GPCR: G protein-coupled receptor
HIFP: Hybrid Interaction FingerPrint
IFP: interaction fingerprint
MFP: molecular fingerprint
ML: machine learning
PLEC: Protein–Ligand Extended Connectivity
PLI: protein-ligand interaction
RMSD: root mean square deviation
SBVS: structural-based virtual screening
Std: standard deviation
*T_C_*: Tanimoto coefficient.

## Acknowledgement

This work has been supported by Coordenação de Aperfeiçoamento de Pessoal de Nível Superior (CAPES) through the process 23038.004007/2014-82, Conselho Nacional de Desenvolvimento Científico e Tecnológico (CNPq), and Fundação de Amparo à Pesquisa do Estado de Minas Gerais (FAPEMIG), as well as by grant number 2018-191905 from the Chan Zuckerberg Initiative DAF, an advised fund of the Silicon Valley Community Foundation (to M.J.K.), Pfizer (to M.J.K), a CTSI TL1 Postdoctoral Fellowship (to J.M.), NIH T32 GM067547 (L.S.), and the UCSF Graduate Division (L.S.). None of the funding agencies influenced the study design and collection, analysis and interpretation of data, nor the writing of the manuscript.

## Footnotes

1 https://github.com/keiserlab/LUNA

2 https://pypi.org/project/mmh3/

3 https://github.com/keiserlab/LUNA

