## Supporting information - additional tables and figures for "Prioritizing virtual screening with interpretable interaction fingerprints"

Table S1: Space covered during the optimization of PLI and IFP parameters.

| Parameter | Value |
|---|---|
| **PLI parameters** | |
| Include residue-residue interactions | Yes / No |
| Include non-covalent interactions | Yes / No |
| Include covalent interactions | Yes / No |
| Include atom-atom contacts (atom overlap, van der Waals clash, and van der Waals) | Yes / No |
| **IFP parameters** | |
| IFP length | 1,024, 2,048, or 4,096 |
| Number of levels | [1, 9] |
| Radius growth radius | [0.1, 7] |
| Protein and ligand generate independent identifiers | Yes / No |
| Feature representation | Bit / Count |
| Type | EIFP, FIFP, or HIFP |

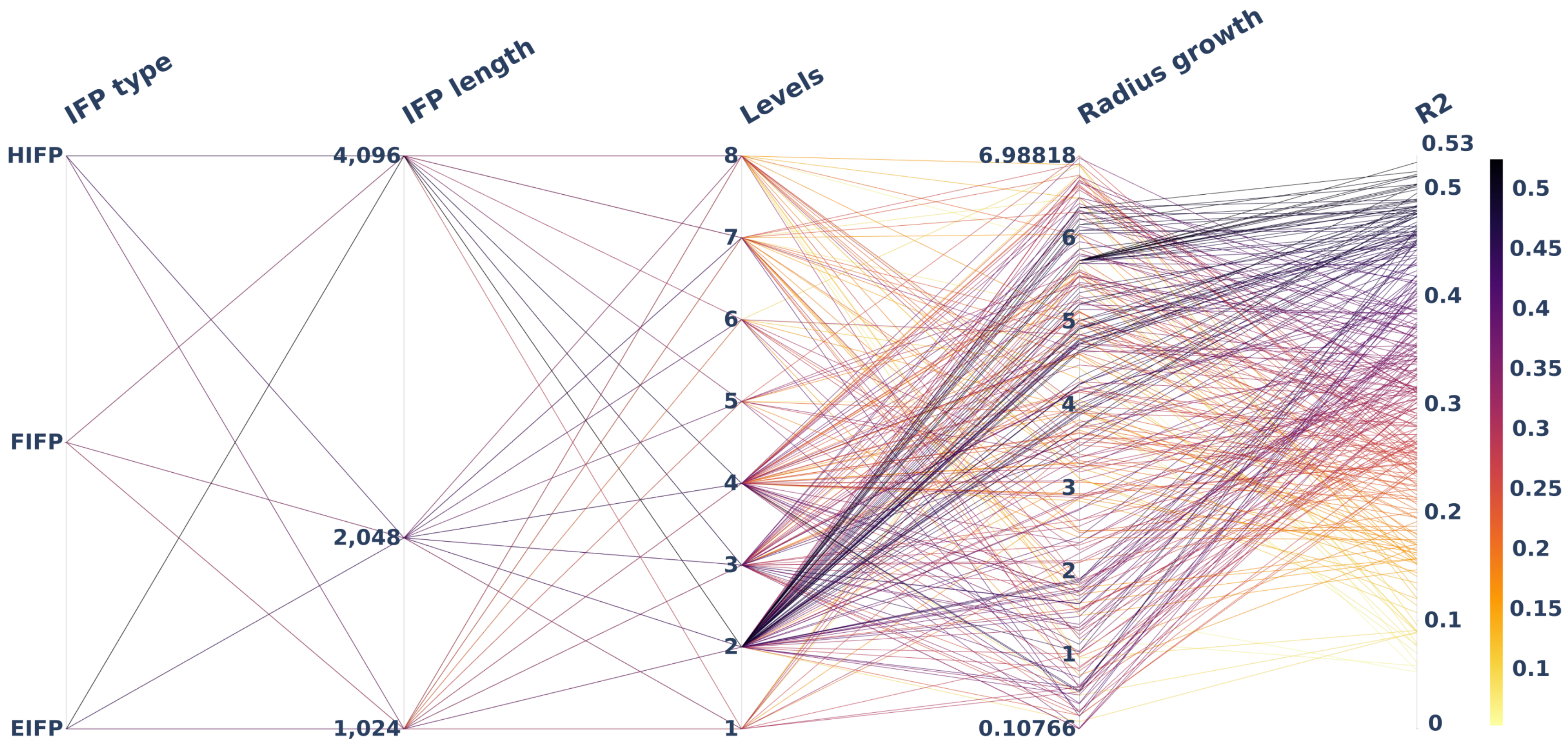


Figure S1: Hyperparameter space covered for the IFP parameters: type, length, number of levels, and radius growth rate. The color scale varies with the $R^2$ from 0 (lighter color) to 0.53 (darker color).

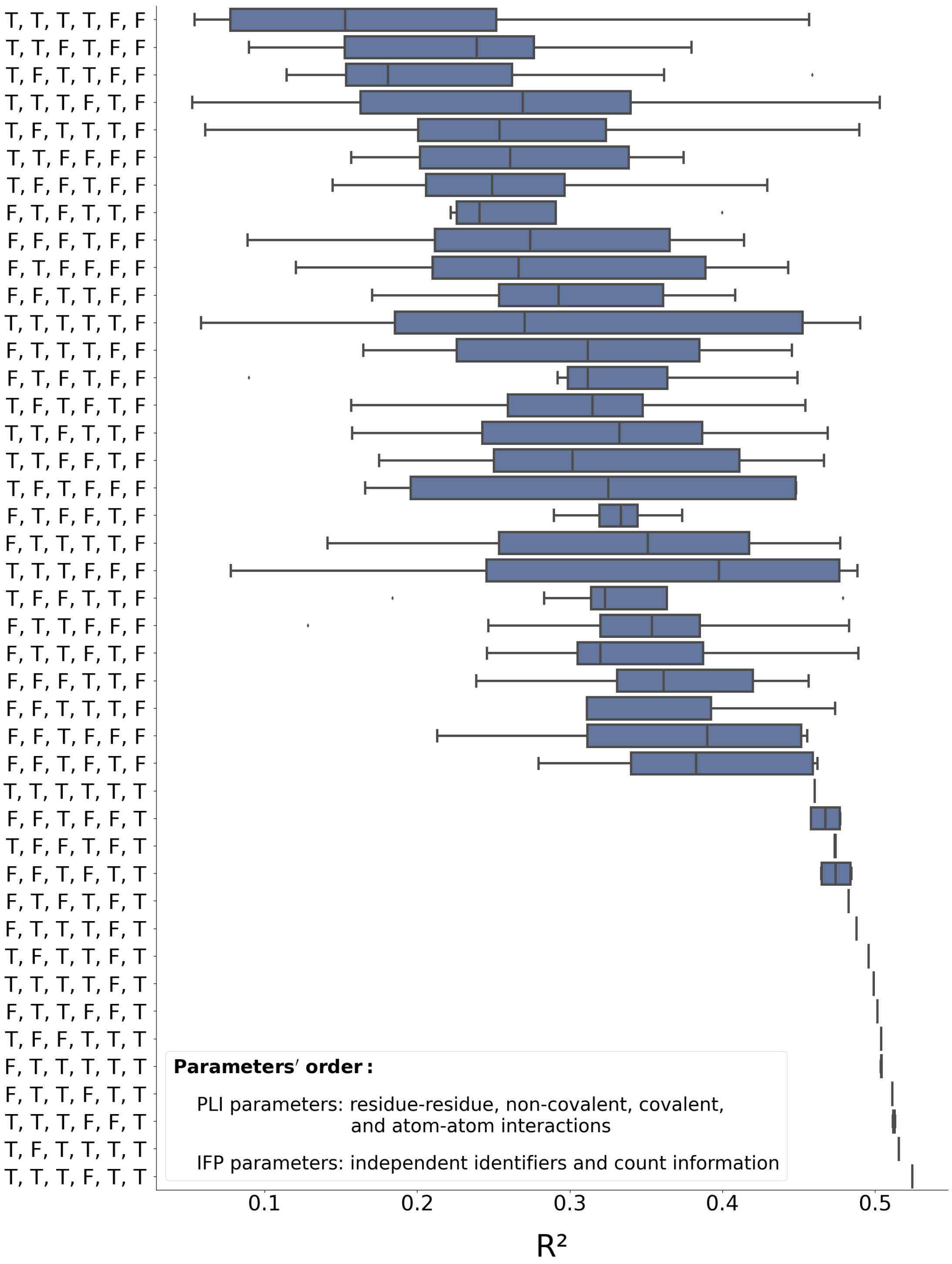


Figure S2: Influence of different Boolean flag combinations on the model performance. The box plots represent all observed $R^2$s for a given parameter combination during PLI and IFP hyperparameter search.

Table S2: Best observed PLI and IFP parameters.

| Parameter | Value |
|---|---|
| **PLI parameters** | |
| Include residue-residue interactions | Yes |
| Include non-covalent interactions | Yes |
| Include covalent interactions | Yes |
| Include atom-atom contacts (atom overlap, van der Waals clash, and van der Waals) | No |
| **IFP parameters** | |
| IFP length | 4,096 |
| Number of levels | 2 |
| Radius growth radius | 5.73171 |
| Protein and ligand generate independent identifiers | Yes |
| Feature representation | Count |
| Type | EIFP |
| $\boldsymbol{R^2}$ | **0.524** |

Table S3: Parameter space covered during DNN hyperparameter optimization.

| Parameter | Values |
|---|---|
| Batch size | 32, 64, 128, 256, and 512 |
| Dropout | [0, 0.35]* |
| Dropout decay | [0, 0.5]* |
| Optimizer | Adam |
| Learning rate | [8e-8, 6e-7]† |
| Weight decay | [1e-3, 1e-1]† |
| Architecture | see Excel file in the Supporting Information. |

* Uniform distribution † Reciprocal distribution (log-uniform distribution).

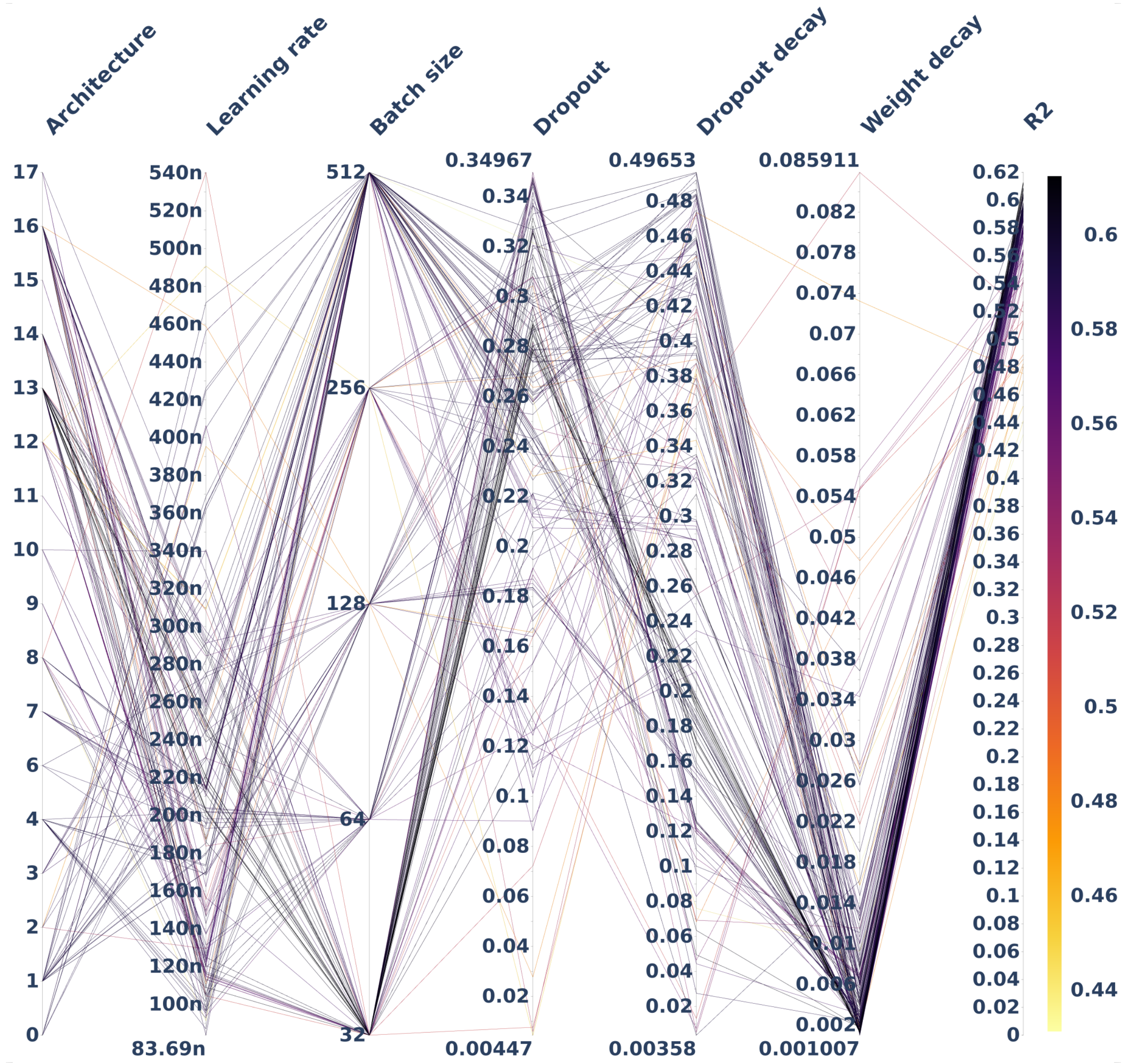


Figure S3: DNN hyperparameter space covered for *EIFP-4,096*. The covered architectures are presented by their identifier and their structures are available as an Excel file in the Supporting Information. The color scale varies with the $R^2$ from 0 (lighter color) to 0.62 (darker color).

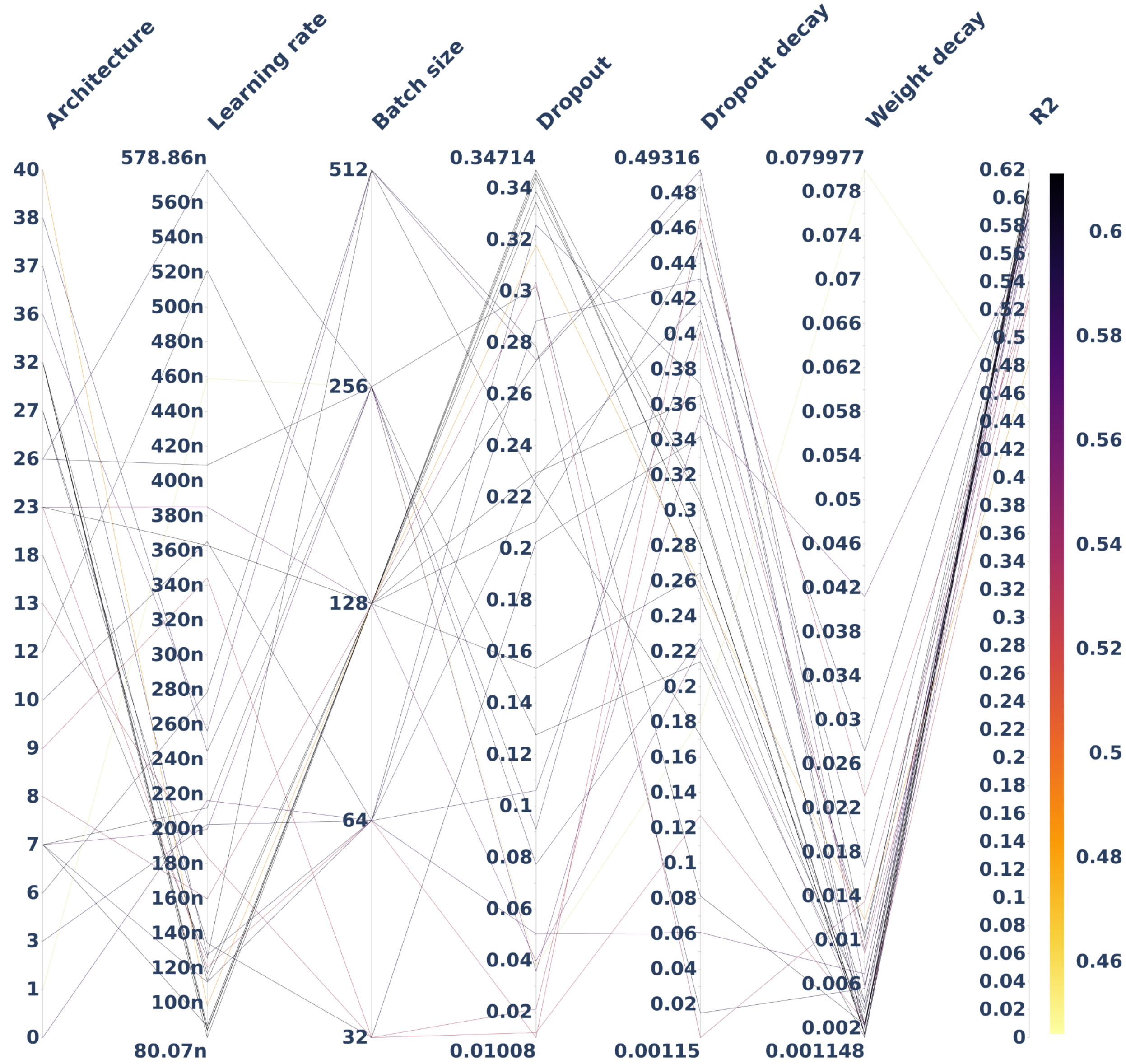


Figure S4: DNN hyperparameter space covered for *EIFP-16,384*. The covered architectures are presented by their identifier and their structures are available as an Excel file in the Supporting Information. The color scale varies with the $R^2$ from 0 (lighter color) to 0.62 (darker color).

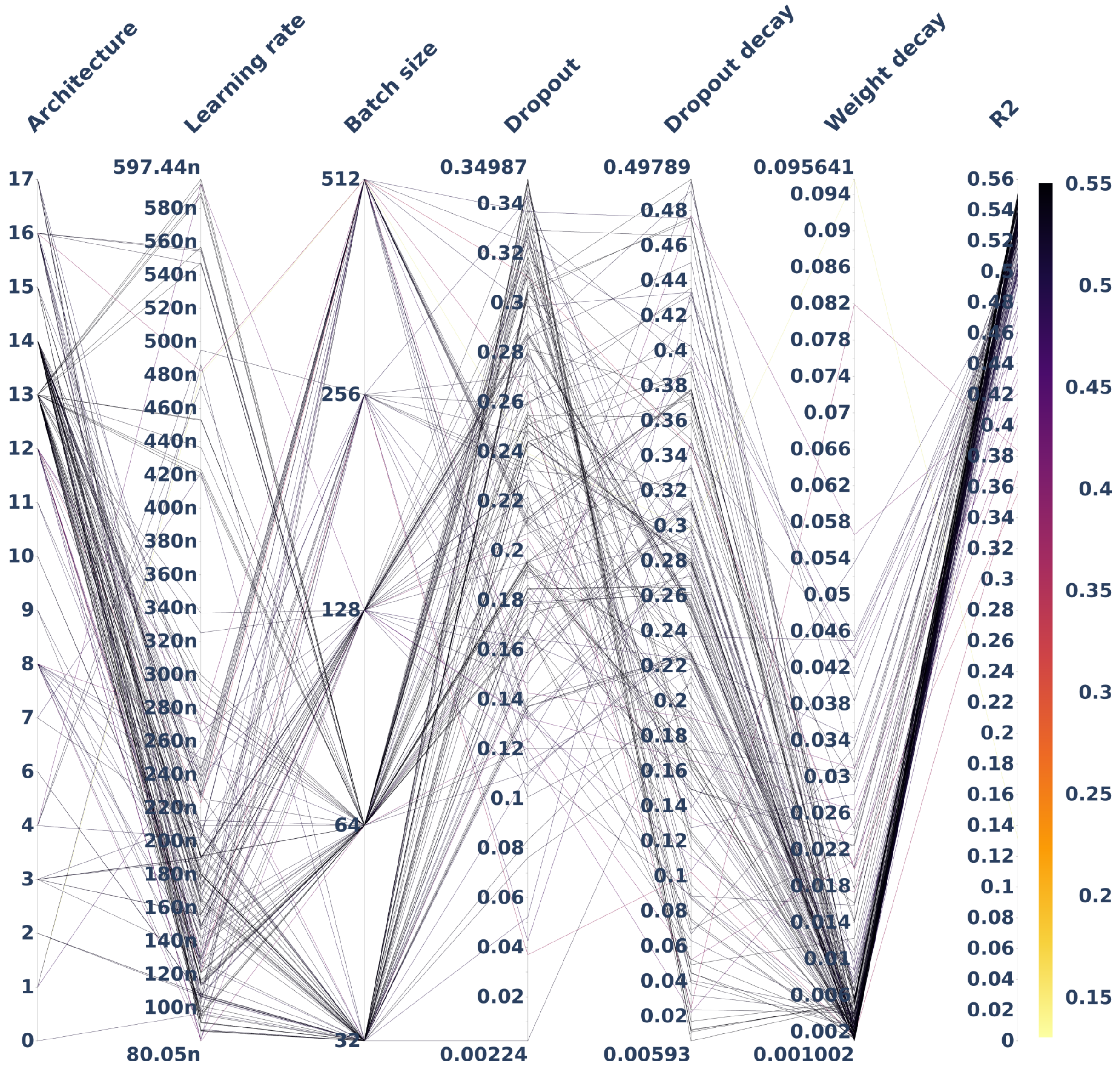


Figure S5: DNN hyperparameter space covered for ECFP. The covered architectures are presented by their identifier and their structures are available as an Excel file in the Supporting Information. The color scale varies with the $R^2$ from 0 (lighter color) to 0.56 (darker color).

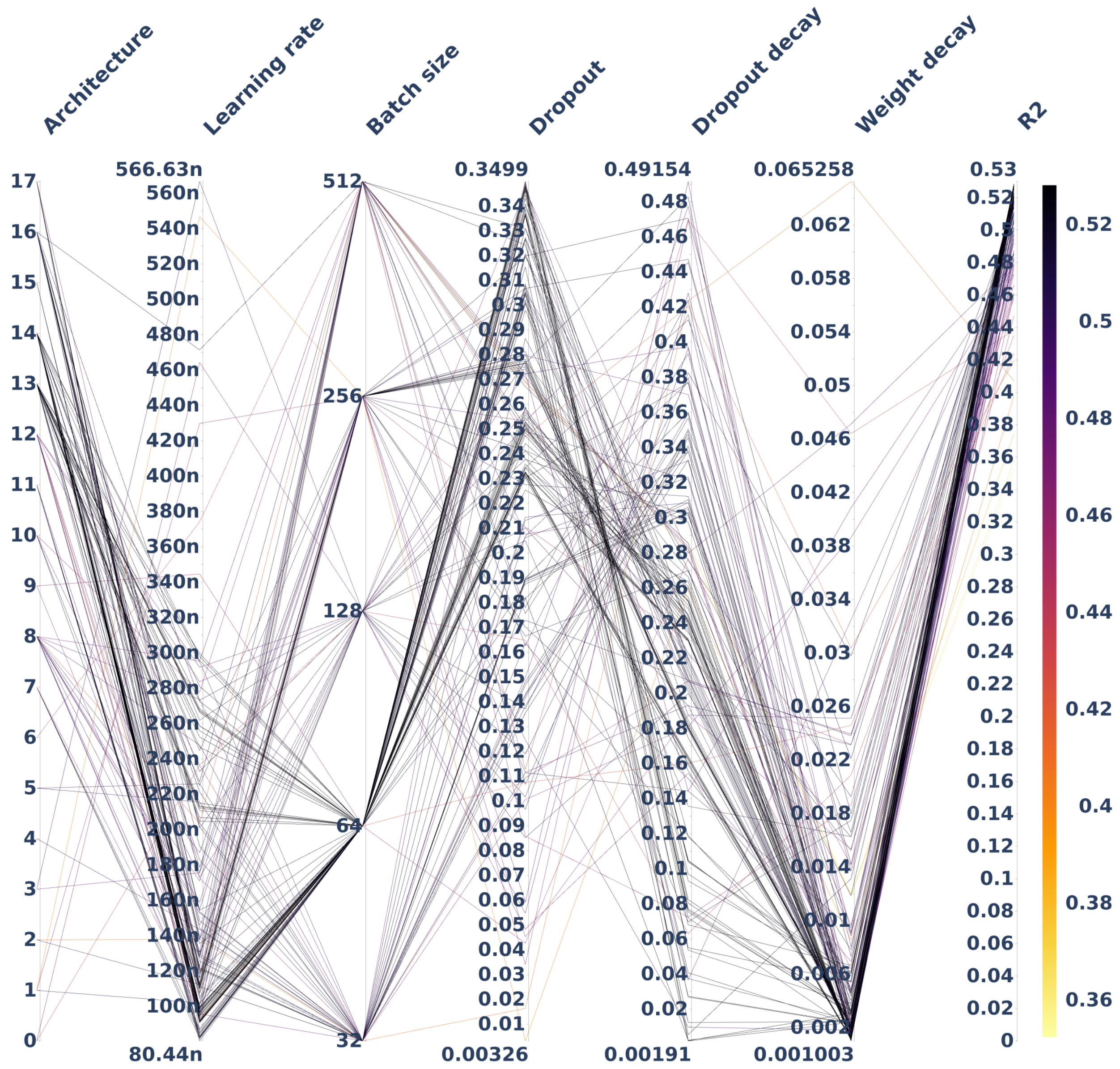


Figure S6: DNN hyperparameter space covered for FCFP. The covered architectures are presented by their identifier and their structures are available as an Excel file in the Supporting Information. The color scale varies with the $R^2$ from 0 (lighter color) to 0.53 (darker color).

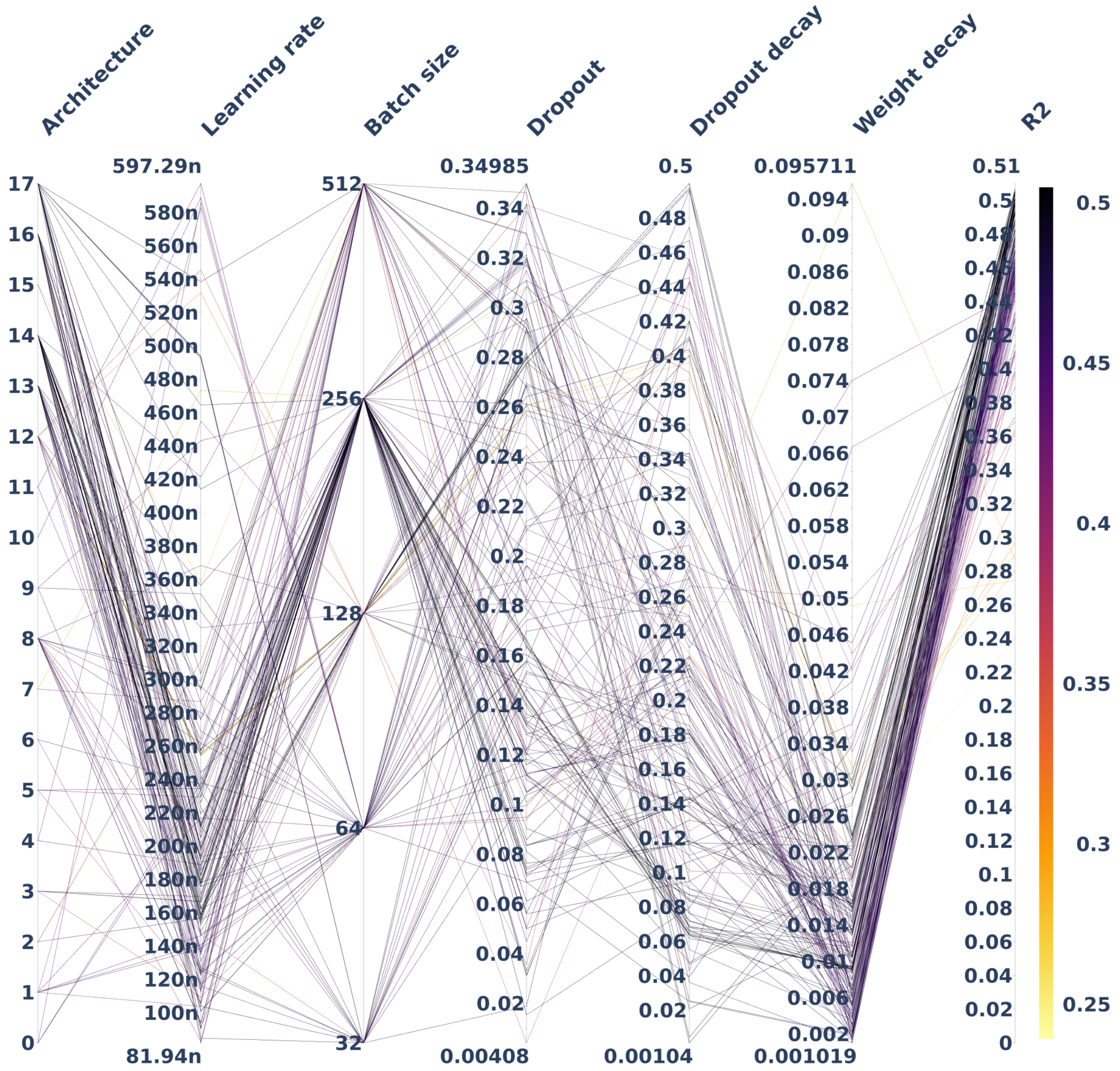


Figure S7: DNN hyperparameter space covered for E3FP. The covered architectures are presented by their identifier and their structures are available as an Excel file in the Supporting Information. The color scale varies with the $R^2$ from 0 (lighter color) to 0.51 (darker color).

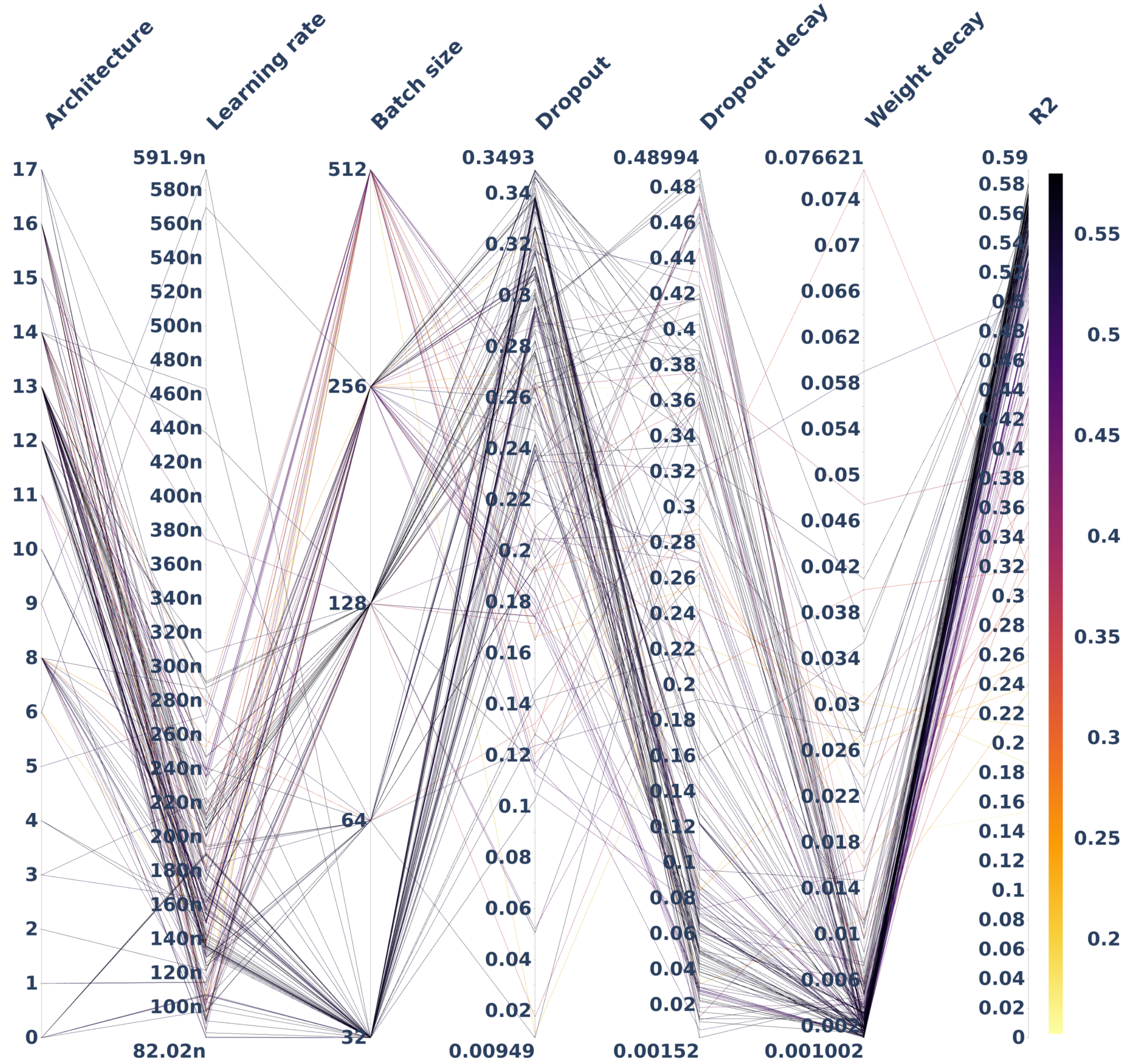


Figure S8: DNN hyperparameter space covered for *PLEC-4,096*. The covered architectures are presented by their identifier and their structures are available as an Excel file in the Supporting Information. The color scale varies with the $R^2$ from 0 (lighter color) to 0.59 (darker color).

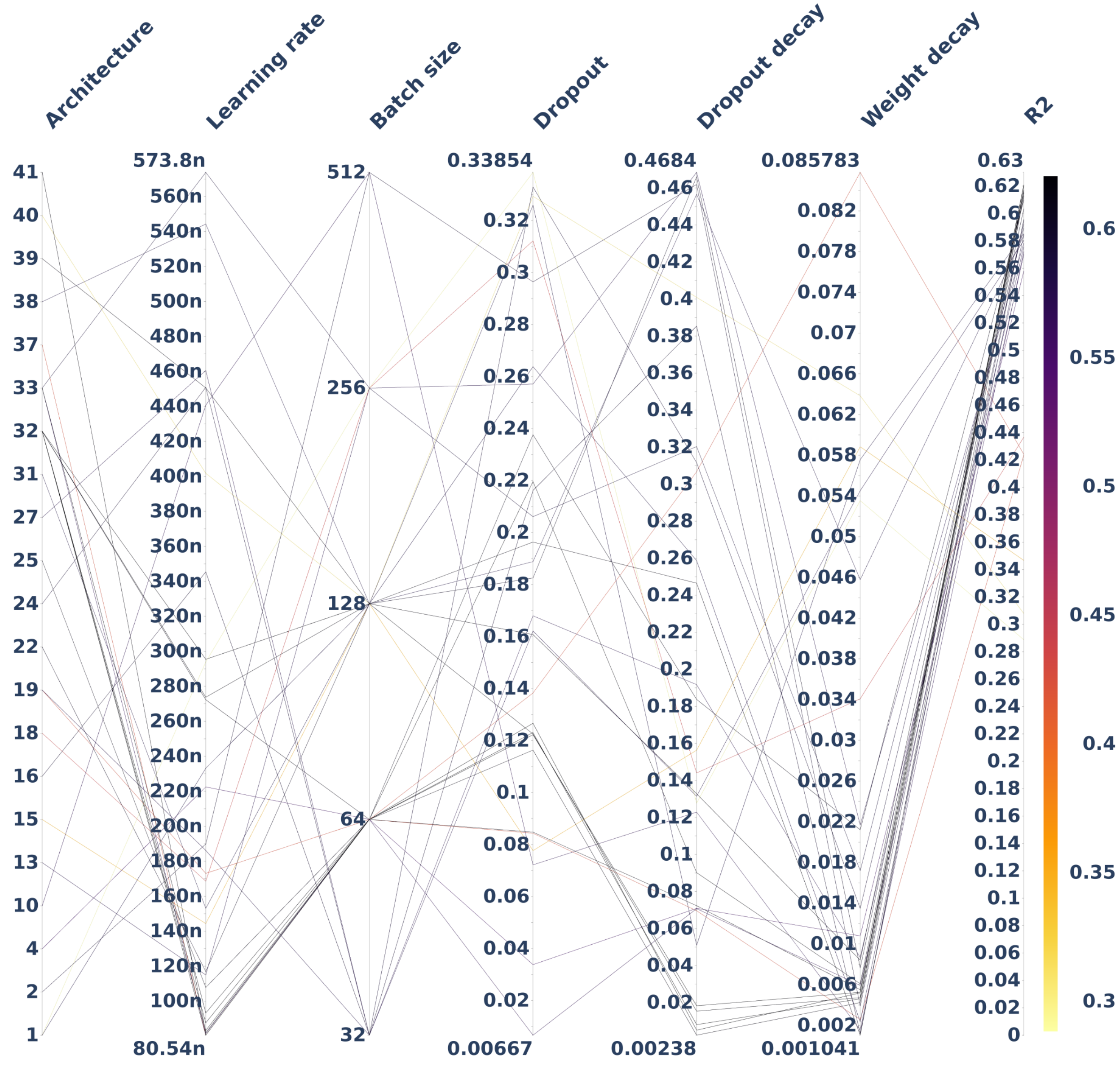


Figure S9: DNN hyperparameter space covered for *PLEC-16,384*. The covered architectures are presented by their identifier and their structures are available as an Excel file in the Supporting Information. The color scale varies with the $R^2$ from 0 (lighter color) to 0.63 (darker color).

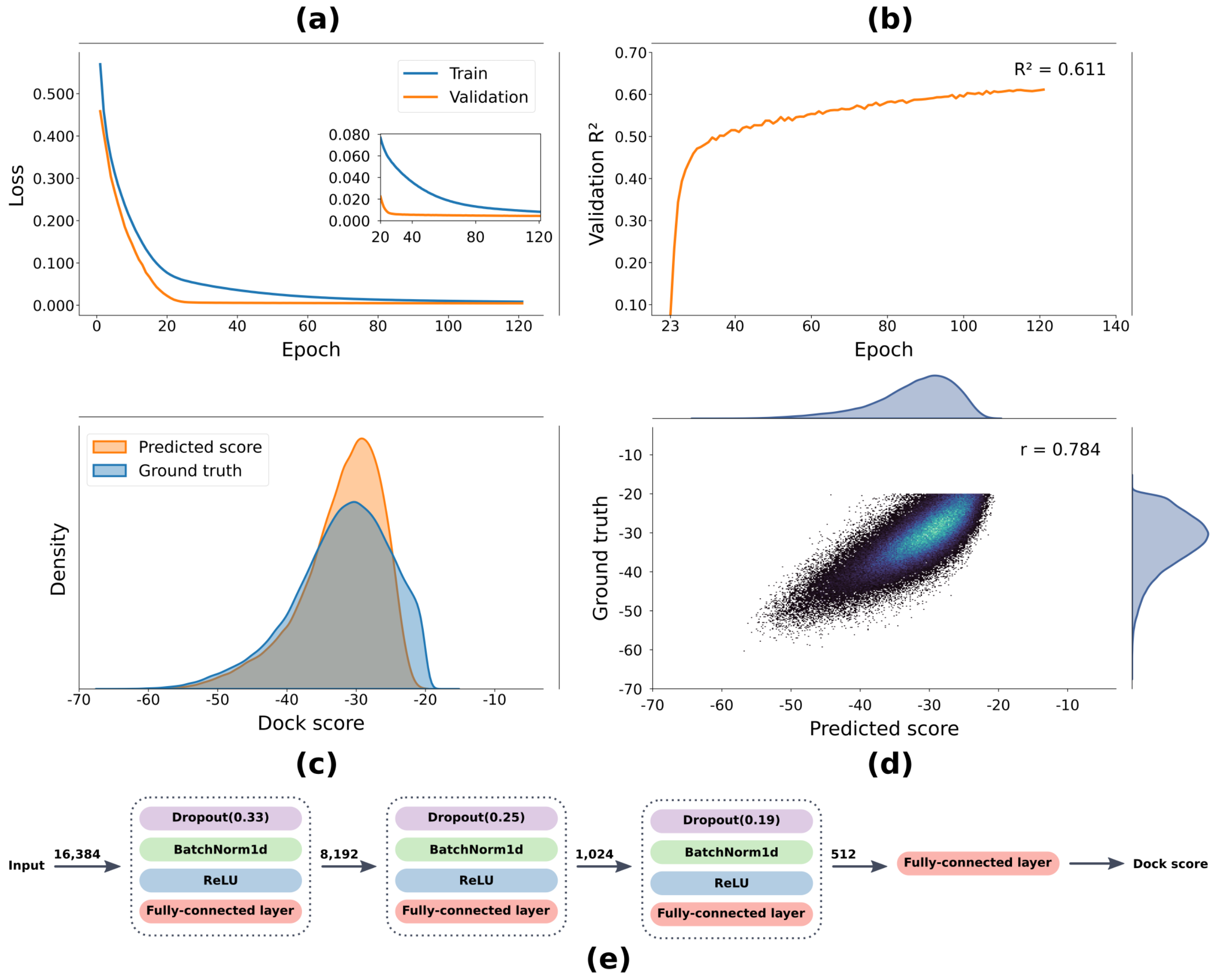


Figure S10: Model performance for the best architecture and combination of parameters for *EIFP-16384*. (a) The loss curve for the training and validation sets across the epochs. (b) The validation $R^2$ across the epochs. (c) The predicted and ground truth score distributions. (d) A scatter plot for the ground truth versus the predicted scores, as well as the Pearson correlation coefficient ($r$) between them. (e) The best architecture found for *EIFP-16384*.

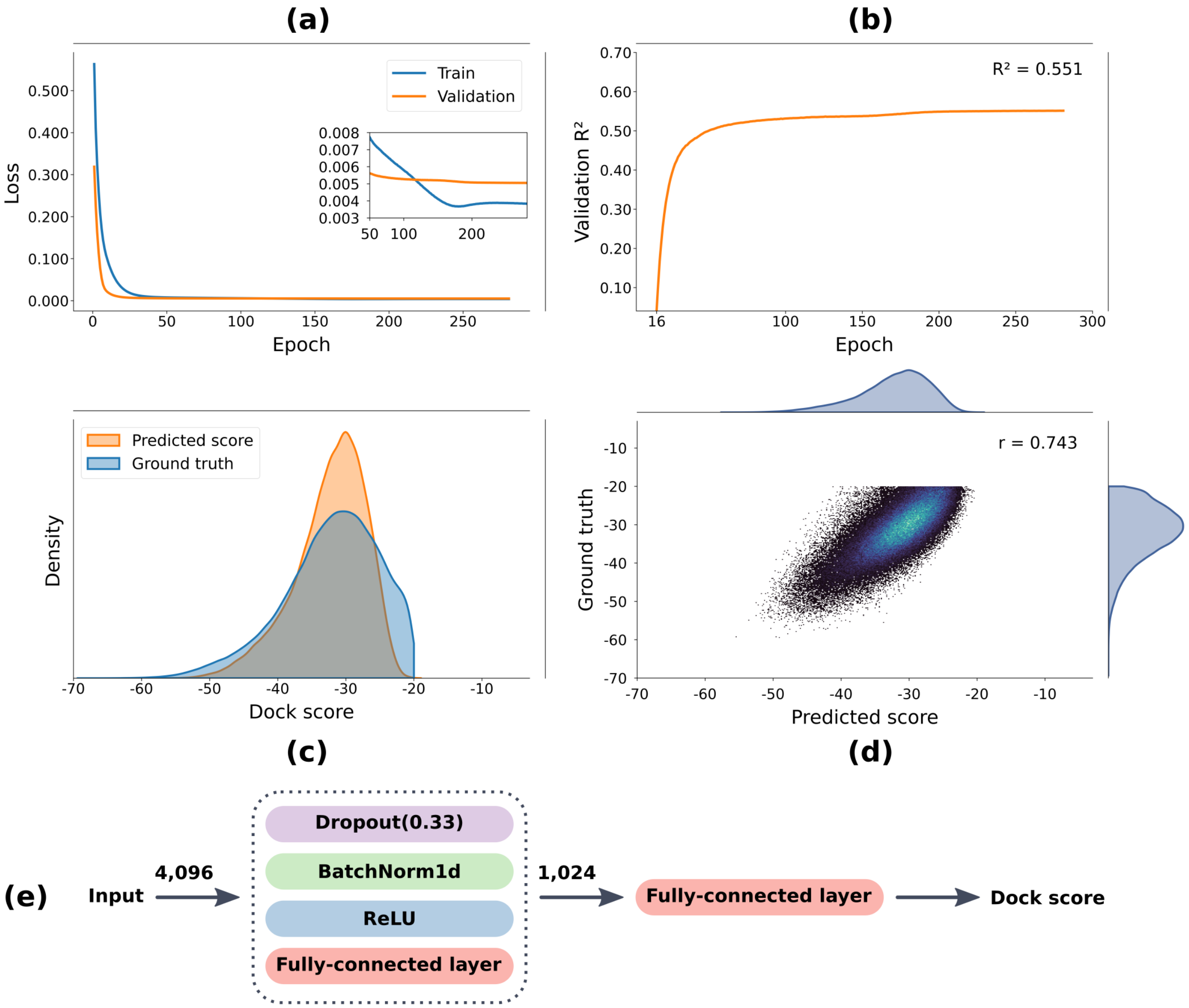


Figure S11: Model performance for the best architecture and combination of parameters for ECFP. (a) The loss curve for the training and validation sets across the epochs. (b) The validation $R^2$ across the epochs. (c) The predicted and ground truth score distributions. (d) A scatter plot for the ground truth versus the predicted scores, as well as the Pearson correlation coefficient ($r$) between them. (e) The best architecture found for ECFP.

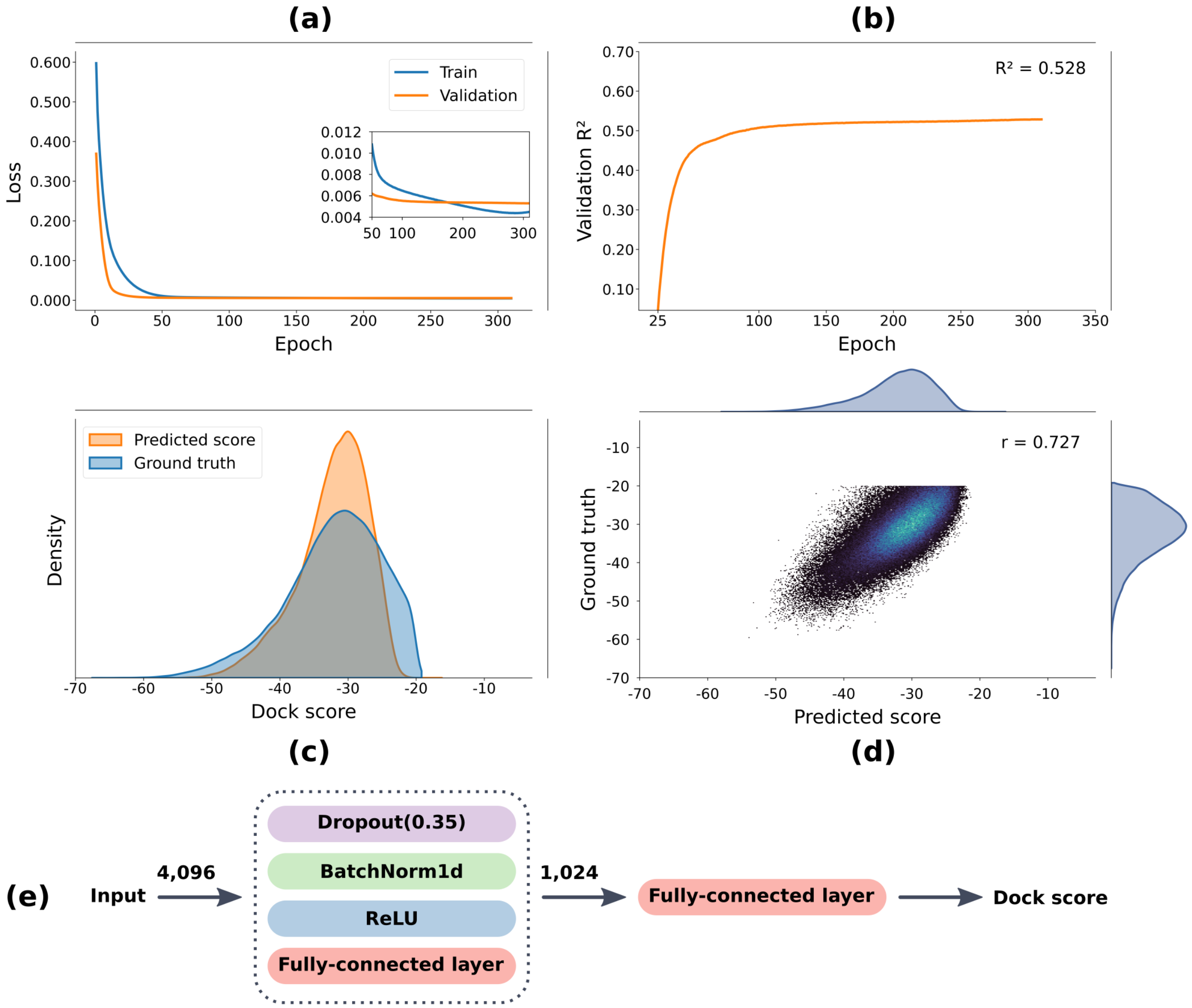


Figure S12: Model performance for the best architecture and combination of parameters for FCFP. (a) The loss curve for the training and validation sets across the epochs. (b) The validation $R^2$ across the epochs. (c) The predicted and ground truth score distributions. (d) A scatter plot for the ground truth versus the predicted scores, as well as the Pearson correlation coefficient ($r$) between them. (e) The best architecture found for FCFP.

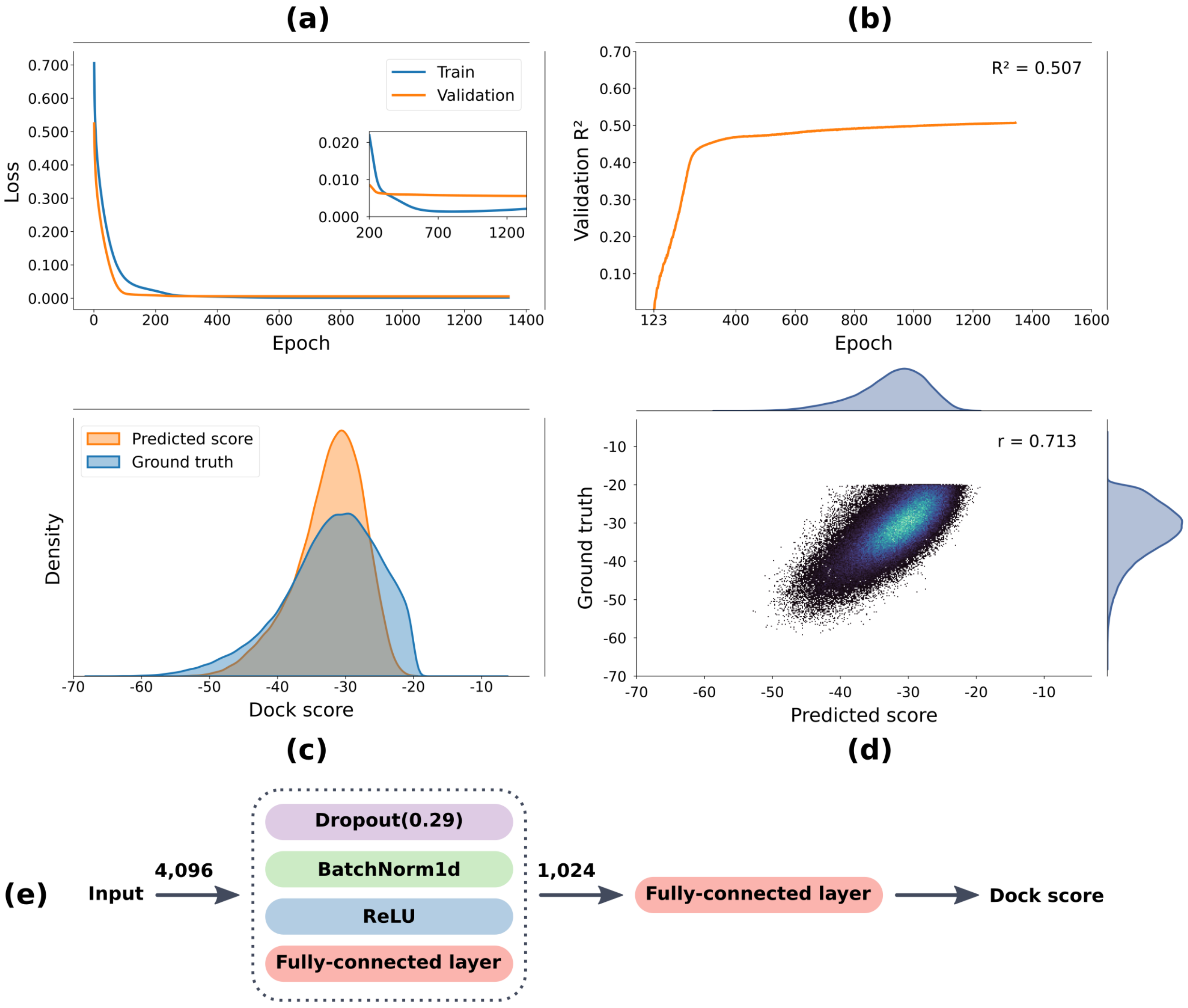


Figure S13: Model performance for the best architecture and combination of parameters for E3FP. (a) The loss curve for the training and validation sets across the epochs. (b) The validation $R^2$ across the epochs. (c) The predicted and ground truth score distributions. (d) A scatter plot for the ground truth versus the predicted scores, as well as the Pearson correlation coefficient ($r$) between them. (e) The best architecture found for E3FP.

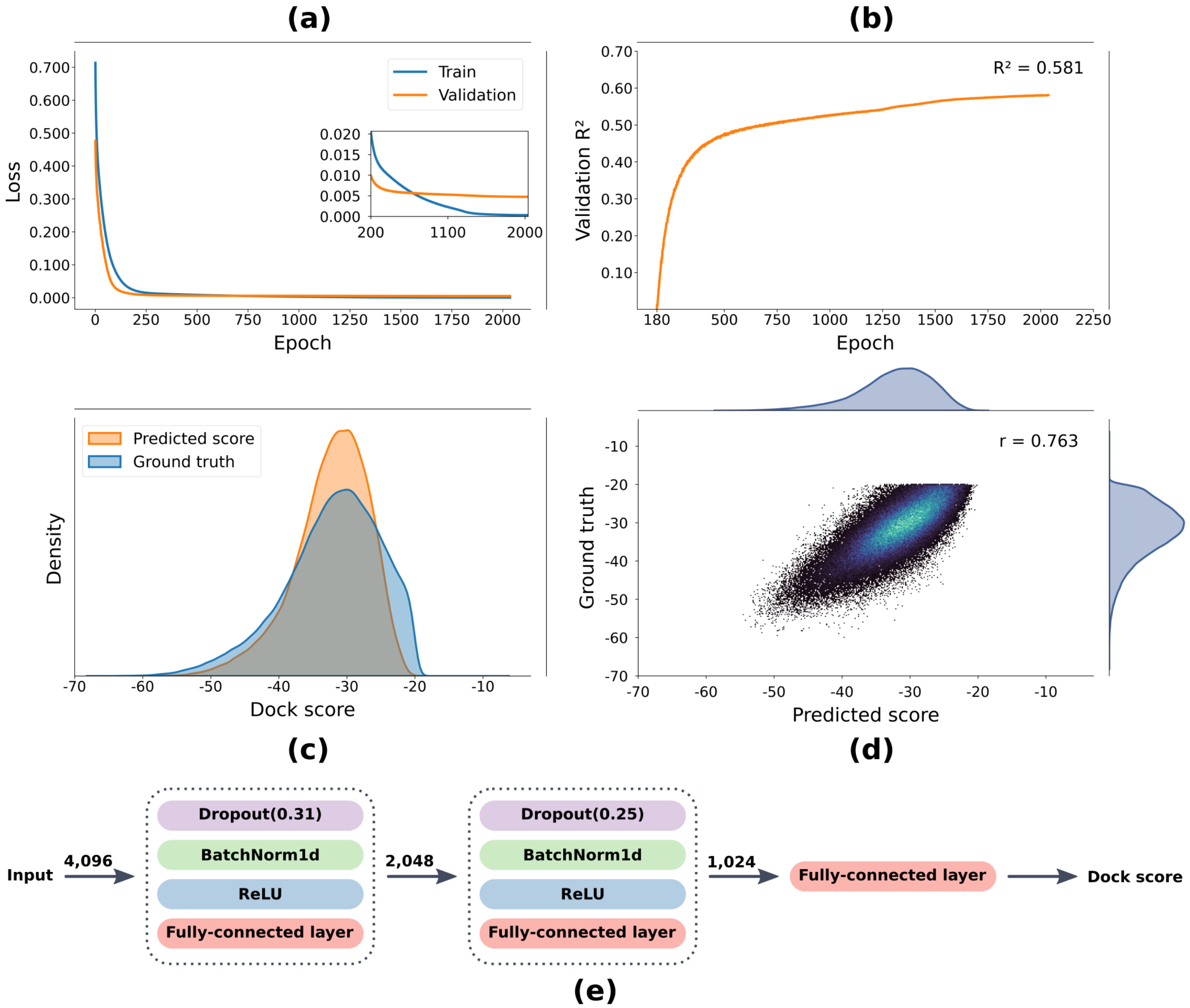


Figure S14: Model performance for the best architecture and combination of parameters for *PLEC-4096*. (a) The loss curve for the training and validation sets across the epochs. (b) The validation $R^2$ across the epochs. (c) The predicted and ground truth score distributions. (d) A scatter plot for the ground truth versus the predicted scores, as well as the Pearson correlation coefficient ($r$) between them. (e) The best architecture found for *PLEC-4096*.

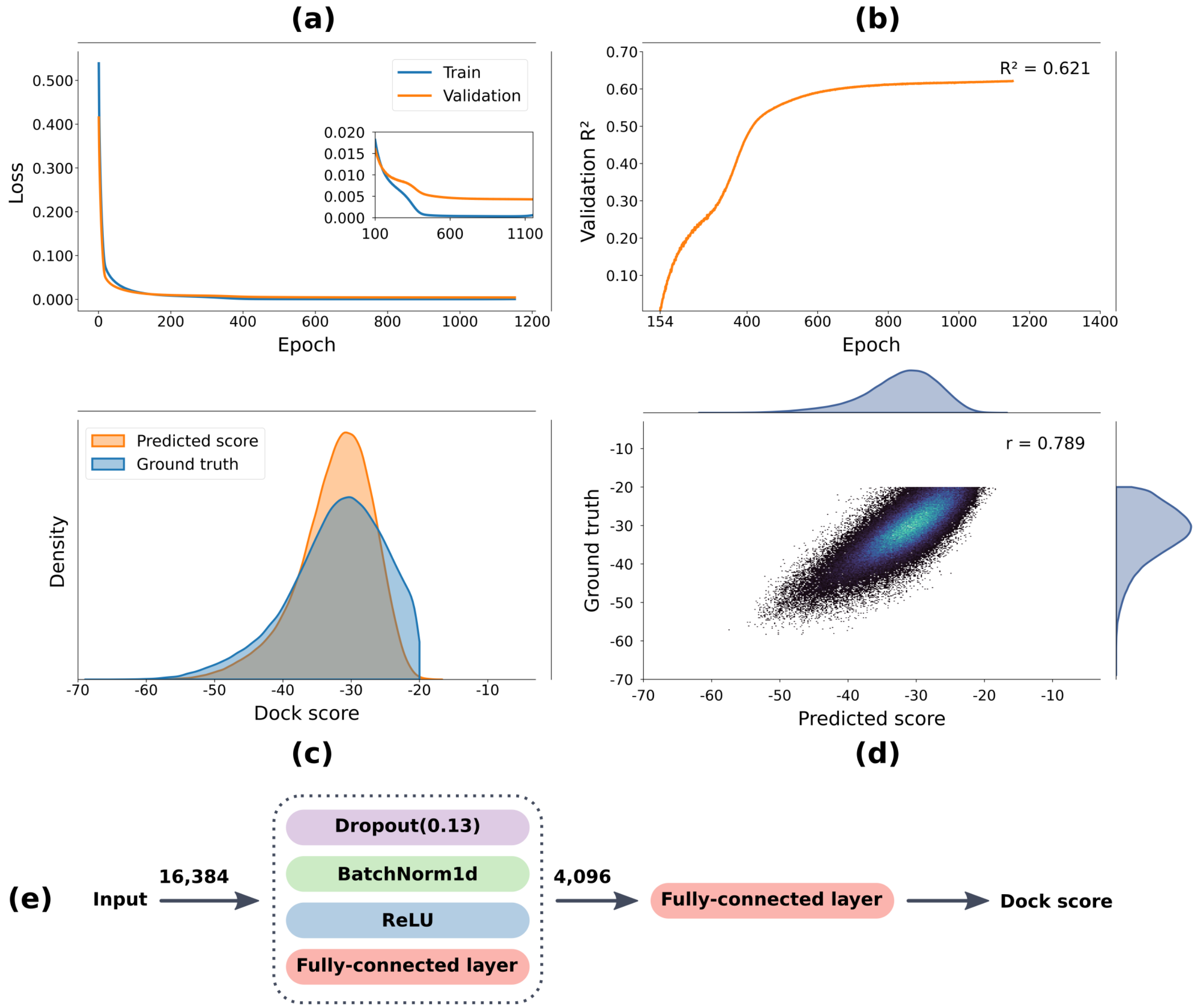


Figure S15: Model performance for the best architecture and combination of parameters for *PLEC-16384*. (a) The loss curve for the training and validation sets across the epochs. (b) The validation $R^2$ across the epochs. (c) The predicted and ground truth score distributions. (d) A scatter plot for the ground truth versus the predicted scores, as well as the Pearson correlation coefficient ($r$) between them. (e) The best architecture found for *PLEC-16384*.

Table S4: 5-fold cross-validation results for the DNN strategy.

| Fingerprint | Mean | Std |
|---|---|---|
| *PLEC-16,384* | **0.609** | 0.003 |
| *EIFP-16,384* | 0.603 | 0.002 |
| *EIFP-4,096* | 0.597 | 0.014 |
| *PLEC-4,096* | 0.536 | 0.004 |
| ECFP | 0.536 | 0.012 |
| FCFP | 0.518 | 0.003 |
| E3FP | 0.466 | 0.012 |

Table S5: 5-fold cross-validation results for the XGBoost strategy.

| Fingerprint | Mean | Std |
|---|---|---|
| *EIFP-16,384* | **0.545** | 0.001 |
| *EIFP-4,096* | 0.536 | 0.001 |
| *PLEC-16,384* | 0.486 | 0.002 |
| ECFP | 0.477 | 0.002 |
| E3FP | 0.459 | 0.001 |
| FCFP | 0.458 | 0.002 |
| *PLEC-4,096* | 0.440 | 0.002 |

Table S6: 5-fold cross-validation results for the random forest strategy.

| Fingerprint | Mean | Std |
|---|---|---|
| *EIFP-16,384* | **0.528** | 0.001 |
| *EIFP-4,096* | 0.518 | 0.001 |
| ECFP | 0.489 | 0.004 |
| E3FP | 0.471 | 0.002 |
| FCFP | 0.466 | 0.002 |
| *PLEC-16,384* | 0.459 | 0.001 |
| *PLEC-4,096* | 0.376 | 0.002 |

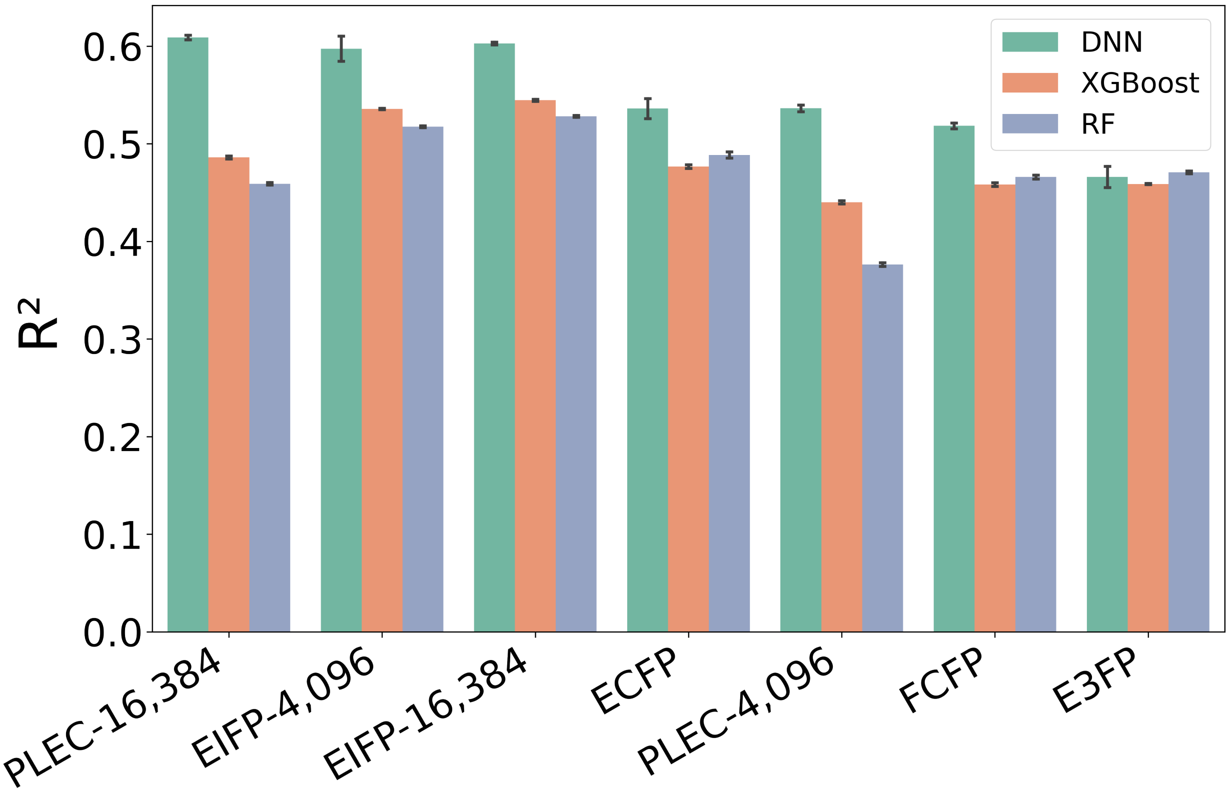


Figure S16: 5-fold cross-validation for all FPs and ML methods. To guarantee reproducibility, we used an initial seed (54,343) to generate five random integer values that were used as seeds to randomly define the hold-out set. FPs were sorted based on DNN results in descending order.

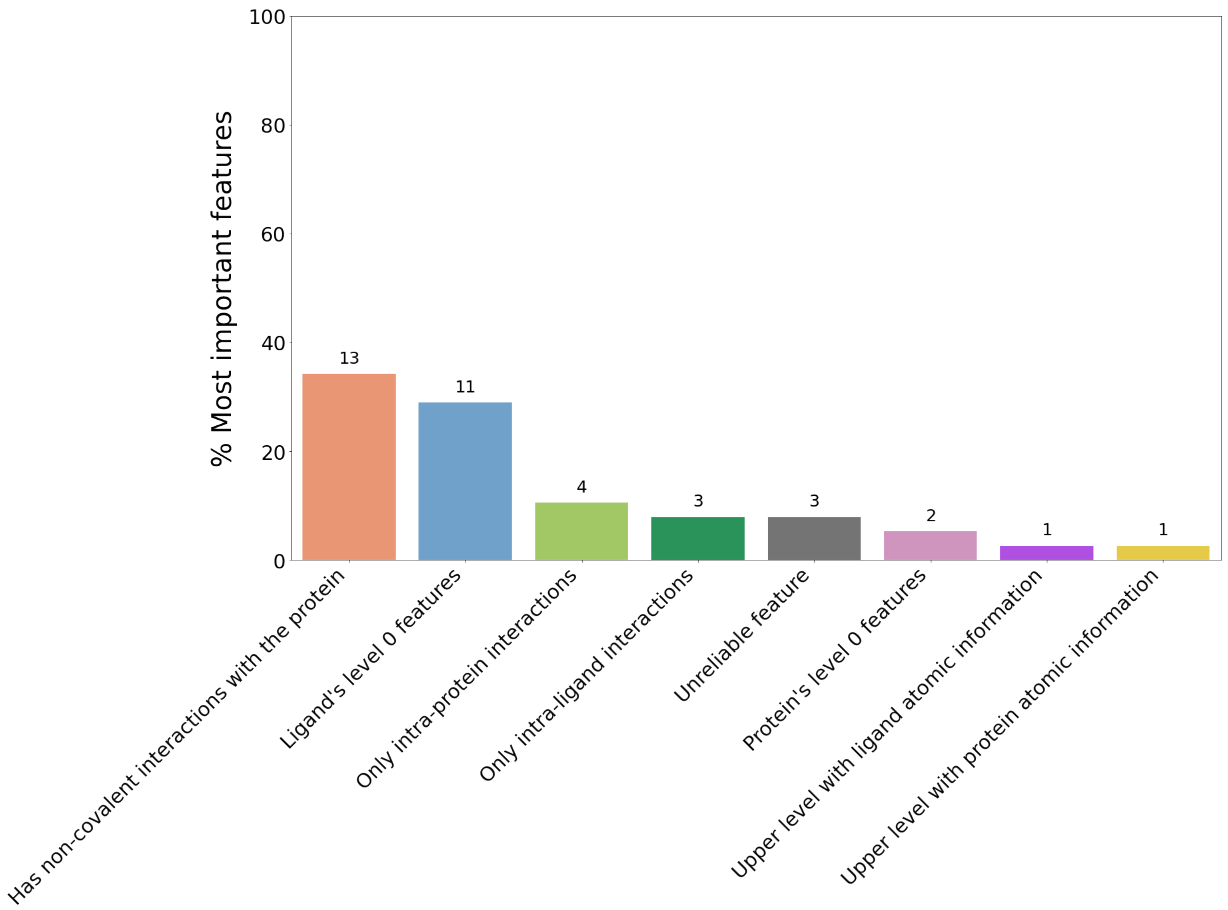


Figure S17: Percentage of each class attributed to the 38 most important features. Numbers above the bars represent the total number of features having a given feature class.

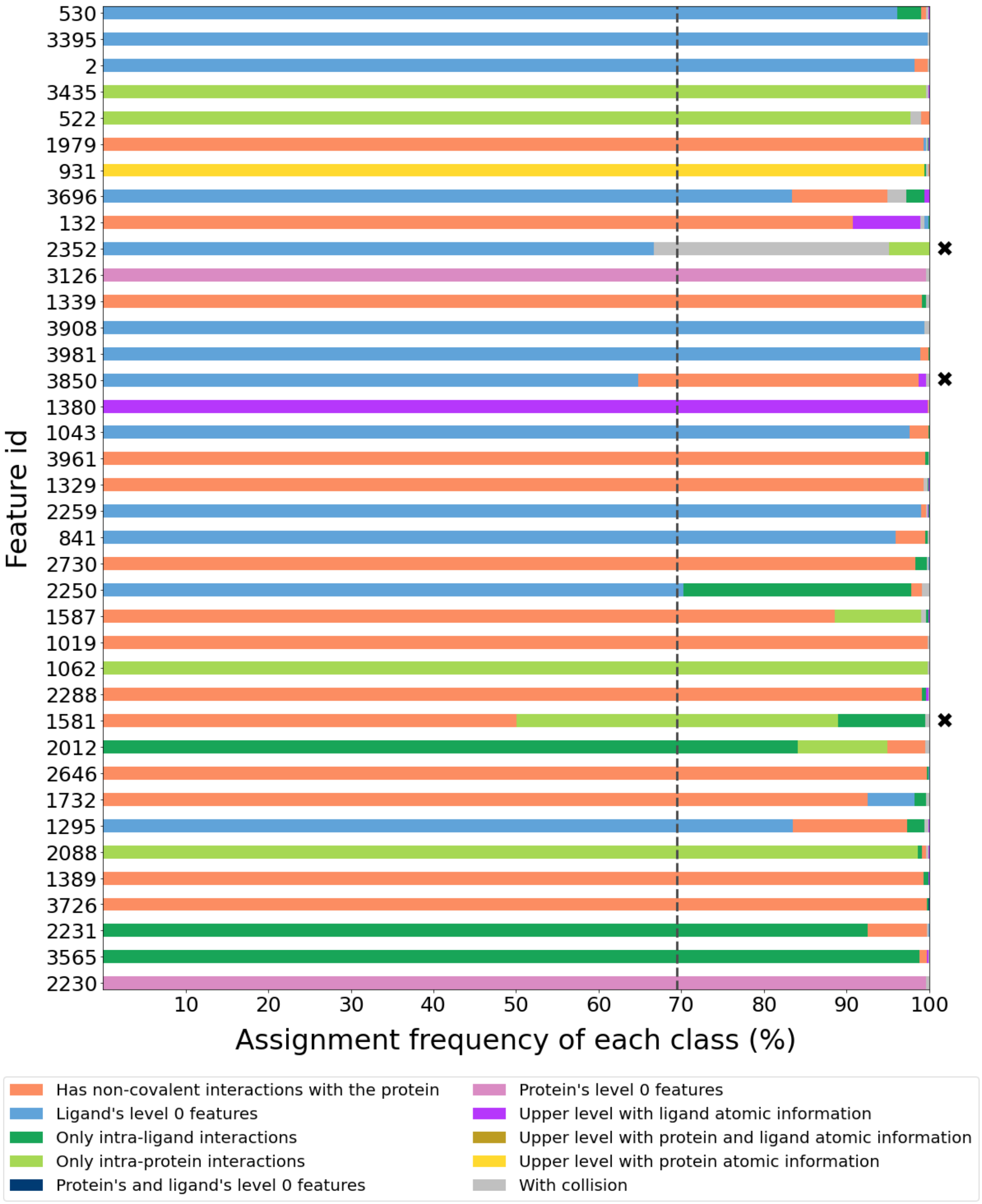


Figure S18: Percentage of all classes assigned to the 38 most important features. The most common class in a given feature was only assigned to the feature whether its percentage was greater than or equal to the threshold (69.49%, chosen by z-scores; dashed line). Features are sorted in descending order of the importance score and cross markers point out features considered unreliable.

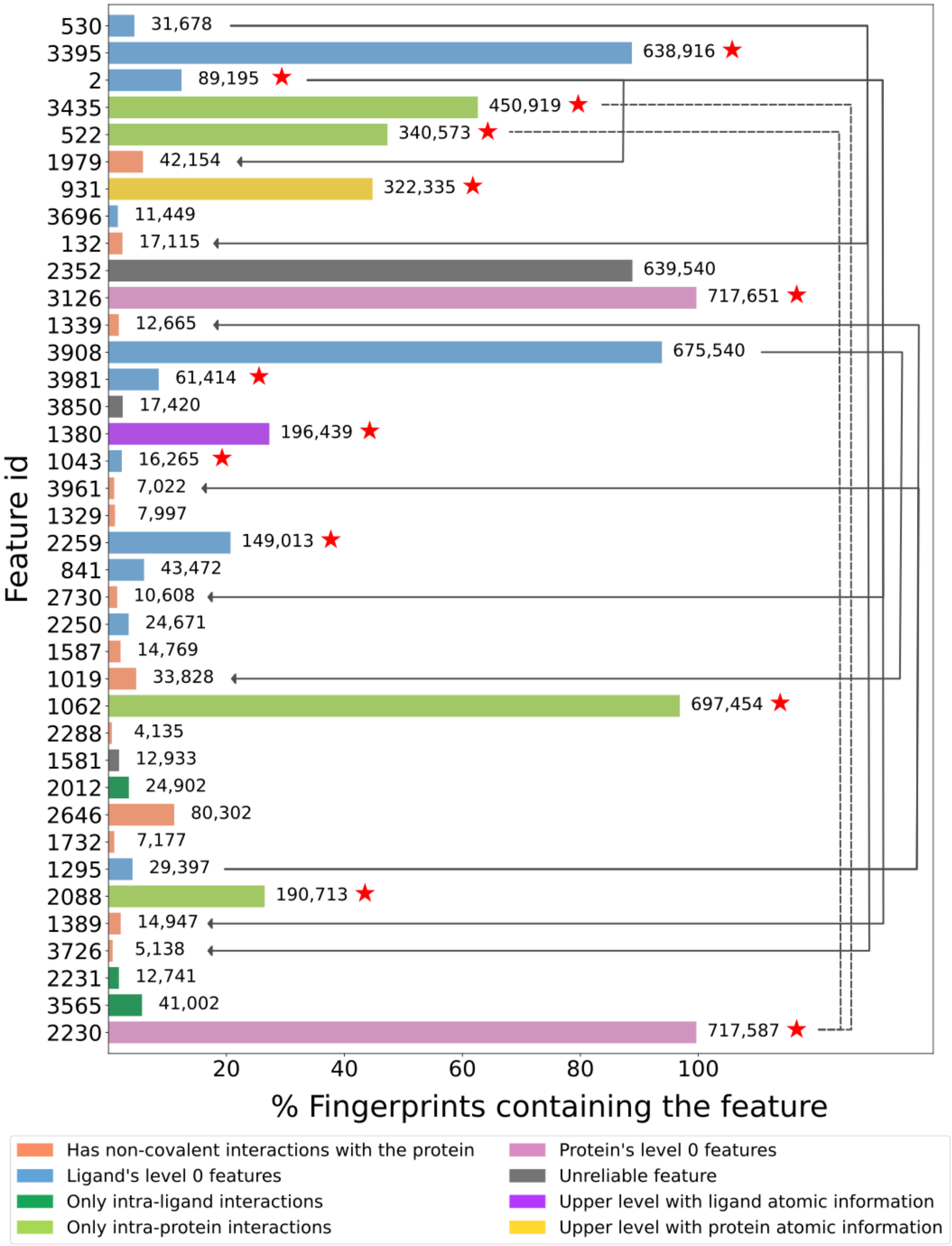


Figure S19: Number of ligands containing the most important features. Features are sorted in descending order of the importance score and red star markers point out features that are highly correlated to another feature. Dashed lines connect features that are correlated to each other, while directed solid lines connect ligands' atoms to the interactions established by them.

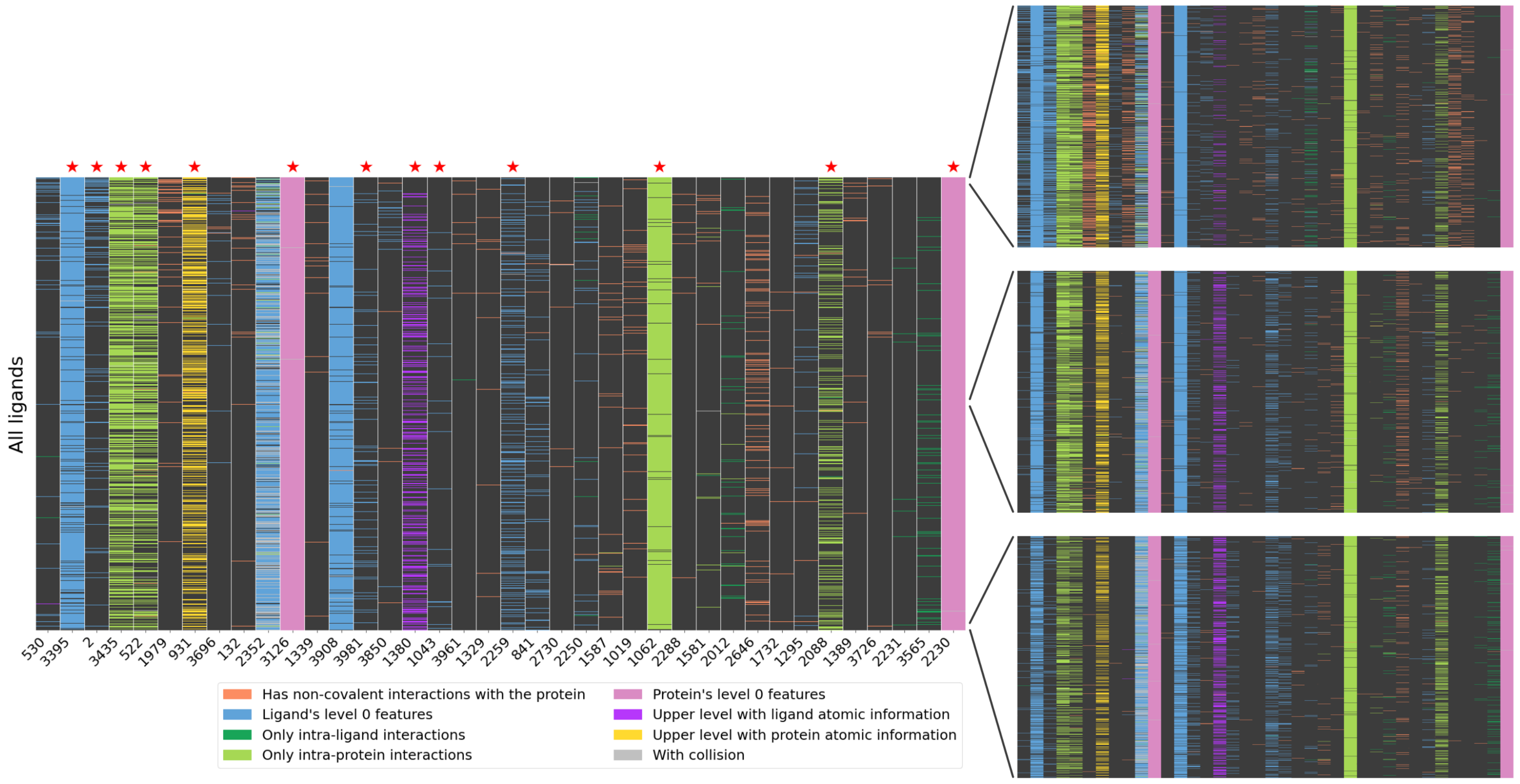


Figure S20: Heat map for the 38 most important features across all ligands in the training set. Zoomed-in heat maps highlight the presence of a feature among the best, median, and worst ligands. Features are sorted in descending order of the importance score and red star markers point out features that are highly correlated to another feature.

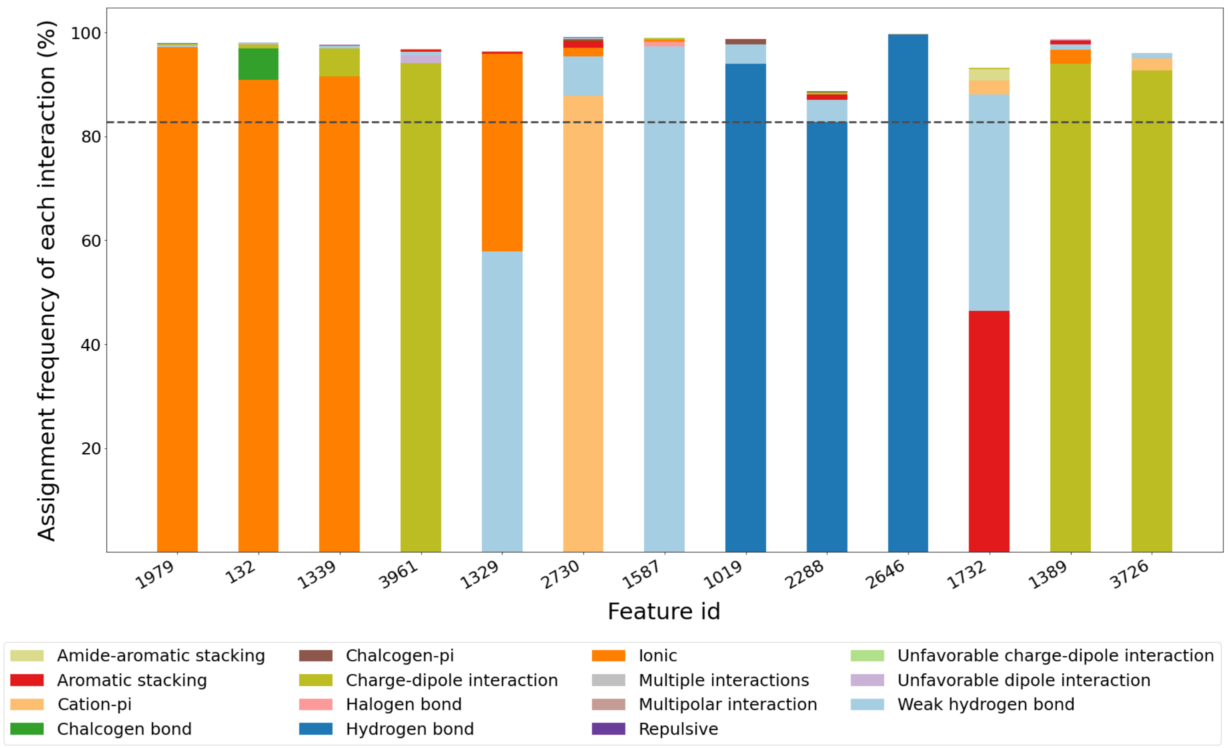


Figure S21: Percentage of all interactions assigned to the thirteen most important features comprising protein-ligand interactions. The most common interaction in a given feature was only assigned to the feature whether its percentage was greater than or equal to the threshold (82.78%, chosen by z-scores; dashed line). Features are sorted in descending order of the importance score.

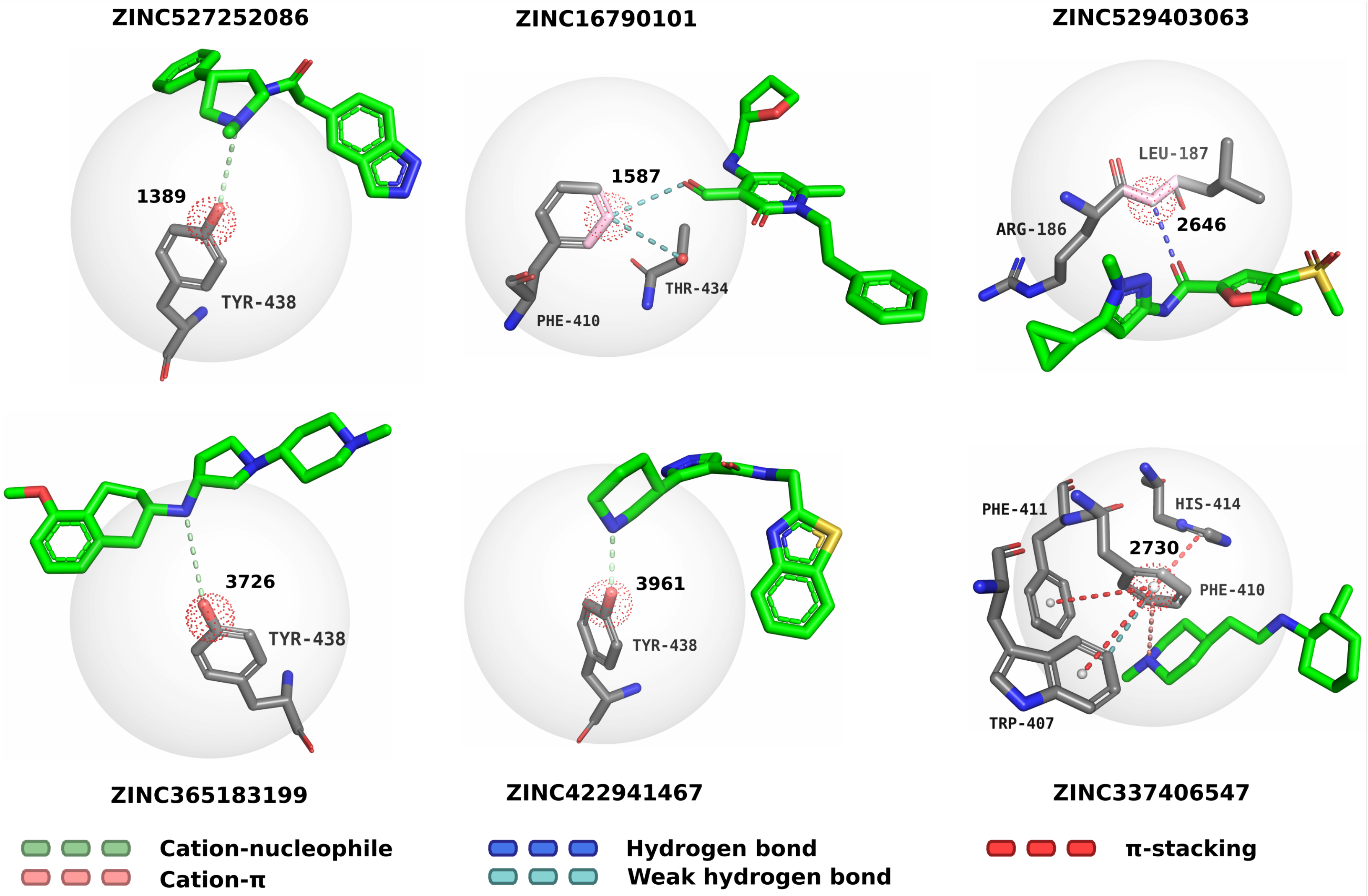


Figure S22: Examples of six out of the thirteen most important features comprising protein-ligand interactions. The feature is represented by the bigger sphere, while the dotted red sphere highlights its centroid. Covalent bonds comprising the feature are depicted as pink sticks. Features are sorted in descending order of the importance score.

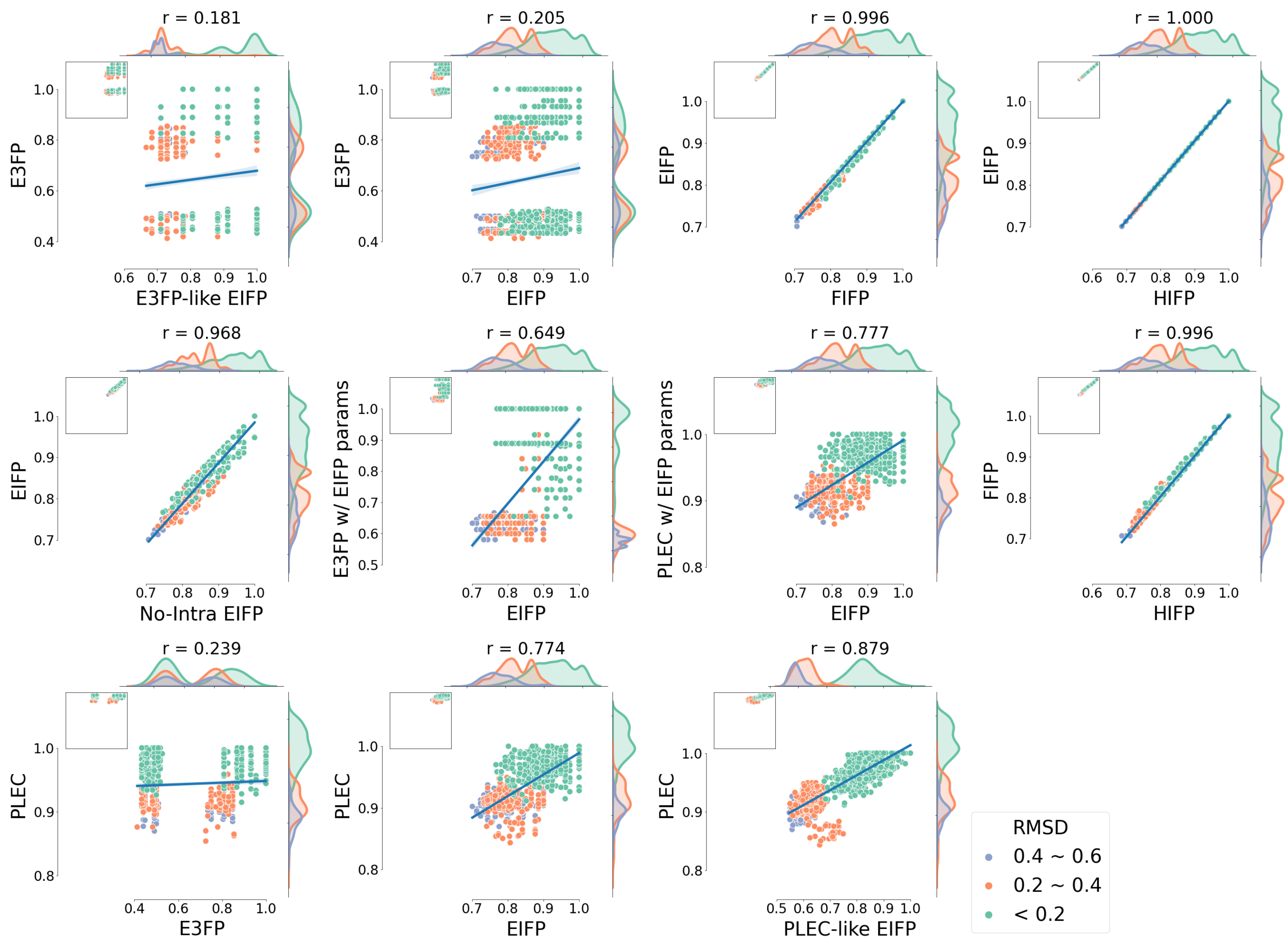


Figure S23: Correlation between fingerprint pairs using the data set "*Automatically generated conformers (0.4 Å)*". The correlation data points were colored according to the RMSD between the pairs of conformers.

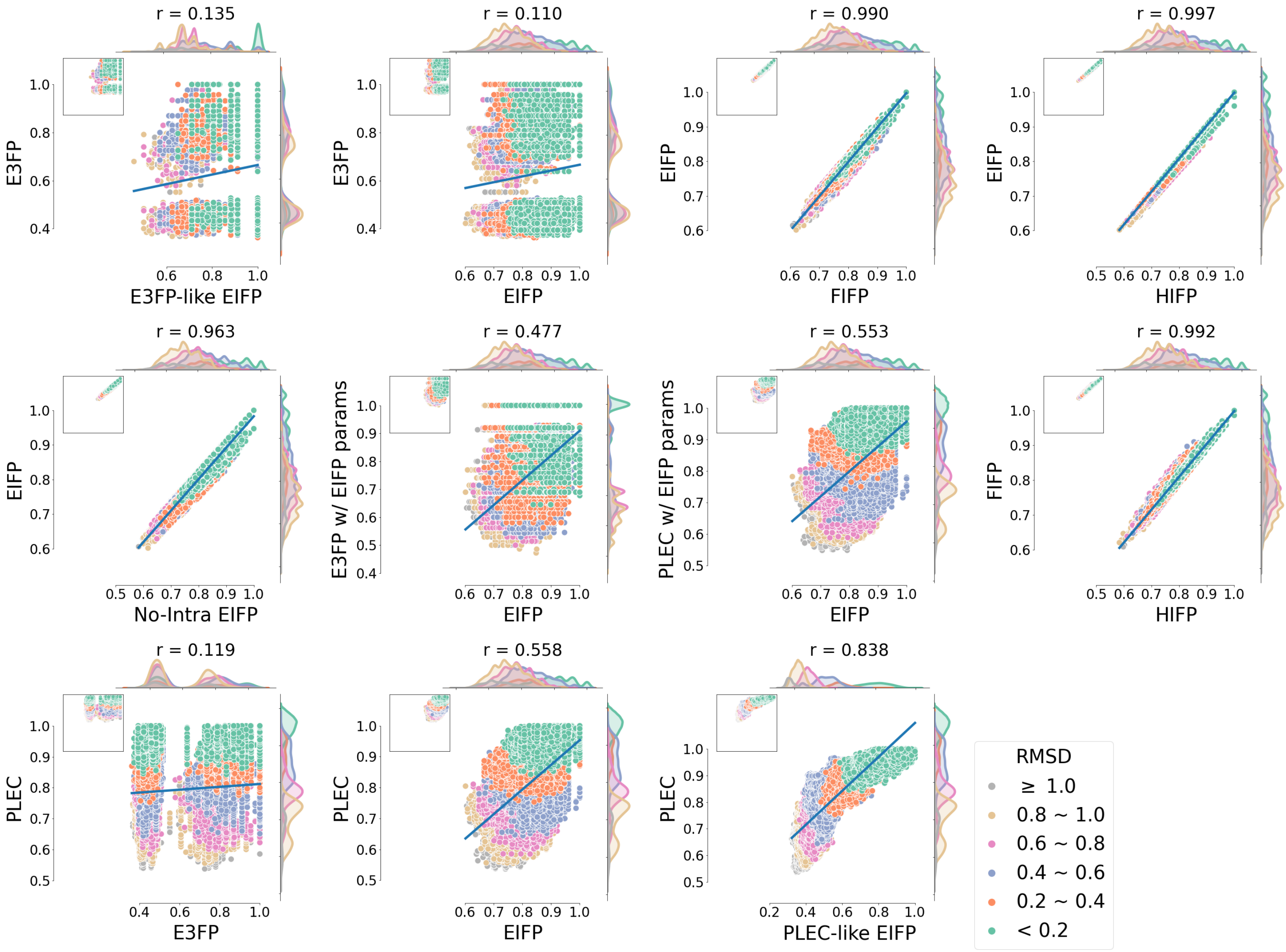


Figure S24: Correlation between fingerprint pairs using the data set "*Automatically generated conformers (0.8 Å)*". The correlation data points were colored according to the RMSD between the pairs of conformers.

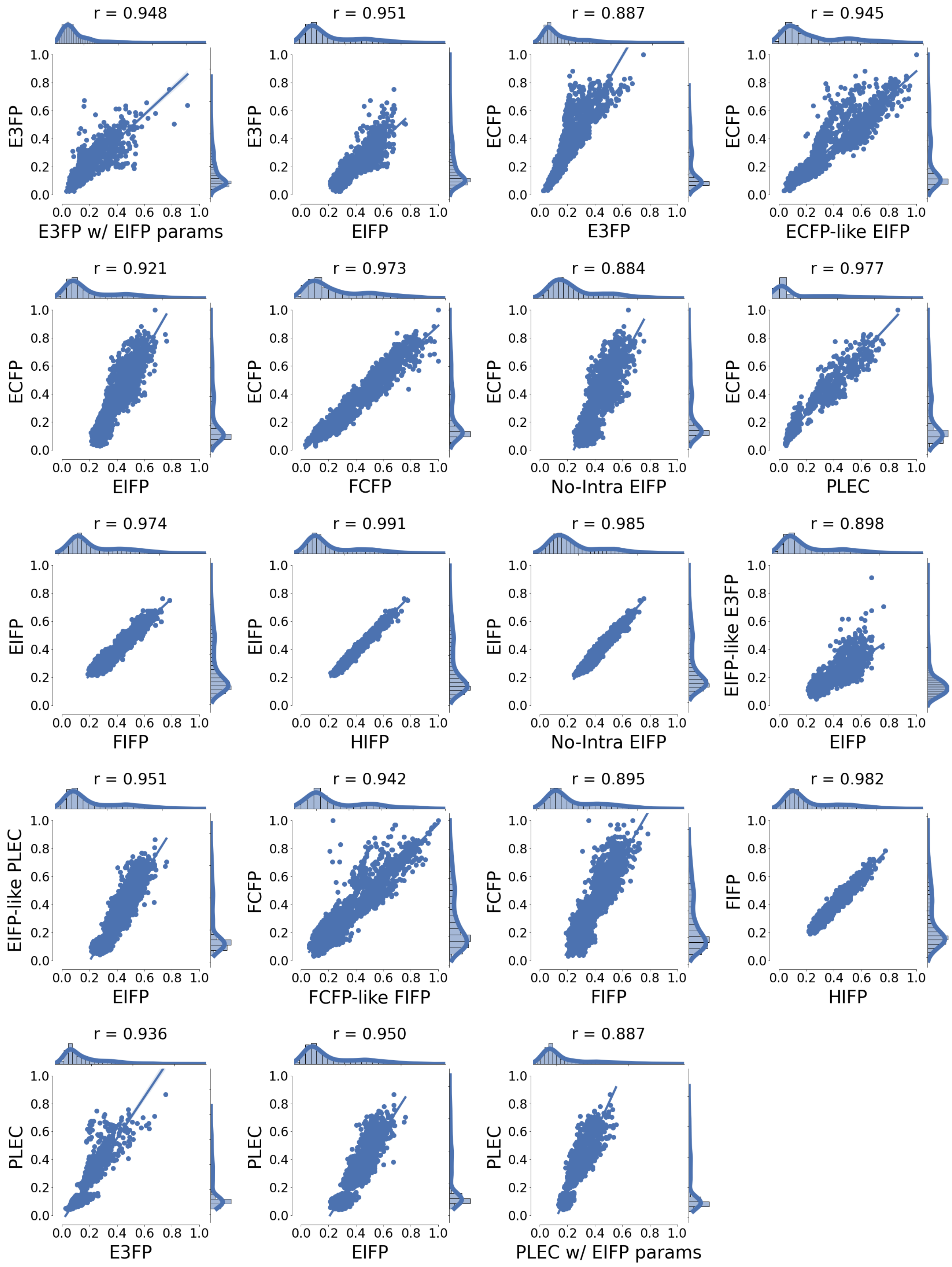


Figure S25: Correlation between fingerprint pairs using the data set "*CDK2 complexes from Schonbrunn's work*".

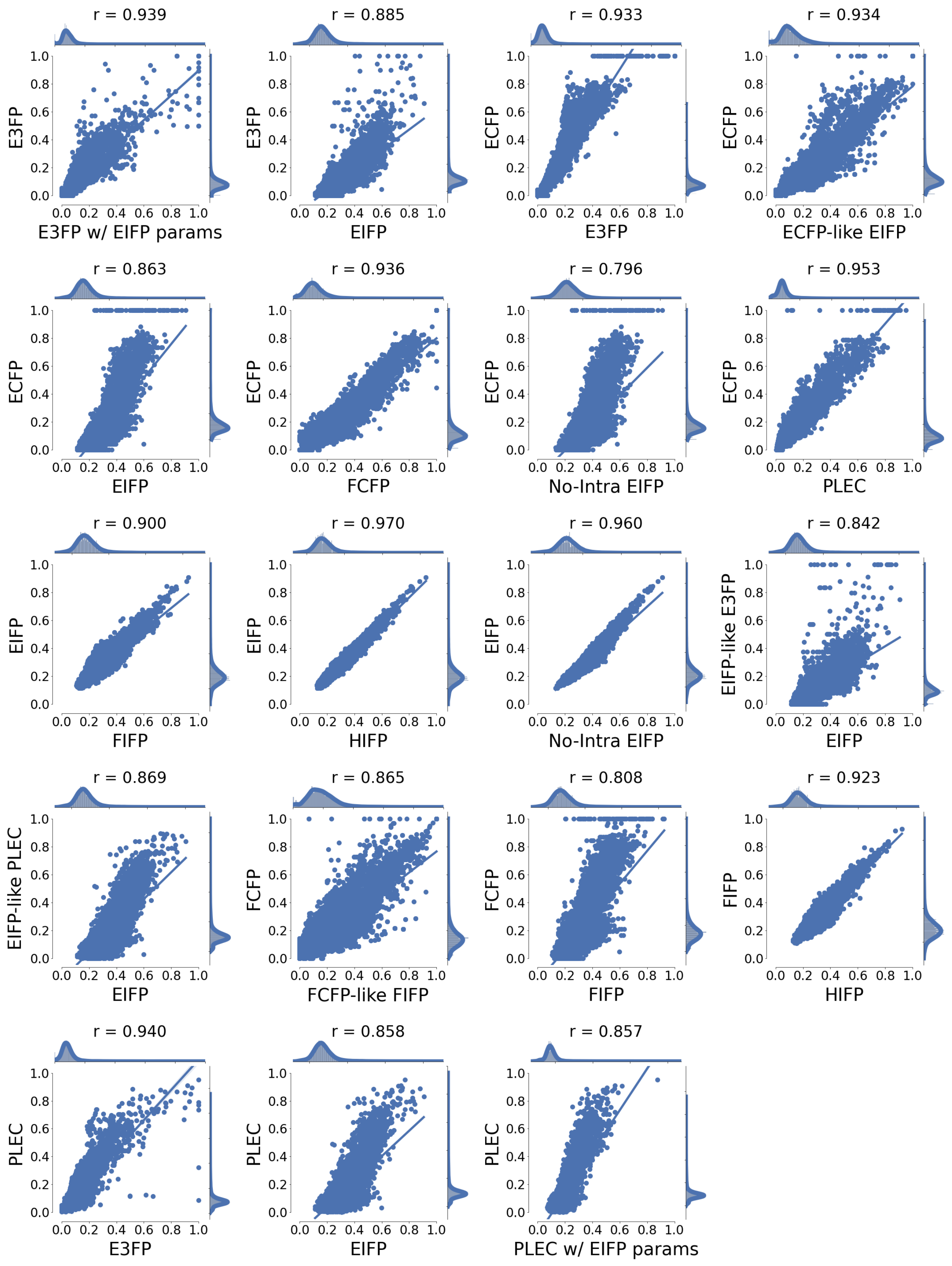


Figure S26: Correlation between fingerprint pairs using the data set "*Extended CDK2 set*".

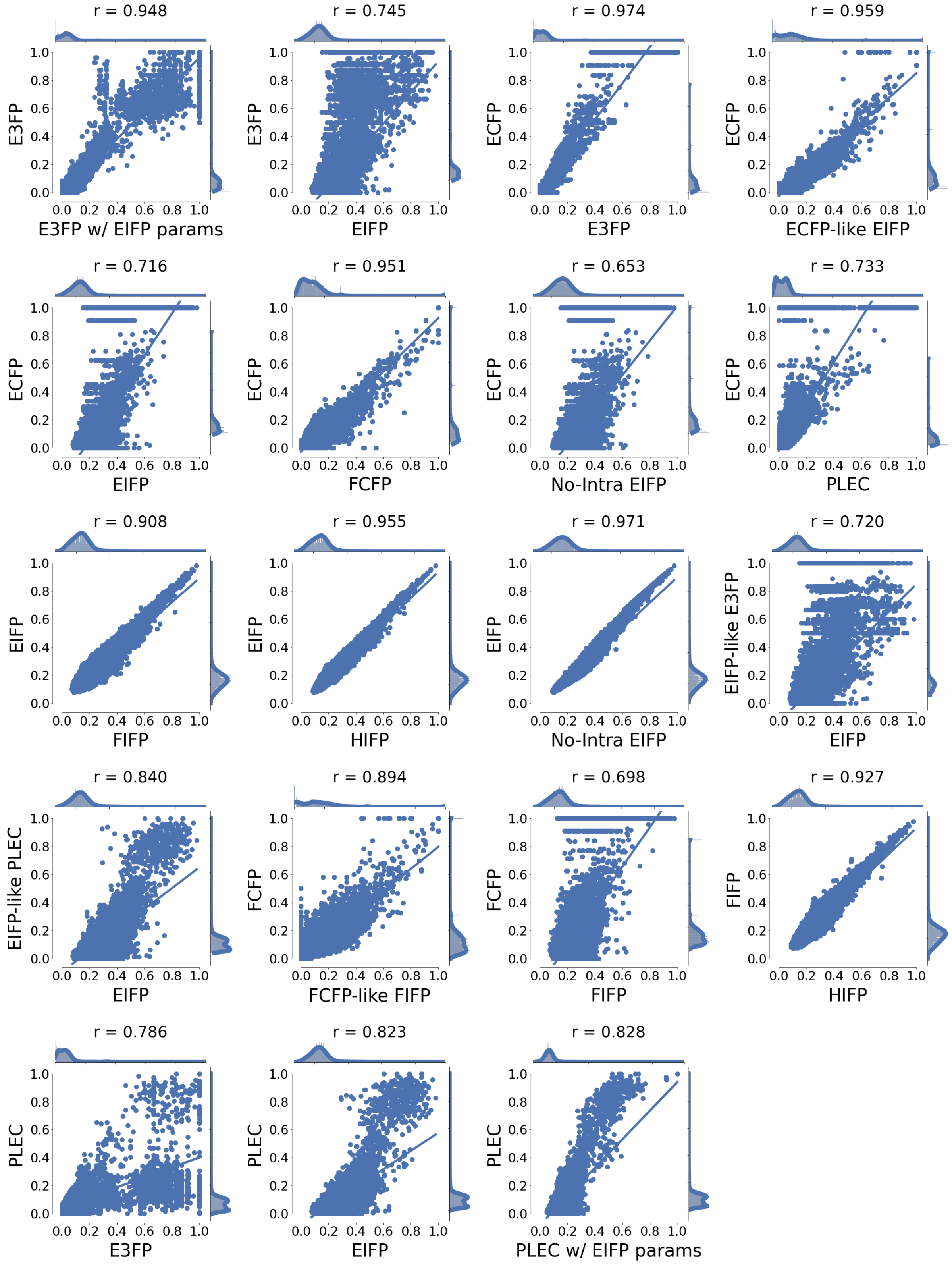


Figure S27: Correlation between fingerprint pairs using the data set "*Other non-CDK2 kinases*".

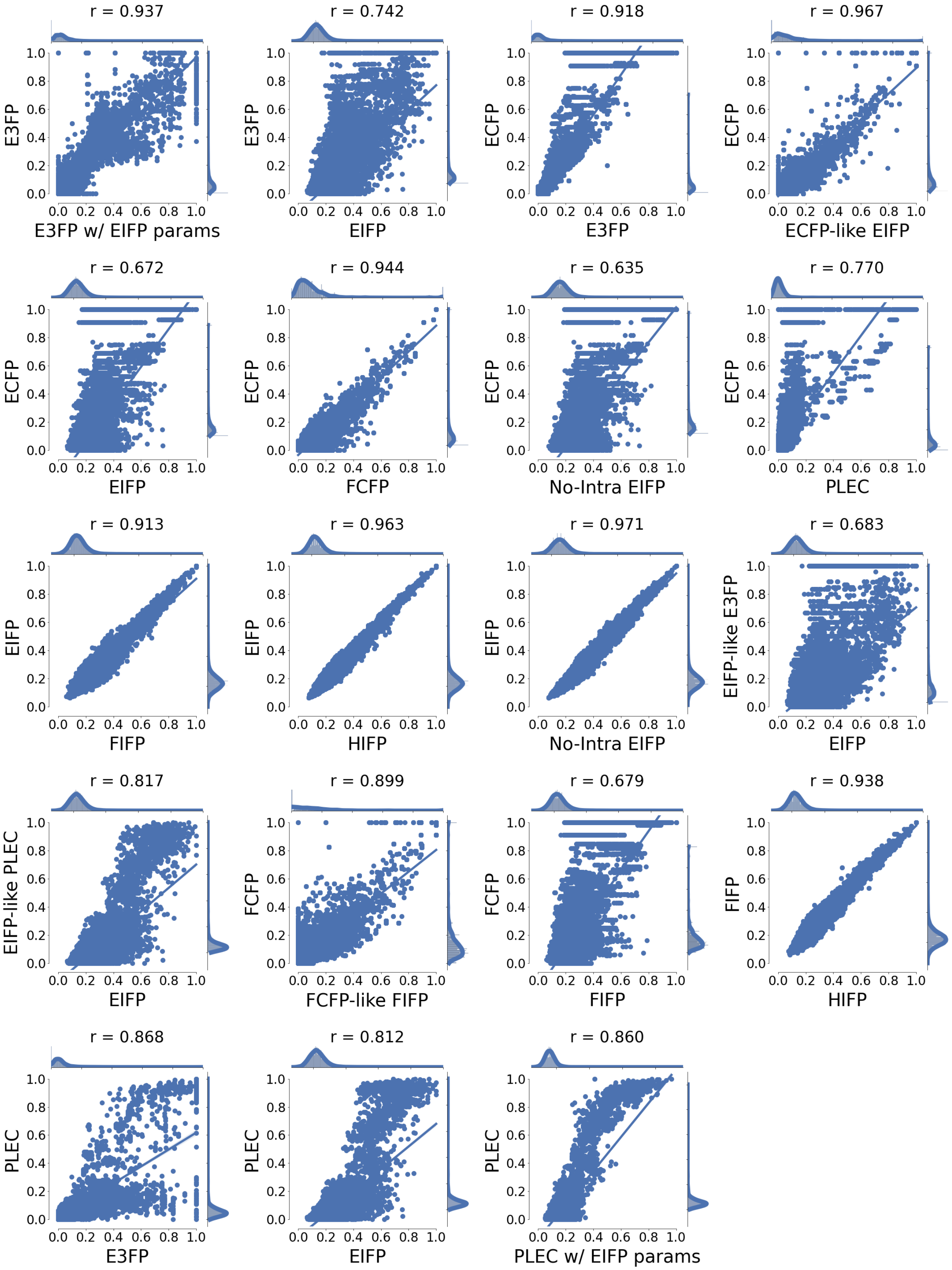


Figure S28: Correlation between fingerprint pairs using the data set "*Diverse proteins*".

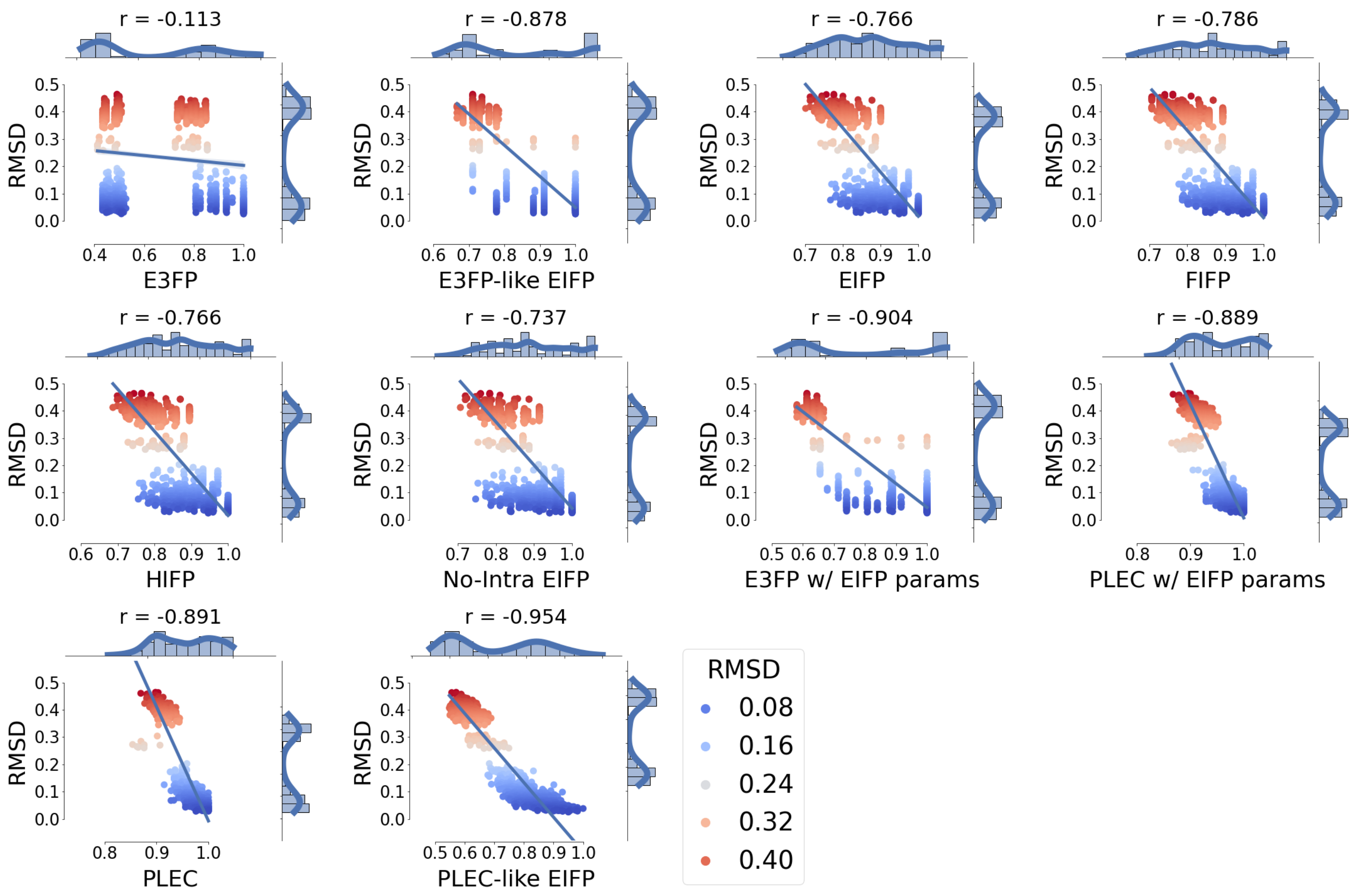


Figure S29: Correlation between the conformers' RMSD and the similarity of fingerprint pairs using the dataset "*Automatically generated conformers (0.4 Å)*".

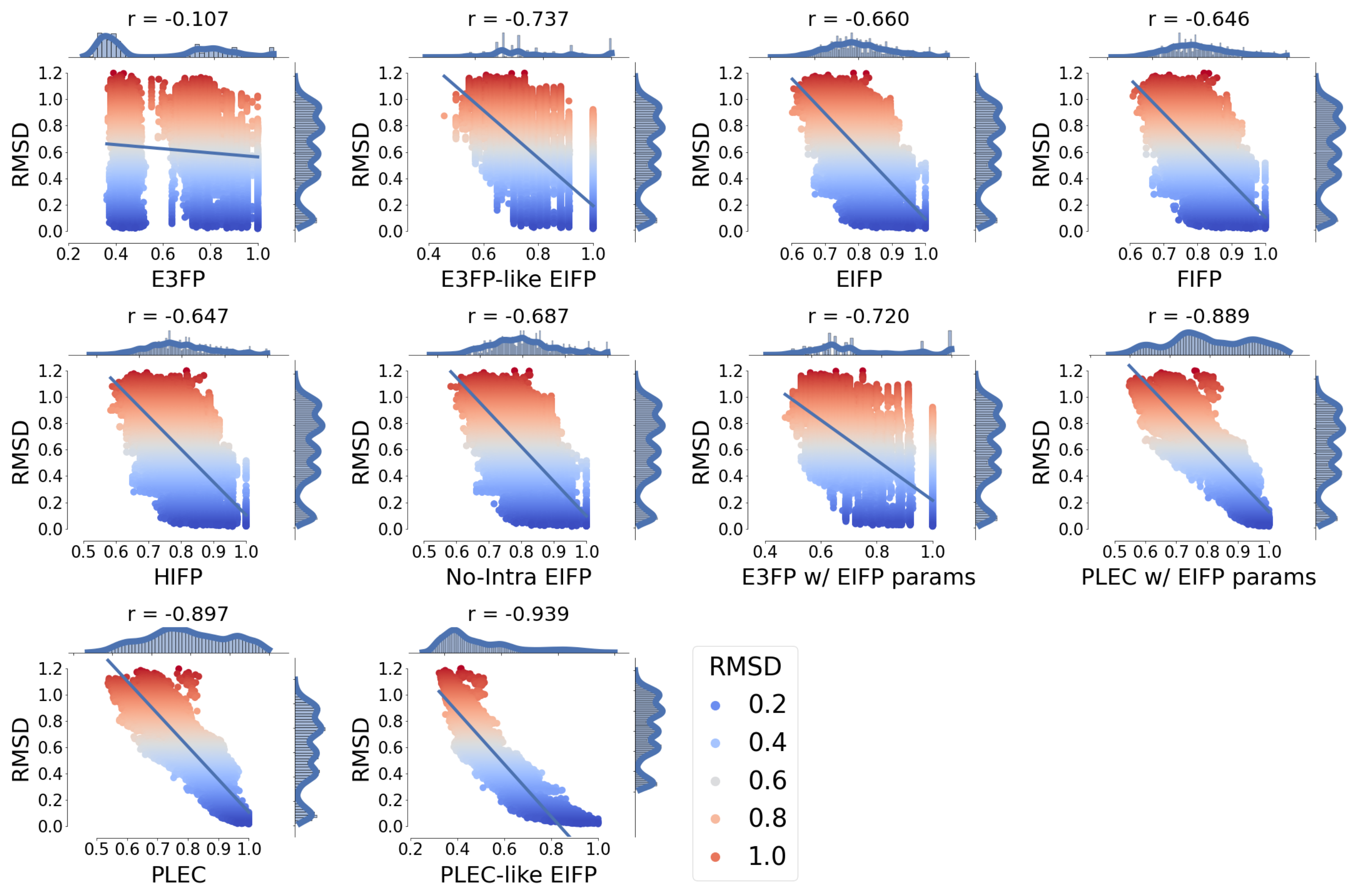


Figure S30: Correlation between the conformers' RMSD and the similarity of fingerprint pairs using the dataset "*Automatically generated conformers (0.8 Å)*".

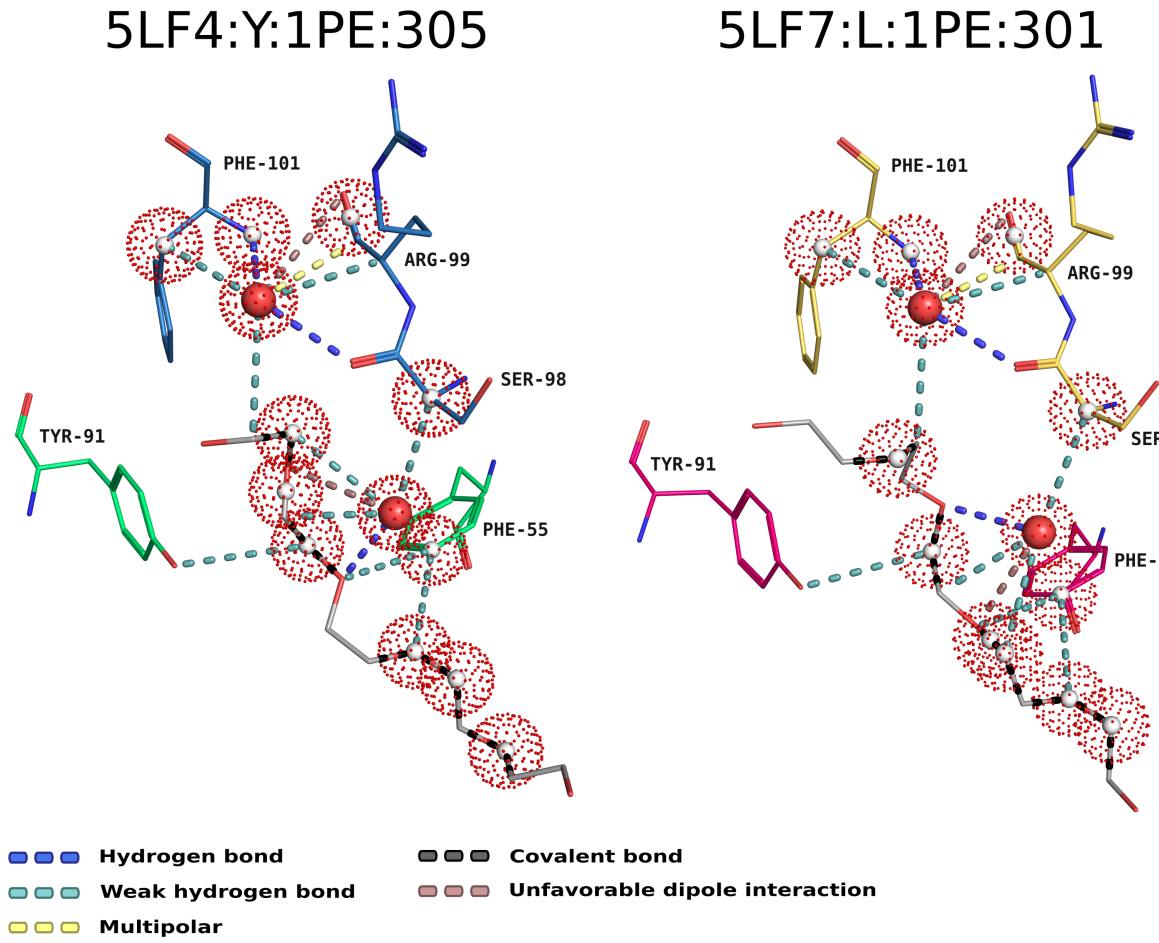


Figure S31: Common EIFP features between two complexes containing different proteins and the same ligand.

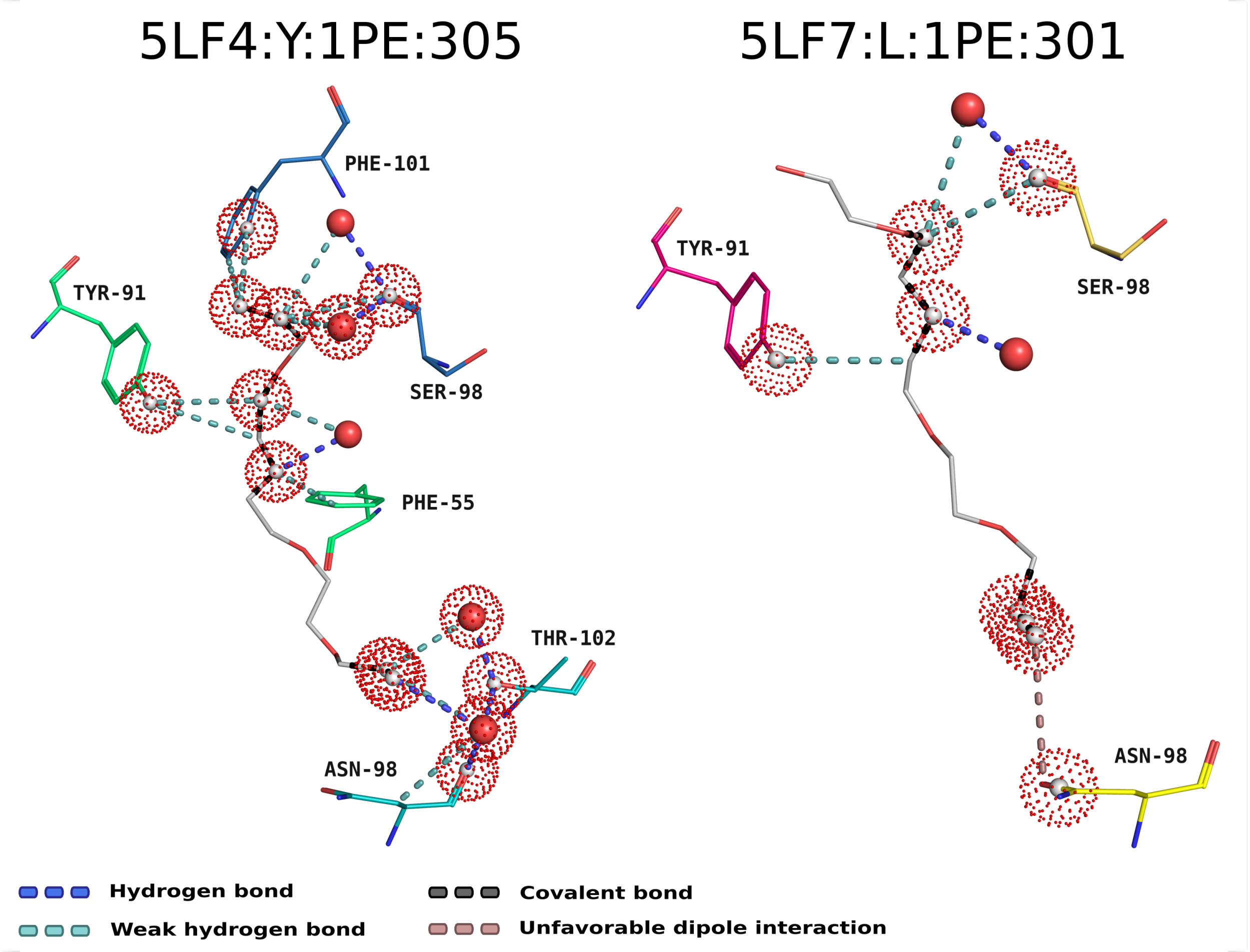


Figure S32: Exclusive EIFP features between two complexes containing different proteins and the same ligand.

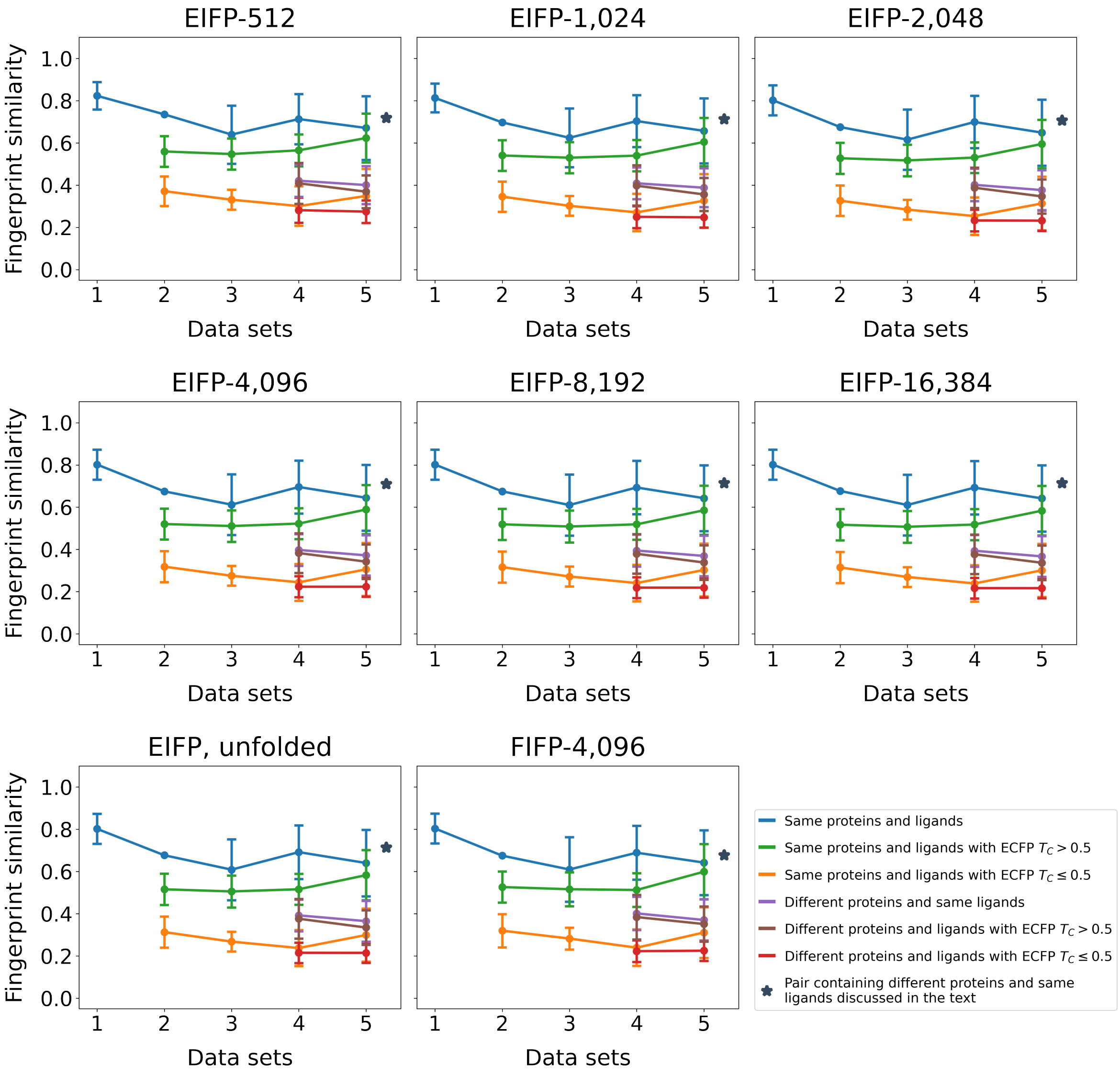


Figure S33: Similarity between EIFP pairs considering different FP lengths for the five data sets presented in Table 3. Pairs of complexes were divided into six groups based on their proteins and the molecular similarity (ECFP) between their ligands. Marker represents complexes with same ligands discussed in the text.

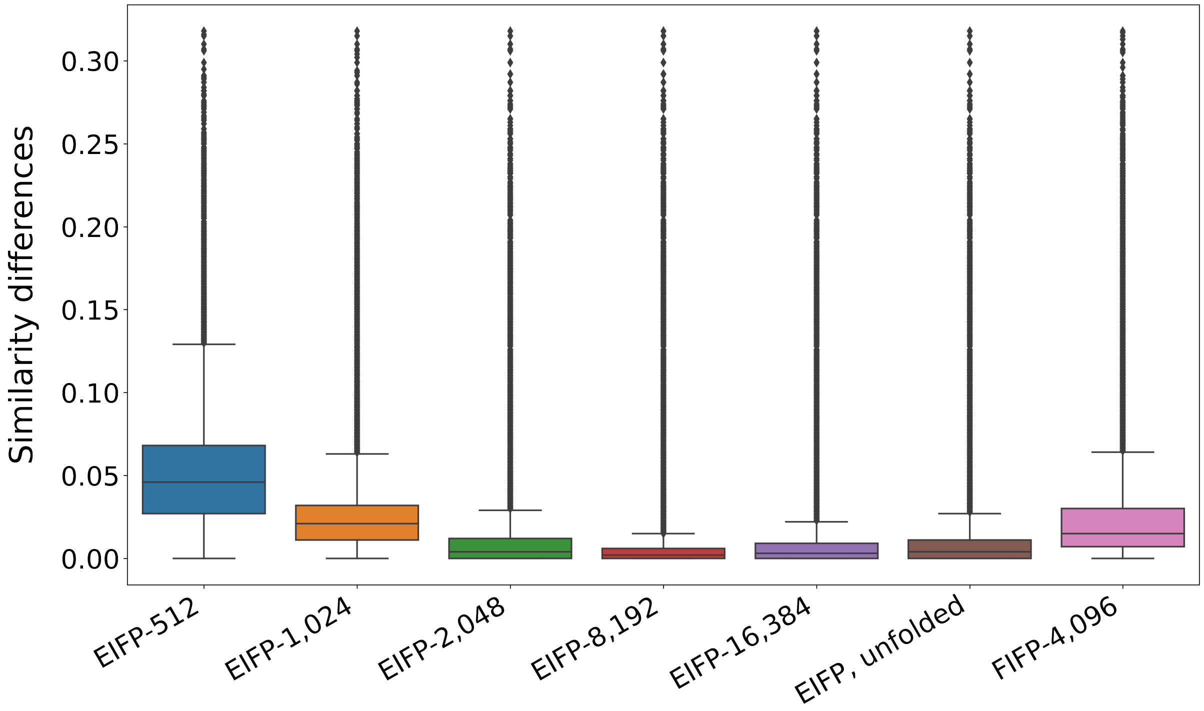


Figure S34: Similarity differences between *EIFP-4,096* and other IFPs using all PLI complexes' pairs.

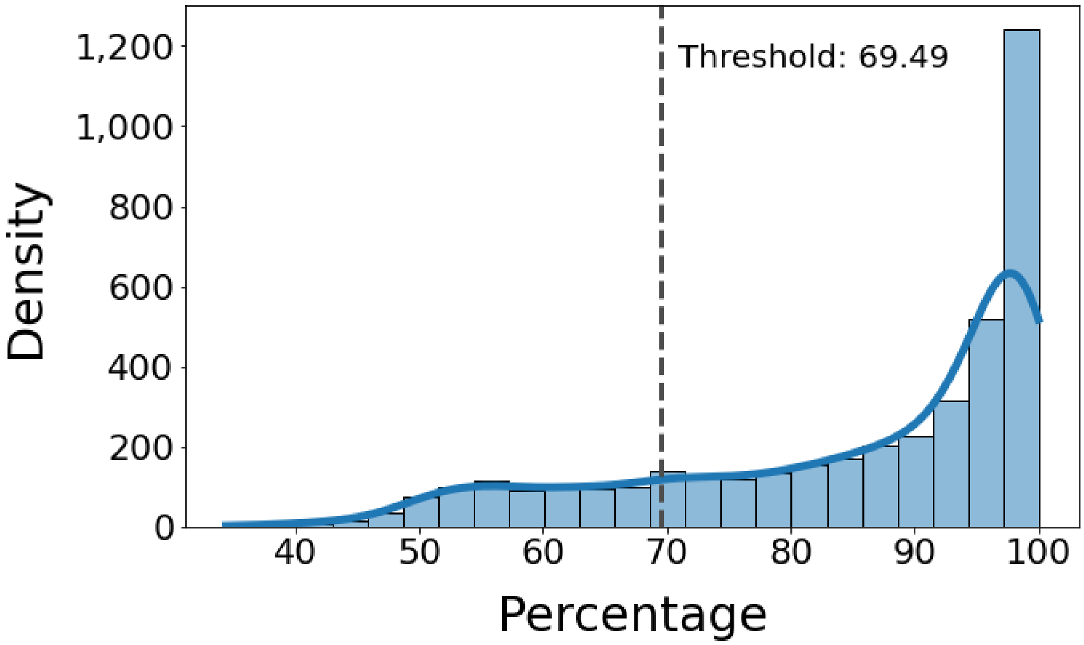


Figure S35: Percentage frequencies for the prevalent classes at each one of the 4,096 features. The most common class in a given feature was only assigned to the feature whether its percentage was greater than or equal to the threshold (69.49%, chosen by z-scores; dashed line), which was defined as the minimum percentage among the features presenting z-scores higher than 1. The blue line depicts the kernel density estimate for the percentages.

(a)

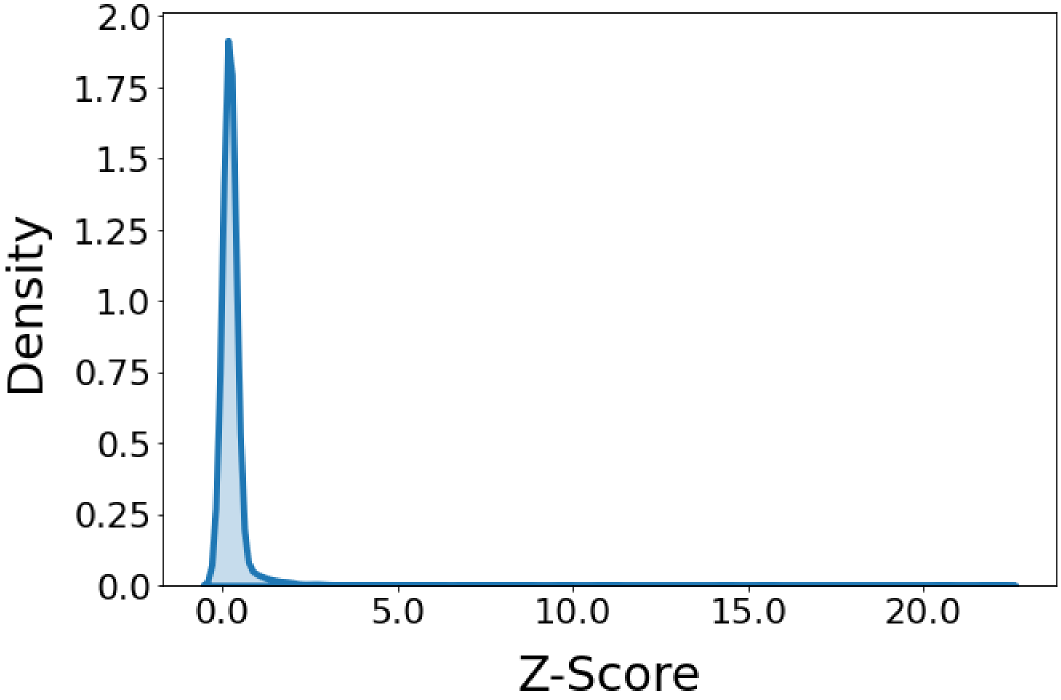


(b)

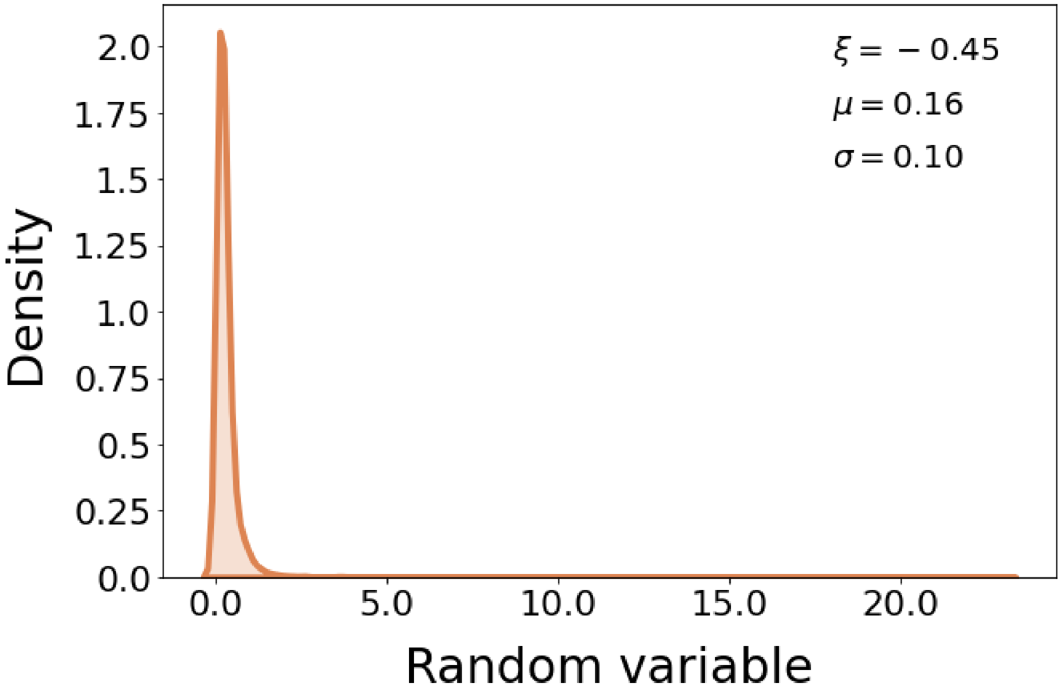


Figure S36: (a) Kernel density estimates for z-scores calculated from the feature importance scores and (b) random variables obtained from a generalized extreme value distribution fitted to the z-scores.

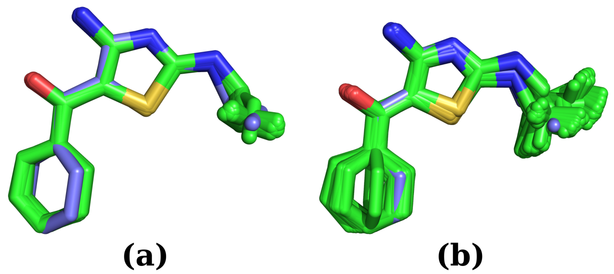

Figure S37: Automatically generated conformers whose distance to the ligand X02 (PDB id 3QQK) is 0.4 Å(a) and 0.8 Å(b). The original pose and the conformers are shown as blue and green sticks, respectively.

Table S7: List of ligands considered crystallography artifacts.[1,2]

| Ligand ids |
|---|
| ACE, ACT, BME, CSD, CSW, EDO, FMT, GOL, MSE, NAG, NO3, PO4, SGM, SO4, TPO |

Table S8: List of ions.[3]

| Ligand ids |
|---|
| SO3, CU, AUC, DTI, SB, TEA, HG, T1A, BR, 118, YB, IOD, CA, PT, NO3, GD3, PO3, MLI, GA, 3NI, OAA, CON, NGO, CHT, 2HP, MBH, CU1, PT4, NH4, OS, TRA, HAI, V, AZI, BF4, EU, RB, RU, 2MO, EMC, TBA, SEK, SMO, GEP, DSC, SOH, ACT, FPO, 3CO, BCT, MO6, SR, BA, AG, IN, NI, THE, OC1, RHD, Y1, NRU, CUA, 1AL, BO4, IR, MN3, LCO, CUZ, PO4, CS, WO4, SO4, CR, PTN, CE, EDR, AL, 4MO, OC2, PER, CD, FE2, IRI, LCP, DME, DMI, AU, LI, TMA, CO3, MLT, K, AU3, TL, ALF, CO, 3MT, VO4, CYN, CSB, CL, NO2, FE, PB, OC4, PR, TCN, PI, OH, SM, MOS, PBM, SE4, MG, F, PD, YT3, MMC, SCN, NET, IUM, EU3, WO5, MN, IR3, ZN, BEF, OS4, OXL, ATH, MH2, MN6, LA, CAC, LU, NCO, MAC, TB, YB2, W, MOO, NA, 6MO |

Table S9: List of cofactors.[4,5]

| Ligand ids |
|---|
| 01A, 01K, 0AF, 0ET, 0HG, 0HH, 0UM, 0WD, 0XU, 0Y0, 0Y1, 0Y2, 18W, 1C4, 1CV, 1CZ, 1DG, 1HA, 1JO, 1JP, 1R4, 1TP, 1TY, 1U0, 1VU, 1XE, 1YJ, 29P, 2CP, 2MD, 2NE, 2TP, 2TY, 36A, 37H, 3AA, 3CD, 3CP, 3GC, 3H9, 3HC, 48T, 4AB, 4CA, 4CO, 4IK, 4LS, 4LU, 4YP, 5AU, 5GY, 62X, 6FA, 6HE, 6NR, 6V0, 76H, 76J, 76K, 76L, 76M, 7AP, 7HE, 8EF, 8EL, 8EO, 8FL, 8ID, 8JD, 8PA, 8Z2, A3D, ABY, ACO, ACP, ADP, AGQ, AHE, AMP, AMX, ANP, AP0, ASC, AT5, ATA, ATP, B12, BCA, BCO, BHS, BIO, BOB, BSJ, BTI, BTN, BYC, BYG, BYT, C2F, CA3, CA5, CA6, CA8, CAA, CAJ, CAO, CCH, CDP, CIC, CMC, CMP, CMX, CNC, CND, CO6, CO8, COA, COB, COD, COF, COH, COM, COO, COT, COW, COY, COZ, CTP, D7K, DCA, DCC, DDH, DG1, DHE, DN4, DPM, DTB, EAD, EEM, EN0, ENA, EPY, ESG, F43, FA8, FAA, FAB, FAD, FAE, FAM, FAO, FAS, FCG, FCX, FDA, FDE, FED, FFO, FMI, FMN, FNR, FNS, FON, FOZ, FRE, FSH, FYN, G27, GBI, GBP, GBX, GDN, GDP, GDS, GF5, GGC, GIP, GMP, GNB, GNP, GPR, GPS, GRA, GS8, GSB, GSF, GSH, GSM, GSN, GSO, GTB, GTD, GTP, GTS, GTX, GTY, GVX, H2B, H4B, HAG, HAS, HAX, HBI, HCC, HDD, HDE, HEA, HEB, HEC, HEM, HIF, HMG, HSC, HTL, HXC, IBG, ICY, IRF, ISW, JM2, JM5, JM7, K15, L9X, LEE, LNC, LPA, LPB, LZ6, M43, M6T, MCA, MCD, MCN, MDE, MDO, MGD, MH0, MLC, MNH, MNR, MPL, MQ7, MSS, MTE, MTQ, MTV, MYA, N01, N1T, N3T, NA0, NAD, NAE, NAI, NAJ, NAP, NAQ, NAX, NBD, NBP, NDC, NDE, NDO, NDP, NHD, NHM, NHQ, NHW, NMX, NOP, NPL, NPW, ODP, OXK, P1H, P2Q, P3Q, P5F, PAD, PCD, PDP, PLP, PLR, PMP, PNS, PP9, PQQ, PXP, PZP, R1T, RBF, RFL, RGE, S0N, S1T, SA8, SAD, SAE, SAH, SAM, SCA, SCD, SCO, SDX, SFD, SFG, SH0, SHT, SMM, SND, SOP, SRM, SX0, T1G, T5X, T6F, TAD, TAP, TC6, TD6, TD7, TD8, TD9, TDK, TDL, TDM, TDP, TDT, TDW, TGG, THD, THF, THG, THH, THV, THW, THY, TOQ, TP7, TP8, TPP, TPQ, TPU, TPW, TPZ, TQQ, TRQ, TS5, TT8, TXD, TXE, TXP, TXZ, TYQ, TYY, TZD, UAH, UMP, UQ1, UQ2, UQ5, UQ6, VWW, WCA, WSD, WWF, XAX, XP8, XP9, Y7Y, YNC, ZBF, ZEM, ZID, ZNH, ZOZ |

(5) Mukhopadhyay, A.; Borkakoti, N.; Pravda, L.; Tyzack, J. D.; Thornton, J. M.; Velankar, S. Finding enzyme cofactors in Protein Data Bank. *Bioinformatics* **2019**, *35*, 3510–3511.
