## Supporting information - additional methods for "Prioritizing virtual screening with interpretable interaction fingerprints"

#### Physicochemical feature assignment

The feature assignment is performed on the fly for all molecules within a certain distance (in Å) of the defined target, which can be a chain, a compound, or a list of compounds.

Thus, the target and the recovered molecules around it define the binding site scope for the interaction analysis. For each one of these compounds, LUNA applies three main procedures.

First, it verifies whether the current compound contains a defined molecule file, which is usually the case for docking campaigns since the ligand pose may be available as a separate molecular file. In this particular case, the third step is promptly initialized.

Secondly, the tool will identify all molecules covalently bound to the current compound and convert their structures from PDB to Mol format using Open Babel.[1] This conversion is essential because a PDB file does not contain crucial chemical information for a proper physicochemical property perception. Moreover, it is wise to convert the current compound with its bound neighbors in order to keep correct bond orders. If necessary, hydrogens can also be added to the molecules during the conversion according to the specified pH.

Another optional step consists of validating the converted molecules, standardize protein residues, and amending simple problems related to valence and charge that may occur when reading molecules from PDB files. That happens precisely because the PDB format does not have sufficient chemical information, leading Open Babel or RDKit to perceive an atom's bond order, aromaticity, valence, or charge incorrectly. Our implemented solutions cover simple problems that are more recurrent during these conversions. For example, atom charges are amended whether they do not match the expected charge according to our implementation of OpenEye's charge model[1]. Additionally, valences are amended only for ammonium nitrogen whose structure was not previously ionized. In this case, Open Babel may perceive such atoms as hypervalent and attribute an incorrect valence to the nitrogen.

Finally, LUNA perceives physicochemical properties through a set of chemical rules specified as a SMARTS-based language string and stored in a feature definition file format (FDef).[2] Our rules comprise both atoms and groups of atoms, which can represent a functional group or an arrangement of atoms. Importantly, tautomeric forms of chemical groups were also envisioned in our rules as a means to account for the biological environment dynam-

[1] https://docs.eyesopen.com/toolkits/python/oechemtk/valence.html

ics. Another optional step can be applied after the third procedure and consists of grouping hydrophobic atoms to form a hydrophobe group. See Section *Hydrophobic interaction* for more information.

The complete set of rules are presented in the next section.

#### Physicochemical property definitions

***Hydrogen donor*** labels are assigned to atoms that fulfill the following rules:

1. It is a tertiary amine N (‘`$([Nv3](-C)(-C)-C)`’) that is not:
    (a) an amide-like N (‘`$([#7][C,P,S]=O)`’);
    (b) an amidine-like N (‘`[N;![$([#7][C,P,S]=O)];$(N=[CX3][N;![$([#7][C,P,S]=O)]]),$(N[CX3]=[N;![$([#7][C,P,S]=O)]])]`’).
2. It is a tautomeric aromatic N (‘`[$(n[n;H1]),$(nc[n;H1])]`’) not in a tetrazole (‘`[nR1r5;$(n:n:n:n:c),$(n:n:n:c:n)]`’);
3. It is a tautomeric N in a guanidine-like group (‘`[$([NX2H0]=[CH0X3](N)N)]`’);
4. It is a N/O/S atom bound to a hydrogen atom (‘`[N!H0v3,N!H0+v4,nH+0,OH2v2,OH+0,SH+0]`’), where the N is not acidic (‘`[$([NH,NH0-1](S(=O)(=O))(C(=O))),$([NH1,NH0-1;R](C(=O))(C(=[O,S])))]`’) and does not belong to a tetrazole (‘`[nR1r5;$(n:n:n:n:c),$(n:n:n:c:n)]`’), while O and S are also not acidic (‘`[$([O][C,S,P](=[O,S]))]`’);
5. It is a tautomeric O in a ketene acetal-like group (‘`[$([O;H1,H0&-1]-[#6;X3]-,:[#8])]`’).

***Halogen donor*** labels are assigned to atoms that fulfill the following rules:

1. It is a Cl/Br/I bound to C/S (‘`[$([Cl,Br,I;X1]-[#6])]`’).

***Chalcogen donor*** labels are assigned to atoms that fulfill the following rules:

1. It is a divalent S/Se/Te bound to C/S (‘`[$([#16,#34,#52;v2;H0](-,:[#6,#16])-,:[#6,#16]),$([#16,#34,#52;v2;H1][#6,#16])]`’) or in an isothiazole-like group (‘`[$([#16,#34,#52;v2;H0;a](n)c)]`’).

***Hydrogen/Halogen/Chalcogen acceptor*** labels are assigned to atoms that fulfill the following rules:

1. It is a tautomeric aromatic N (‘`[$([n;H1]n),$([n;H1]cn)]`’) not in a tetrazole (‘`[nR1r5;$(n:n:n:n:c),$(n:n:n:c:n)]`’);
2. It is an aromatic N with a double bond and no hydrogen (‘`[n;+0;H0;!X3]`’);
3. It is a N that is not:
   (a) a positive N (‘`[*;+1,+2,+3]`’);
   (b) an aniline-like N (‘`[$(N[a])]`’);
   (c) an amide-like N (‘`[$([#7][C,P,S]=O)]`’);
   (d) an amidine-like N (‘`[N;![$([#7][C,P,S]=O)];$(N=[CX3][N;![$([#7][C,P,S]=O)]]),$(N[CX3]=[N;![$([#7][C,P,S]=O)]])]`’);
   (e) a nitro-like N (‘`[$([N+]-[O-])]`’);
   (f) a basic N (‘`[$([N;H2&+0][CX4]),$([N;H1&+0]([CX4])[CX4]),[$([N;H0&+0]([CX4])([CX4])[CX4])]`’).
4. It is a chalcogen (‘`[O,$([S;!v4;!v6])]`’).

***Weak hydrogen donor*** labels are assigned to atoms that fulfill the following rules:

1. It is a C bound to at least one hydrogen (’`[#6;!H0]`’).

***Weak hydrogen acceptor*** labels are assigned to atoms that fulfill the following rules:

1. It is a neutral aromatic O or S (‘`[o,s;+0]`’);
2. It is a F bound to C (‘`[F;$(F-[#6]);!$(FC[F,Cl,Br,I])]`’).

***Positively ionizable*** labels are assigned to atom and atom groups that fulfill the following rules:

1. It is a basic N (‘`[$([N;H2&+0][CX4]),$([N;H1&+0]([CX4])[CX4]),[$([N;H0&+0]([CX4])([CX4])[CX4])]`’);
2. It is a guanidine-like group (‘`N[CH0X3](=N)N`’);
3. It is an amidine-like group (‘`[N;![$([#7][C,P,S]=O)]]=[CX3][N;![$([#7][C,P,S]=O)]]`’);
4. It is a 4-aminopyridine (‘`Nc1cc[nH0]cc1`’) or 2-aminopyridine (‘`Nc1cccc[nH0]1`’);
5. It is an imidazole-like group (‘`[n;R1]1[c;R1][n;R1][c;R1][c;R1]1`’);
6. It is positive atom not bound to a negative atom (‘`[[*;+1,+2,+3];!$(*~[*;-1,-2,-3])]`’).

***Negatively ionizable*** labels are assigned to atom and atom groups that fulfill the following rules:

1. It is a carboxylic-like group (‘`C(=[O,S])-[O;H1,H0&-1]`’);
2. It is a ketene acetal-like group (‘`[O;H1,H0&-1]-[#6;X3]-,:[#8]`’);
3. It is a tetrazole (‘`c1nnn[nH,n-1]1`’);
4. It is a barbiturate-like group (‘`O=C1CC(=O)[NH1,NH0-1;R]C(=O)[NH1,NH0-1;R]1`’);
5. It is a thiazolidinedione-like group (‘`O=C1[NH1,NH0-1;R]C(=[O,S])[SX2H0R]C1`’);
6. It is a diformamide-like group (‘`[NH1,NH0-1;R](C(=O))(C(=O))`’);
7. It is one of the hydroxamic acid forms: O anion (‘`C(=O)[NX3]-[O;H1,H0&-1]`’), N anion (‘`C(=O)[N-1]-[OH1]`’) or its resonance form (‘`C(-[O-1])=N-[OH1]`’);
8. It is a sulfuric-like (‘`S(=[O,S])(=O)(-O)-[O;H1,H0&-1]`’) or sulfonic-like (‘`S(=[O,S])(=O)-[O;H1,H0&-1]`’) or sulfinic-like (‘`S(=[O,S])-[O;H1,H0&-1]`’) acid group;

9. It is a acyl sulfonamide-like (‘[NH,NH0-1](S(=O)(=O))(C(=O))’) or a sulfonamide-like (‘[N;!H0,H0&-1]S(=O)(=O)’) group;
10. It is a phosphoric-like (‘P(=[O,S])(-O)(-O)-[O;H1,H0&-1]’) or phosphonic-like (‘P(=[O,S])(-O)-[O;H1,H0&-1]’) or phosphinic-like (‘P(=[O,S])-[O;H1,H0&-1]’) acid groups;
11. It is a negative atom not bound to a positive atom (‘[[*;-1,-2,-3]; !$(*∼[*;+1,+2,+3])]’).

***Nucleophile*** labels are assigned to atom groups that fulfill the following rules:

1. It is a halogen from a haloalkane (‘[F,Cl,Br,I;X1][#6]’);
2. It is an O in a carbonyl but not in a carboxylic-like group (‘[O;!$(O=C[O;H1, H0&-1])]=[C;!$(C([O;H1,H0&-1])=O)’);
3. It is an O in an alcohol but not in a carboxylic-like group (‘[O;v2;H1;!$(OC=[O,S])][#6;!$(C(=[O,S])[O;v2;H1])]’);
4. It is a cyano-like N (‘N#C’);
5. It is an O in a water molecule (‘[O;v2;H2]’);
6. It is a sulfonyl-like O (‘[$(O=[S;v4,v6]([#6])[#6])]=[$([S;v4,v6]([#6])([#6])=O)]’);
7. It is ketene acetal-like O (‘[$([O;H1,H0&-1]-[#6;X3]-,:[#8])][$([#6;X3](-,:[#8])[O;H1,H0&-1])]’);
8. It is a nitro-like O (‘[$(O=[N;D3;+][O-]),$([O-][N;D3;+]=O)]∼[$([N;D3;+](=O)[O-])]’);
9. It is an ether-like O (‘[$([#8;v2]([#6])[#6])][$([#6][#8;v2][#6])]’);
10. It is a thioether-like S (‘[$([#16;v2]([#6])[#6])][$([#6][#16;v2][#6])]’);
11. It is a hydroxylamine-like O (‘[$(O-[#7;H0;X3]([#7,#6])[#7,#6])][$([#7;H0;X3]([#7,#6])([#7,#6])O)]’).

***Electrophile*** labels are assigned to atom groups that fulfill the following rules:

1. It is a haloalkane C (‘`[#6][F,Cl,Br,I;X1]`’);
2. It is a C in a carbonyl but not in a carboxylic-like group (‘`[#6;!$(C([O;H1,HO&-1])=[O,S])]=[O;!$([O;H1,HO&-1]C=[O,S])]`’);
3. It is a cyano-like C (‘`C#N`’);
4. It is a nitro-like N (‘`[$([N;D3;+](=O)[O-])]~[$(O=[N;D3;+][O-]),$([O-][N;D3;+]=O)]`’).

***Aromatic*** labels are assigned to atom groups that fulfill the following rules:

1. 4-membered rings (‘`[a;r4,!R1&r3]1:[a;r4,!R1&r3]:[a;r4,!R1&r3]:[a;r4,!R1&r3]:1`’);
2. 5-membered rings (‘`[a;r5,!R1&r4,!R1&r3]1:[a;r5,!R1&r4,!R1&r3]:[a;r5,!R1&r4,!R1&r3]:[a;r5,!R1&r4,!R1&r3]:[a;r5,!R1&r4,!R1&r3]:1`’);
3. 6-membered rings (‘`[a;r6,!R1&r5,!R1&r4,!R1&r3]1:[a;r6,!R1&r5,!R1&r4,!R1&r3]:[a;r6,!R1&r5,!R1&r4,!R1&r3]:[a;r6,!R1&r5,!R1&r4,!R1&r3]:[a;r6,!R1&r5,!R1&r4,!R1&r3]:[a;r6,!R1&r5,!R1&r4,!R1&r3]:1`’);
4. 7-membered rings (‘`[a;r7,!R1&r6,!R1&r5,!R1&r4,!R1&r3]1:[a;r7,!R1&r6,!R1&r5,!R1&r4,!R1&r3]:[a;r7,!R1&r6,!R1&r5,!R1&r4,!R1&r3]:[a;r7,!R1&r6,!R1&r5,!R1&r4,!R1&r3]:[a;r7,!R1&r6,!R1&r5,!R1&r4,!R1&r3]:[a;r7,!R1&r6,!R1&r5,!R1&r4,!R1&r3]:[a;r7,!R1&r6,!R1&r5,!R1&r4,!R1&r3]:1`’);
5. 8-membered rings (‘`[a;r8,!R1&r7,!R1&r6,!R1&r5,!R1&r4,!R1&r3]1:[a;r8,!R1&r7,!R1&r6,!R1&r5,!R1&r4,!R1&r3]:[a;r8,!R1&r7,!R1&r6,!R1&r5,!R1&r4,!R1&r3]:[a;r8,!R1&r7,!R1&r6,!R1&r5,!R1&r4,!R1&r3]:[a;r8,!R1&r7,!R1&r6,!R1&r5,!R1&r4,!R1&r3]:[a;r8,!R1&r7,!R1&r6,!R1&r5,!R1&r4,!R1&r3]:[a;r8,!R1&r7,!R1&r6,!R1&r5,!R1&r4,!R1&r3]:[a;r8,!R1&r7,!R1&r6,!R1&r5,!R1&r4,!R1&r3]:1`’);

6. 9-membered rings (‘`[a;r9,!R1&r8,!R1&r7,!R1&r6,!R1&r5,!R1&r4,!R1&r3]1:[a;r9,!R1&r8,!R1&r7,!R1&r6,!R1&r5,!R1&r4,!R1&r3]:[a;r9,!R1&r8,!R1&r7,!R1&r6,!R1&r5,!R1&r4,!R1&r3]:[a;r9,!R1&r8,!R1&r7,!R1&r6,!R1&r5,!R1&r4,!R1&r3]:[a;r9,!R1&r8,!R1&r7,!R1&r6,!R1&r5,!R1&r4,!R1&r3]:[a;r9,!R1&r8,!R1&r7,!R1&r6,!R1&r5,!R1&r4,!R1&r3]:[a;r9,!R1&r8,!R1&r7,!R1&r6,!R1&r5,!R1&r4,!R1&r3]:[a;r9,!R1&r8,!R1&r7,!R1&r6,!R1&r5,!R1&r4,!R1&r3]:[a;r9,!R1&r8,!R1&r7,!R1&r6,!R1&r5,!R1&r4,!R1&r3]:1`’).

***Hydrophobic*** labels are assigned to atoms that fulfill the following rules:

1. It is a divalent sulfur, halogen (except fluorine), or a carbon not bound to an electronegative atom (‘`[s,S&H0&v2,Br,I,Cl,At,[#6;+0;![#6;$([#6]~[#7,#8,#9])]];+0]`’).

***Amide*** labels are assigned to atom groups that fulfill the following rules:

1. It is an amide-like group (‘`[NX3][CX3](=[OX1])`’).

***Atom*** labels are assigned to atoms that fulfill the following rules:

1. It is any heavy atom (‘`[!#1])`’).

### Geometrical criteria for computing molecular interactions

In this section, we present the geometrical criteria and the models employed to calculate molecular interactions in LUNA. Figures S1, S2, and S3 show the geometrical models for most of the interactions. Not all interactions are shown in the diagrams because they require only the evaluation of Euclidean distances between two atoms (atom groups). For these cases, the methods are discussed directly in their respective section.

It is important to mention that all geometrical criteria and models presented in this section consist of the default model of LUNA, but all of them are customizable.

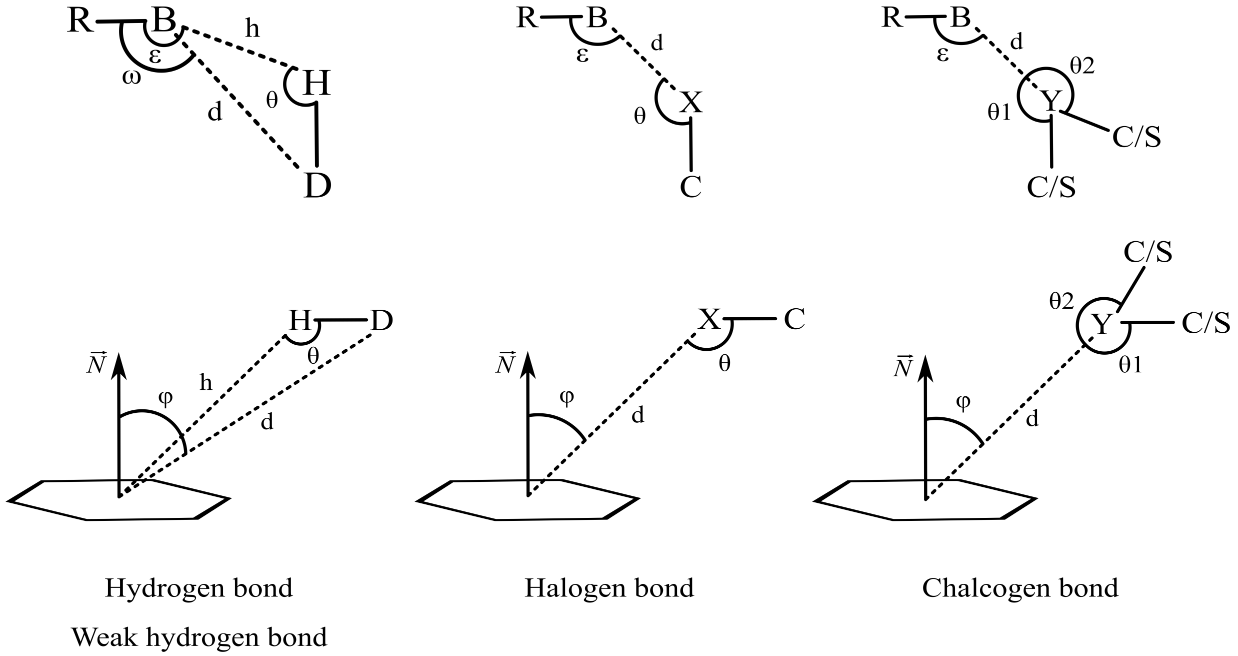


Figure S1: Models for calculating hydrogen, weak hydrogen, halogen, and chalcogen bonds. The definitions for each letter and angle depicted in the diagrams are explained in their respective interaction section.

##### Hydrogen bonds

Hydrogen bonds are identified according to[3] and its calculation depends on five parameters as shown in Figure S1, where $D$, $H$, $B$, and $R$ are the donor, hydrogen, a Lewis base (acceptor), and an atom covalently bound to the acceptor, respectively. Below we provide the default values extracted from literature[3–6] for each of the parameters presented in Figure S1.

- Default distance thresholds: $d \leq 3.9$Å and $h \leq 2.8$Å, respectively;
- Default angle thresholds: $\theta, \varepsilon, \omega \geq 90°$.

In Figure S1, the model depicts only one hydrogen to the donor and one neighbor for the acceptor. However, if the donor contains two or more hydrogens, each one of them

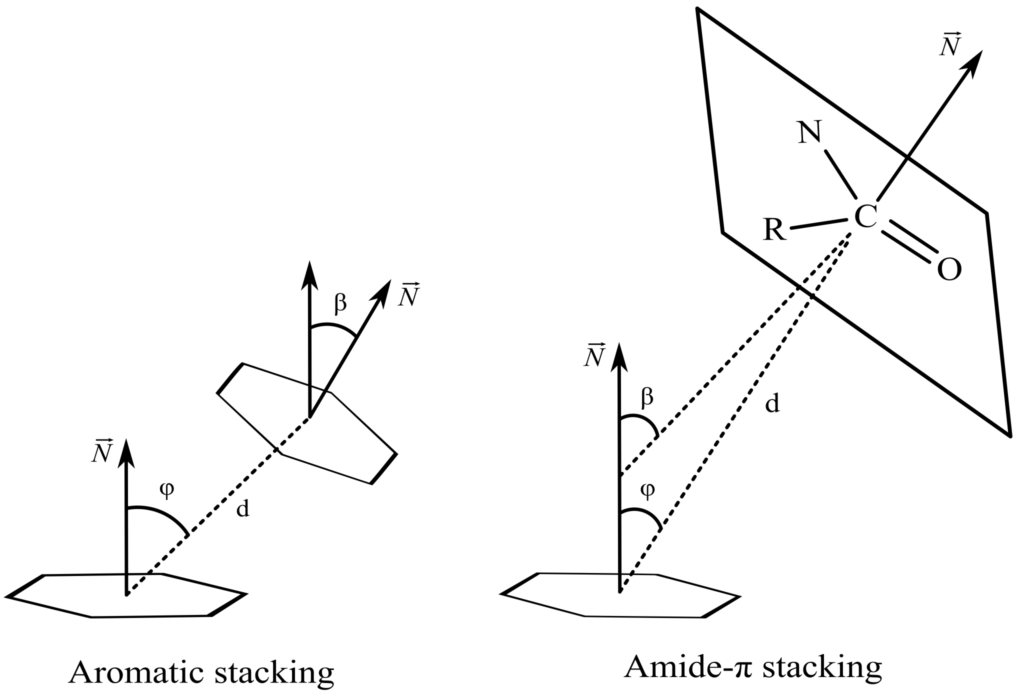


Figure S2: Models for calculating aromatic stackings and amide-$\pi$ stackings. The definitions for each letter and angle depicted in the diagrams are explained in their respective interaction section.

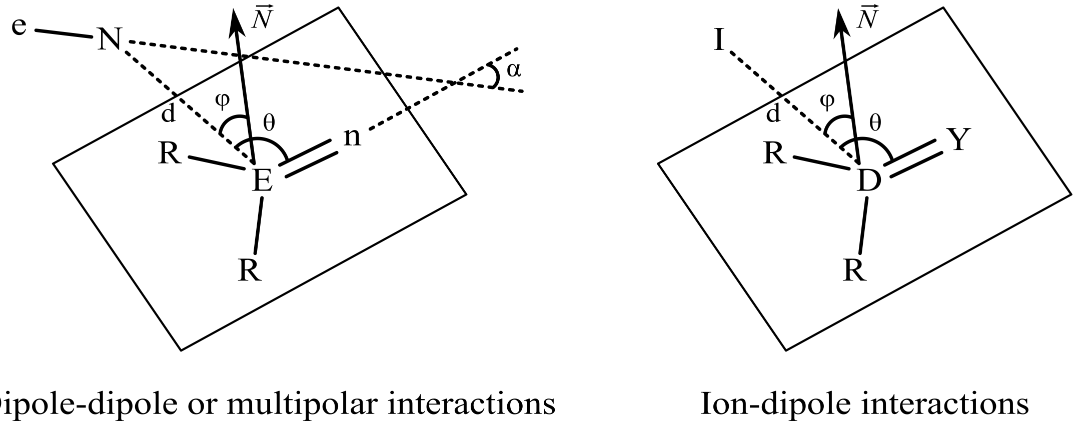


Figure S3: Models for calculating dipole-dipole (multipolar interactions) and ion-dipole interactions. The definitions for each letter and angle depicted in the diagrams are explained in their respective interaction section.

is evaluated against an acceptor, resulting in multiple hydrogen bonds. Similarly, when acceptors contain more than one neighbor, the hydrogen bond is only accepted if the angles involving these atoms match the angle criteria.

Lastly, the algorithm has two modes for applying the above criteria, a strict and loose one.

In the strict mode, donor atoms must have their hydrogens explicitly defined as the algorithm requires the hydrogens' coordinates. However, not all PDB structures contain hydrogens and sometimes a potential donor is not in its ionized or tautomeric form, which would impede the algorithm from detecting hydrogen bonds. Also, consider the dynamic of water molecules. In Open Babel, for instance, when hydrogens are included in the structure, water molecules usually can have their hydrogens added in different manners. As a consequence, if the exact position of hydrogens is taken into account, several results would be possible for the same system.

Bearing that in mind, we also provide a loose mode where the system dynamics are considered. When the loose mode is activated, hydrogen bonds are identified despite the presence of hydrogens. Furthermore, with this mode, all hydrogen bonds involving solvents are loosely validated even if they contain hydrogens explicitly defined, which is a measure acknowledging their dynamic and multiple possible hydrogen placement.

The loose mode works similarly as in McDonald and Thornton:[3] angles involving hydrogens are ignored, and the hydrogen-acceptor distance is calculated as if the hydrogen was placed 1 Å away from the donor in the direction of the acceptor. As a mathematical expression, it can be defined as follows:

$$h = d - 1 \leq 2.8 \text{ Å} \tag{1}$$

It is noteworthy that this method may generate more interactions than the strict one, and some of the identified interactions may be false positives.

##### Weak hydrogen bonds

Weak hydrogen bonds are identified through two models depending on the acceptor atom. For single atoms (Lewis bases), the model is the same as presented for hydrogen bonds. However, herein, $D$ is a weak hydrogen donor (a carbon with an attached hydrogen bond), and $B$ is an acceptor or a weak acceptor (any aromatic oxygen/sulfur or fluorine).[7–10] In its turn, the second model comprises aromatic rings as acceptors. Below we provide the default values extracted from literature[8,11,12] for each of the parameters presented in Figure S1.

- Conventional weak hydrogen bonds:
  - Default distance thresholds: $d \leq 4$Å and $h \leq 3$Å, respectively;
  - Default angle thresholds: $\theta \geq 110°$ and $\varepsilon, \omega \geq 90°$;
- Weak hydrogen bonds involving aromatic rings:
  - Default distance thresholds: $d \leq 4.5$Å and $h \leq 3.5$Å, respectively. The distances are calculated in relation to the ring centroid;
  - Default angle thresholds: $\theta \geq 120°$ and $\varphi \leq 40°$, where $\varphi$ (displacement angle) is the angle formed by the ring normal ($\overrightarrow{N}$) and the vector between the ring centroid and the donor atom.

Similarly to hydrogen bonds, multiple weak hydrogen bonds can also be identified whether more than one hydrogen is covalently bound to the weak hydrogen donor. It also evaluates all possible angles with the acceptor's neighbor.

Additionally, weak hydrogen bonds can also work on either strict or loose modes as presented for hydrogen bonds. Thus, the distance expression in the loose mode is defined as follows:

$$h = d - 1 \leq 3\text{Å} \tag{2}$$

##### Water-bridged hydrogen bond

Water-bridged hydrogen bonds are identified by directly searching for pairs of compound-water hydrogen bonds, where both bonds involve the same water molecule. Departing from a valid pair of hydrogen bonds, a new interaction connecting the involved compounds is created and labeled *water-bridged hydrogen bond.*

These special hydrogen bonds are not identified by default because the interactions required for its computation already account for its existence implicitly. However, if necessary, this interaction can be easily turned on through a flag in the method for calculating interactions.

##### Halogen bond

Halogen bonds are identified according to the model presented in Figure S1, where $X$, $C$, $B$, $R$ are a halogen, a carbon bound to the halogen, a Lewis base (acceptor), and an atom covalently bound to the acceptor, respectively. Below we provide the default values extracted from literature[13,14] for each of the parameters presented in Figure S1.

- Conventional halogen bonds:
  - Default distance threshold: $d \leq 4$Å;
  - Default angle thresholds: $\theta \geq 120°$ and $\varepsilon \geq 80°$;
- Halogen bond involving aromatic rings:
  - Default distance threshold: $d \leq 4.5$Å, where $d$ is calculated in relation to the ring centroid;
  - Default angle thresholds: $\theta \geq 120°$ and $\varphi \leq 60°$, where $\varphi$ (displacement angle) is the angle formed by the ring normal ($\overrightarrow{N}$) and the vector between the ring centroid and the halogen.

As pointed out by Cavallo et al.,[15] multiple carbons may be bound to the halogen, which, in its turn, may result in multiple halogen bonds with the same acceptor. Moreover, when the acceptor contains more than one neighbor, the tool will evaluate all possible angles formed with these atoms, and the interaction is only accepted if all angles match the criteria.

##### Chalcogen bond

Chalcogen bonds are identified according to the model presented in Figure S1, where $Y$, $C/S$, $B$, $R$ are a chalcogen, a carbon/sulfur bound to the chalcogen, a Lewis base (acceptor), and an atom covalently bound to the acceptor, respectively. Below we provide the default values extracted from literature[16–18] for each of the parameters presented in Figure S1.

- Conventional chalcogen bonds:
  - Default distance threshold: $d \leq 4$Å;
  - Default angle thresholds: $\theta \geq 120°$ and $\varepsilon \geq 80°$;
- Chalcogen bond involving aromatic rings:
  - Default distance threshold: $d \leq 4.5$Å, where $d$ is calculated in relation to the ring centroid;
  - Default angle thresholds: $\theta \geq 120°$ and $\varphi \leq 60°$, where $\varphi$ (displacement angle) is the angle formed by the ring normal ($\overrightarrow{N}$) and the vector between the ring centroid and the chalcogen.

We mentioned that the donor atoms might contain more than one neighbor covalently bound to it in hydrogen and halogen bonds. However, note that this statement is only explicitly depicted for chalcogen bonds. That is because they are mainly established by divalent chalcogens. Consequently, chalcogen bonds could also establish multiple chalcogen bonds to the acceptor.

Finally, when multiple neighbors are bound to the acceptor atom, all angles formed with them should match the criteria; otherwise, the interaction is not accepted.

##### Aromatic stacking

Aromatic stackings are identified as presented in Figure S2 and are classified according to their geometrical arrangements into nine different stackings (Figure S4) as in Bhattacharyya et al..[19] Below we provide the default values extracted from literature[19,20] for each of the parameters presented in Figure S2.

- Default distance threshold: $d \leq 6$Å, where $d$ is calculated in relation to the ring centroids;
- Default angle thresholds: each specific stacking depends on a combination of $\varphi$ and $\beta$ angles, as shown in Figure S4, where $\varphi$ (displacement angle) is the angle formed by the ring normal ($\overrightarrow{N}$) and the vector between the two ring centroids; while $\beta$ (dihedral angle) is the angle between the two ring planes, which is calculated by the angle formed by the ring normals. The displacement angle is calculated using both rings as references, and the smallest angle is chosen for defining the stacking type.

If the user decides not to use the angle criteria, the interaction will be labeled as a general *aromatic stacking*.

**Amide-$\pi$ stacking**

Amide-$\pi$ stackings are identified according to the model presented in Figure S2. Below we provide the default values extracted from literature[12,21–23] for each of the parameters presented in Figure S2.

- Default distance threshold: $d \leq 4.5$Å, where $d$ is calculated in relation to the ring and amide centroids;
- Default angle thresholds: $\varphi, \beta \leq 30°$, where $\varphi$ (displacement angle) is the angle formed by the ring normal ($\overrightarrow{N}$) and the vector between the ring and amide centroids; while $\beta$ (dihedral angle) is the angle between the ring and amide planes, which is calculated by the angle formed by their normals.

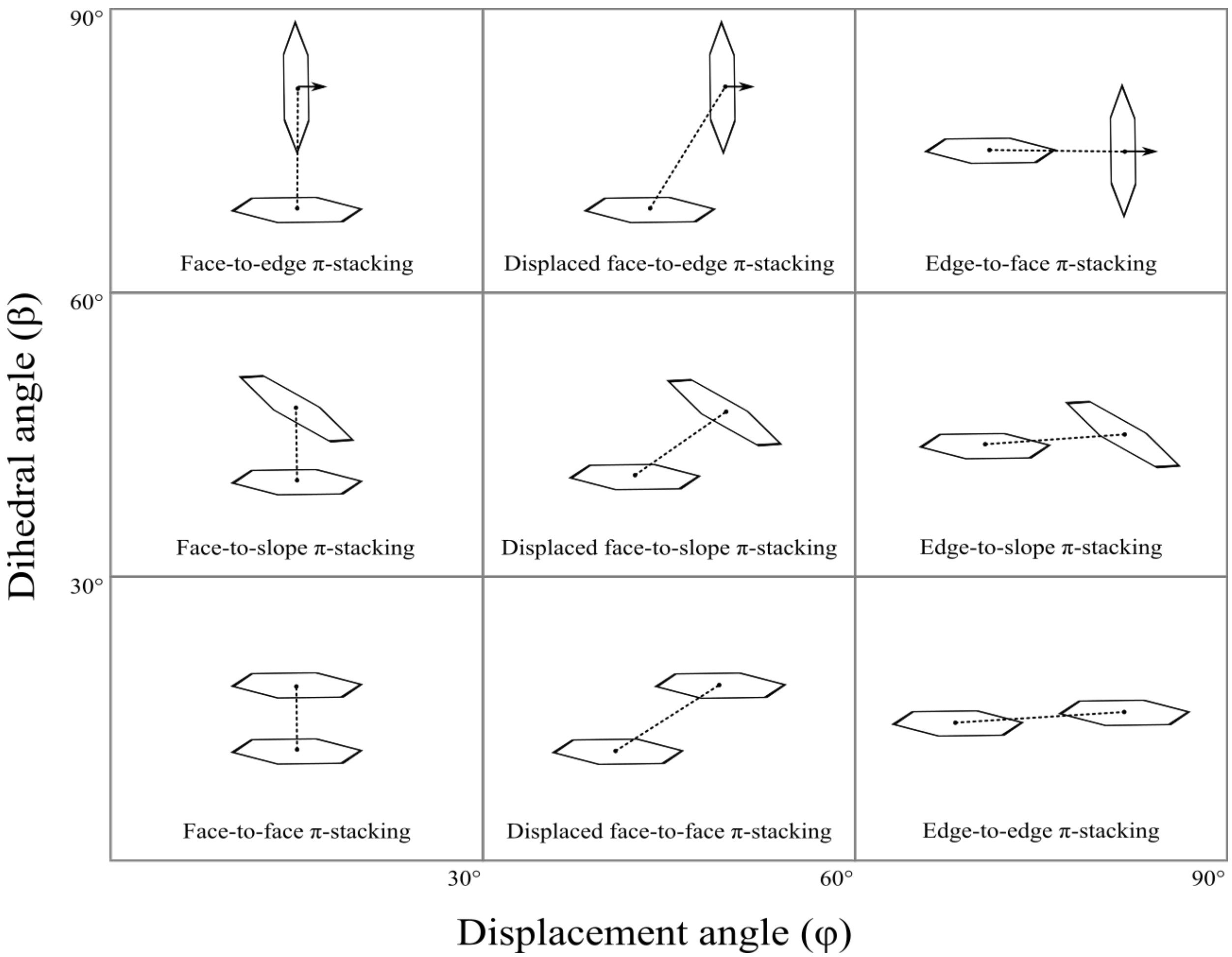


Figure S4: Classification of aromatic stackings according to the angles $\varphi$ (displacement angle) and $\beta$ (dihedral angle) given two aromatic rings.

**Dipole-dipole or multipolar interactions**

Multipolar interactions are identified according to the model presented in Figure S3, where E═n and N—e are the dipoles containing the interacting electrophile ($E$) and nucleophile ($N$); and R are the atoms covalently bound to the electrophile.

There are four possible arrangements for favorable multipolar interactions:[24] *parallel multipolar*, *antiparallel multipolar*, *orthogonal multipolar*, and *tilted multipolar*. Below we provide the default values extracted from literature[24] for each possible arrangement and the parameters presented in Figure S3.

- Default distance threshold: $d \leq 4$Å, where $d$ is calculated in relation to the nucleophilic and electrophilic atoms;

- Default angle thresholds: $70° \leq \theta \leq 110°$ and $\varphi \leq 40°$, where $\varphi$ (displacement angle) is the angle formed by the electrophile normal ($\overrightarrow{N}$) and the vector connecting the interacting electrophile and nucleophile. In its turn, $\alpha$ is the angle formed by the dipole vectors and determines the multipolar arrangements as follows:
  - Parallel multipolar: $\alpha \leq 25°$;
  - Antiparallel multipolar: $\alpha \geq 155°$;
  - Orthogonal multipolar: $70° \leq \alpha \leq 110°$;
  - Tilted multipolar: any $\alpha$ not comprised by the above criteria.

The algorithm also detects unfavorable dipole-dipole interactions, which are classified either as *unfavorable nucleophile-nucleophile* or *unfavorable electrophile-electrophile*. However, the arrangements presented above are not employed for unfavorable interactions. The tool only evaluates the distance $d$ and angles $\theta$ and $\varphi$ in the same way presented for the favorable interactions.

Also, it is noteworthy that although the diagram depicts the classic dipole-dipole interaction involving a carbonyl-like structure in the electrophile side of the interaction, other non-planar substructures are also accepted.

Lastly, in cases where the nucleophile neighbor ($e$) coordinate cannot be determined (e.g., hydrogens in water molecules), the angle $\alpha$ is not available. Therefore, the interaction is classified as a general *multipolar interaction* since it is not possible to define the proper dipole arrangement. In its turn, when the coordinate of the electrophile partner ($n$) is not available, it is only possible to calculate the distance between $E$ and $N$. Thus, we also opted for classifying the interaction as a general *multipolar interaction*. However, the latter case is unlikely to occur because all default pharmacophore rules for electrophiles comprehend heavy atoms bound to an electrophilic atom (see Section *Physicochemical property definitions*).

##### Ion-dipole interaction

Ion-dipole interactions are identified similarly to multipolar interactions (Figure S3), where $I$ is the ion centroid and D$=$Y is the dipole, having $D$ as the electrophile when $Y$ is the nucleophile and vice versa. Given the possible combinations of ions and dipoles, there are two favorable (*cation-nucleophile* and *anion-electrophile*) and two unfavorable interactions (*unfavorable anion-nucleophile* and *unfavorable cation-electrophile*).

Below we provide the default values extracted from literature[24] for each of the parameters presented in Figure S3.

- Default distance threshold: $d \leq 4.5$Å, where $d$ is calculated in relation to the nucleophilic/electrophilic atom ($D$) and the ion centroid ($I$);
- Default angle thresholds: $\theta \geq 60°$ and $\varphi \leq 40°$, where $\varphi$ (displacement angle) is the angle formed by the dipole normal ($\overrightarrow{N}$) and the vector connecting the ion centroid and the interacting electrophile/nucleophile.

Similar to dipole-dipole interactions, both planar and non-planar dipoles are accepted. Also, in cases where the atom $Y$ coordinate cannot be determined, only the distance between $D$ and $I$ is evaluated.

##### Ionic and repulsive interactions

Interactions involving ions are classified as either *ionic* or *repulsive* when the ions are oppositely or similarly charged, respectively. The only parameter evaluated in these interactions is the distance between the ion centroids whose upper-limit threshold is 6Å.[5,25–27]

##### Salt bridge

Since a salt bridge consists of a hydrogen bond and an ionic interaction co-occurring between the same interacting partners, salt bridges are identified by directly searching for pairs of hydrogen bonds and ionic interactions that match the mentioned requirement. However, as hydrogen bonds are modeled as an atom-atom interaction and ionic interactions as a group-group interaction, this requirement is fulfilled when the acceptor belongs to one ionic group and the donor to the other.

Salt bridges are not identified by default because the interactions required for its computation already account for its existence implicitly. However, if necessary, this interaction can be easily turned on through a flag in the method for calculating interactions.

##### Cation-$\pi$ interaction

Cation-$\pi$ interactions are identified when the cation and aromatic ring centroids are up to 6Å apart.[28]

##### Hydrophobic interaction

Hydrophobic interactions can be modeled as atom-atom interactions or surface-surface contacts, which is the default approach in our tool. The former is identified when any two hydrophobic atoms are up to 4.5Å apart.[5,29,30]

Surface-surface contacts are identified as follows. Firstly, the tool computes hydrophobic interactions between atoms as explained for the atom-atom model. Then, it identifies all hydrophobic atoms covalently bound to each other and merges them to form a hydrophobic cluster/island, called *hydrophobe*. Finally, the tool converts each atom-atom interaction to its hydrophobe-hydrophobe form by identifying the hydrophobic clusters comprehending each interacting atom, and attributing this interaction to them. However, it may be possible that not all hydrophobic atoms in a cluster participate in the surface-surface contact. For that reason, each interaction contains the hydrophobic cluster information as a whole and keeps track of which of their specific atoms are in contact.

##### Covalent interaction

Covalent bonds are automatically obtained from Open Babel or RDKit.

##### Atom overlap

Atom overlap identifies artifacts generated by low-resolution structures and homology models, consisting of unnatural overlap of two atoms. An overlap is defined as two atoms not covalently bound separated from each other by less than or equal to the sum of their covalent radii.[29]

##### Van der Waals clash

Van der Waals clashes are identified as in Pettersen et al.,[31] which describes a van der Waals clash by the following expression:

$$A_{vdw} + B_{vdw} - d \geq 0.6 \tag{3}$$

Where $d$ is the Euclidean distance between two atoms $A$ and $B$, $A_{vdw}$ and $B_{vdw}$ are their van der Waals radii, and 0.6 is the threshold for van der Waals clashes. Van der Waals radii are derived from Open Babel.

##### Van der Waals interaction

Van der Waals interactions are identified as in Jubb et al.,[29] which describes a van der Waals by the following expression:

$$d \leq A_{vdw} + B_{vdw} + 0.1 \tag{4}$$

Where $d$ is the Euclidean distance between two atoms $A$ and $B$, $A_{vdw}$ and $B_{vdw}$ are their van der Waals radii, and 0.1 is a margin of error. Van der Waals radii are derived from Open Babel.

##### Proximal interactions

Proximal interactions are defined as any two atoms separated by at least 2Å and at most 6Å.

##### Intramolecular interactions

Interactions involving atoms or atom groups from the same molecule are calculated using the specific interaction methods described in previous sections. In addition, there is an additional criterion for intramolecular interactions that consists of evaluating how many bonds separate the interacting atoms or atom groups. Note that two different molecules covalently bound to each other also fall into the rules discussed herein.

For van der Waals clashes, the interaction is only accepted if the atoms are separated by more than 4 bonds. This threshold was defined to avoid invalid clashes typically found in structures like the one shown in Figure S5.

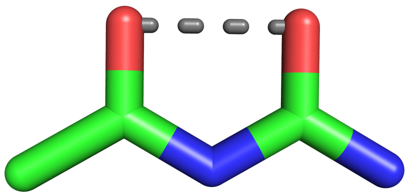

Figure S5: Example of an invalid van der Waals clash (gray dashed line) between two atoms separated by four bonds.

For the other interactions, the number of bonds separating two atoms must be higher than three;[31] otherwise, the interaction is ignored. The only exception is the hydrophobic interaction which is not considered for intramolecular interactions.

(18) Koebel, M. R.; Cooper, A.; Schmadeke, G.; Jeon, S.; Narayan, M.; Sirimulla, S. S$\cdots$O and S$\cdots$N Sulfur Bonding Interactions in Protein–Ligand Complexes: Empirical Consid-

erations and Scoring Function. *Journal of Chemical Information and Modeling* **2016**, *56*, 2298–2309.

(27) Mancini, A. L.; Higa, R. H.; Oliveira, A.; Dominiquini, F.; Kuser, P. R.; Yamagishi, M. E.; Togawa, R. C.; Neshich, G. STING Contacts: a web-based application for

identification and analysis of amino acid contacts within protein structure and across protein interfaces. *Bioinformatics* **2004**, *20*, 2145–2147.

(28) Gallivan, J. P.; Dougherty, D. A. Cation-$\pi$ interactions in structural biology. *Proceedings of the National Academy of Sciences* **1999**, *96*, 9459–9464.

(29) Jubb, H. C.; Higueruelo, A. P.; Ochoa-Montaño, B.; Pitt, W. R.; Ascher, D. B.; Blundell, T. L. Arpeggio: A Web Server for Calculating and Visualising Interatomic Interactions in Protein Structures. *Journal of Molecular Biology* **2017**, *429*, 365–371.

(30) Marcou, G.; Rognan, D. Optimizing Fragment and Scaffold Docking by Use of Molecular Interaction Fingerprints. *Journal of Chemical Information and Modeling* **2007**, *47*, 195–207.

(31) Pettersen, E. F.; Goddard, T. D.; Huang, C. C.; Couch, G. S.; Greenblatt, D. M.; Meng, E. C.; Ferrin, T. E. UCSF Chimera—A visualization system for exploratory research and analysis. *Journal of Computational Chemistry* **2004**, *25*, 1605–1612.
